# Haplotype genetic score analysis in 10,734 mother/infant pairs reveals complex maternal and fetal genetic effects underlying the associations between maternal phenotypes, birth outcomes and adult phenotypes

**DOI:** 10.1101/737106

**Authors:** Jing Chen, Jonas Bacelis, Pol Sole Navais, Amit Srivastava, Julius Juodakis, Amy Rouse, Mikko Hallman, Kari Teramo, Mads Melbye, Bjarke Feenstra, Rachel M. Freathy, George Davey-Smith, Deborah A. Lawlor, Jeffrey C. Murray, Scott M. Williams, Bo Jacobsson, Louis J. Muglia, Ge Zhang

**Affiliations:** Division of Biomedical Informatics, Cincinnati Children’s Hospital Medical Center, Department of Pediatrics, University of Cincinnati College of Medicine, Cincinnati, Ohio, United States of America; Department of Obstetrics and Gynecology, Institute of Clinical Sciences, Sahlgrenska Academy, University of Gothenburg, Gothenburg, Sweden; Region Västra Götaland, Sahlgrenska University Hospital, Department of Obstetrics and Gynecology, Gothenburg, Sweden; Department of Obstetrics and Gynecology, Institute of Clinical Sciences, Sahlgrenska Academy, University of Gothenburg, Gothenburg, Sweden; Division of Human Genetics, Center for Prevention of Preterm Birth, Perinatal Institute, Cincinnati Children’s Hospital Medical Center and March of Dimes Prematurity Research Center Ohio Collaborative, Department of Pediatrics, University of Cincinnati College of Medicine, Cincinnati, Ohio, United States of America; Center for Prevention of Preterm Birth, Perinatal Institute, Cincinnati Children’s Hospital Medical Center and March of Dimes Prematurity Research Center Ohio Collaborative, Department of Pediatrics, University of Cincinnati College of Medicine, Cincinnati, Ohio, United States of America; PEDEGO Research Unit and Medical Research Center Oulu, University of Oulu and Department of Children and Adolescents, Oulu University Hospital, Oulu, Finland; Obstetrics and Gynecology, University of Helsinki and Helsinki University Hospital, Helsinki, Finland; Department of Epidemiology Research, Statens Serum Institut, Copenhagen, Denmark; Department of Clinical Medicine, University of Copenhagen, Copenhagen, Denmark; Department of Medicine, Stanford University School of Medicine, Stanford, California, United States of America; Department of Epidemiology Research, Statens Serum Institut, Copenhagen, Denmark; Institute of Biomedical and Clinical Science, College of Medicine and Health, University of Exeter, Exeter, United Kingdom; MRC Integrative Epidemiology Unit at the University of Bristol, Bristol, United Kingdom; Population Health Science, Bristol Medical School, University of Bristol, Bristol, United Kingdom; and Bristol NIHR Biomedical Research Centre, United Kingdom; Department of Pediatrics, University of Iowa, Iowa City, Iowa, United States of America; Department of Population and Quantitative Health Sciences, Case Western Reserve University School of Medicine, Cleveland, OH, United States of America; Department of Obstetrics and Gynecology, Institute of Clinical Sciences, Sahlgrenska Academy, University of Gothenburg, Gothenburg, Sweden; Department of Genetics and Bioinformatics, Domain of Health Data and Digitalisation, Institute of Public Health, Oslo, Norway

## Abstract

Many maternal traits are associated with a neonate’s gestational duration, birth weight and birth length. These birth outcomes are subsequently associated with late onset health conditions. Based on 10,734 mother/infant duos of European ancestry, we constructed haplotype genetic scores to dissect the maternal and fetal genetic effects underlying these observed associations. We showed that maternal height and fetal growth jointly affect the duration of gestation – maternal height positively influences the gestational duration, while faster fetal growth reduces gestational duration. Fetal growth is influenced by both maternal and fetal effects and can reciprocally influence maternal phenotypes: tall maternal stature and higher blood glucose causally increase birth size; in the fetus, the height and metabolic risk increasing alleles can lead to increased and decreased birth size respectively; birth weight-raising alleles in fetus may reduce gestational duration and increase maternal blood pressure. These maternal and fetal genetic effects can largely explain the observed associations between the studied maternal phenotypes and birth outcomes as well as the life-course associations between these birth outcomes and adult phenotypes.

## INTRODUCTION

Epidemiological studies have demonstrated that maternal physical and physiological traits associate with birth outcomes. For example, maternal height is positively associated with gestational duration(1, 2), birth weight and birth length(3, 4); higher maternal blood glucose is associated with higher birth weight(5); and elevated maternal blood pressure is associated with reduced birth weight(6, 7). These birth outcomes in turn associate with many long-term adverse health outcomes in the offspring, such as obesity(8), type 2 diabetes(9), hypertension(10), and cardiovascular diseases(11, 12). Different mechanisms have been proposed to explain the observed associations between maternal phenotypes and pregnancy outcomes(13–16) (Figure 1A) as well as the life-course associations between birth outcomes and adult phenotypes(17–20) (Figure 1B). Briefly, these include various causal effects (e.g. maternal effects(21)); genetically confounded associations due to genetic sharing (between mothers and infants) or shared genetic effects (between a birth outcome and an adult phenotype)(20); and confounding due to external or intra-uterine environment. Fetal phenotypes can also affect maternal physiology during or even after pregnancy (fetal drive)(22). Dissecting these different underlying mechanisms would increase knowledge of the etiology of these critical pregnancy outcomes and provide insights into how pregnancy outcomes are linked with later onset disorders(13, 23). Understanding the causal effects of modifiable maternal phenotypes could have implications for clinical interventions to prevent adverse birth outcomes(24). The shared genetic causes between pregnancy characteristics and late offspring outcomes could provide insights into the molecular pathways through which these shared genetic effects are mediated(20).

**Figure 1.**
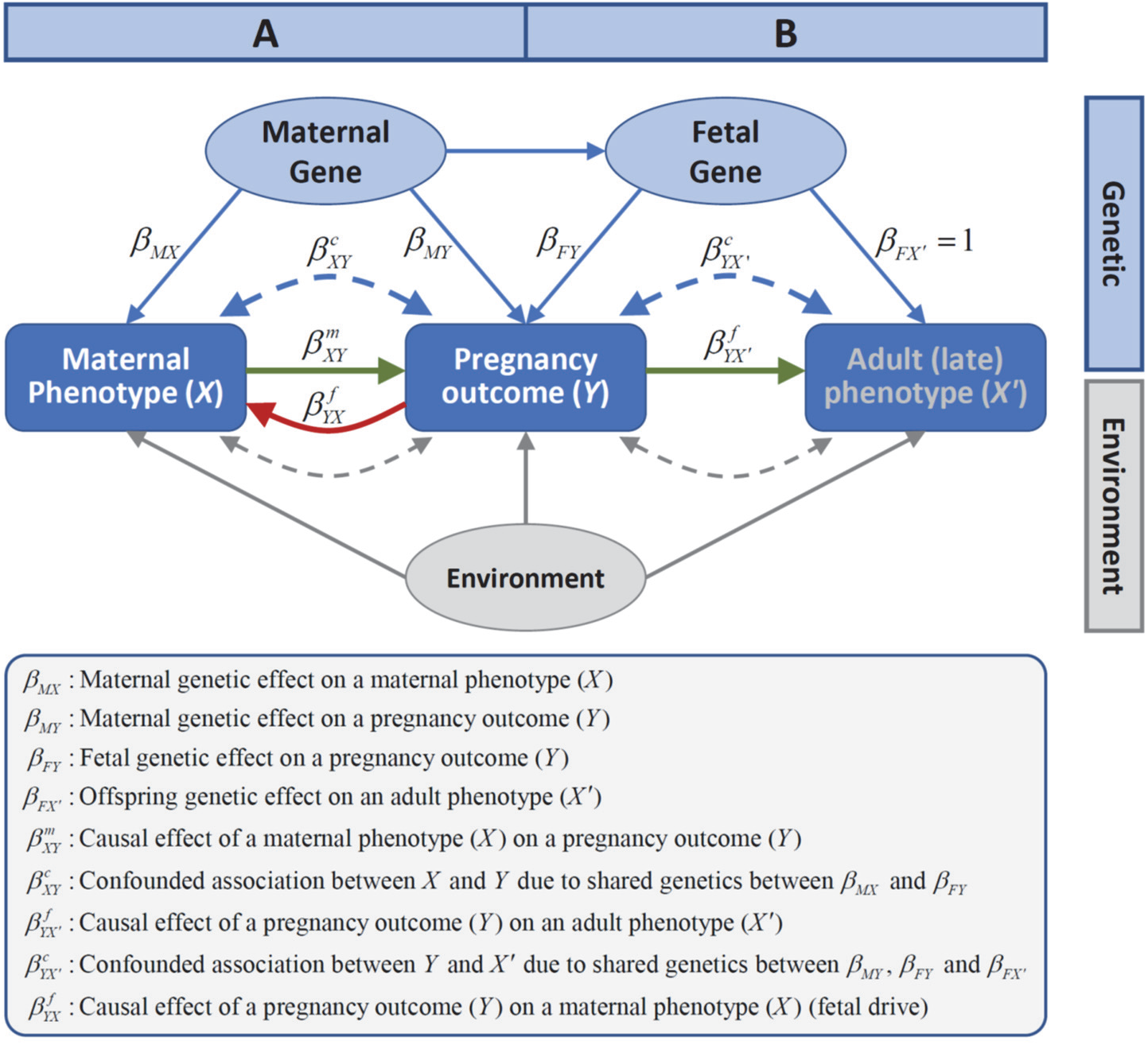
The different mechanisms underlying (A) the associations between maternal phenotypes and pregnancy outcomes and (B) the associations between pregnancy outcomes and late adult phenotypes in offspring. These mechanisms include: 1) causal effects of maternal phenotypes on pregnancy outcomes 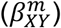 and causal effects of pregnancy outcomes on adult phenotypes 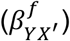 (green arrows); 2) genetically confounded associations between maternal phenotypes and pregnancy outcomes 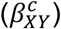 due to genetic sharing between mothers and infants, and genetically confounded associations between birth outcomes and adult phenotypes in offspring 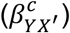 due to shared genetic effects (blue dashed arrows); 3) confounding due to environmental effects (grey dashed arrows, which were not examined in this study) and 4) Fetal drive 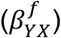 – fetus causally influence maternal phenotypes during pregnancy (red arrow).

Mendelian randomization (MR)(25) studies utilizing maternal genotypes as instrumental variables have been used to probe the causal relationships between maternal phenotypes and pregnancy outcomes(13, 16, 26). Using this approach, Tyrrell et al.(24) demonstrated that higher maternal BMI and blood glucose levels are causally associated with higher birth weight, whilst higher maternal systolic blood pressure causes lower birth weight. Using a genome-wide association (GWA) approach, Horikoshi et al.(20) demonstrated strong inverse genetic correlations between birth weight and adult cardiometabolic diseases, suggesting a strong genetic component underlying the observed associations between low birth weight and cardiometabolic risks. More recently, Warrington et al. developed a structural equation modelling (SEM) method(27) and a weighted linear model (WLM) approximation to separate maternal from fetal effects. Using this method, they estimated maternal and fetal genetic effects on birth weight genome-wide, and investigated associations between those genetic effects on birth weight and adult systolic blood pressure(28). We previously developed a novel MR method that utilizes non-transmitted maternal alleles as a valid genetic instrument for maternal phenotypic effects on fetal/offspring outcomes(15). We showed that the observed association between maternal height and fetal size is mainly due to shared genetics, while the association between maternal height and gestational duration is more likely causal. Using this approach, Horikoshi et al.(20) showed that alleles that increase blood glucose increase birth weight by maternal effect and that type 2 diabetes risk alleles could influence birth weight through both maternal and fetal genetic effects albeit acting in opposite directions. These studies have already provided novel understandings about the causal relationships between many maternal phenotypes and birth outcomes. They have also highlighted genetic contributions to life-course associations between birth weight and late onset diseases(20, 28). However, previous studies have usually examined causal effects of maternal phenotypes on either birth weight or gestational duration separately despite the strong association between them(29, 30). The studies focusing on birth weight have not explored whether any effects on birth weight are driven by effects on gestational duration and have not investigated the effects of fetal growth on gestational duration and maternal phenotypes during pregnancy.

To further our understanding of how various maternal phenotypes are correlated with pregnancy outcomes through maternal or fetal genetic effects, and how fetal growth influences gestational duration and maternal physiological changes during pregnancy, we expanded our haplotype-based method by considering the mother/fetus duo (pregnancy) as the analytical unit(31) and explicitly modeled maternal and fetal genetic effects using haplotype genetic scores (Figure 2). By testing associations between these haplotype genetic scores and birth outcomes, we systematically investigated the maternal and fetal genetic effects underlying the observed associations between four maternal traits (height, BMI, blood pressure, and blood glucose levels) and pregnancy outcomes (gestational duration, birth weight and birth length). Using this approach, we also examined the effects of fetal growth (using gestational age adjusted birth weight as a measure of this) on pregnancy outcomes, maternal blood pressure and blood glucose levels measured during pregnancy.

**Figure 2.**
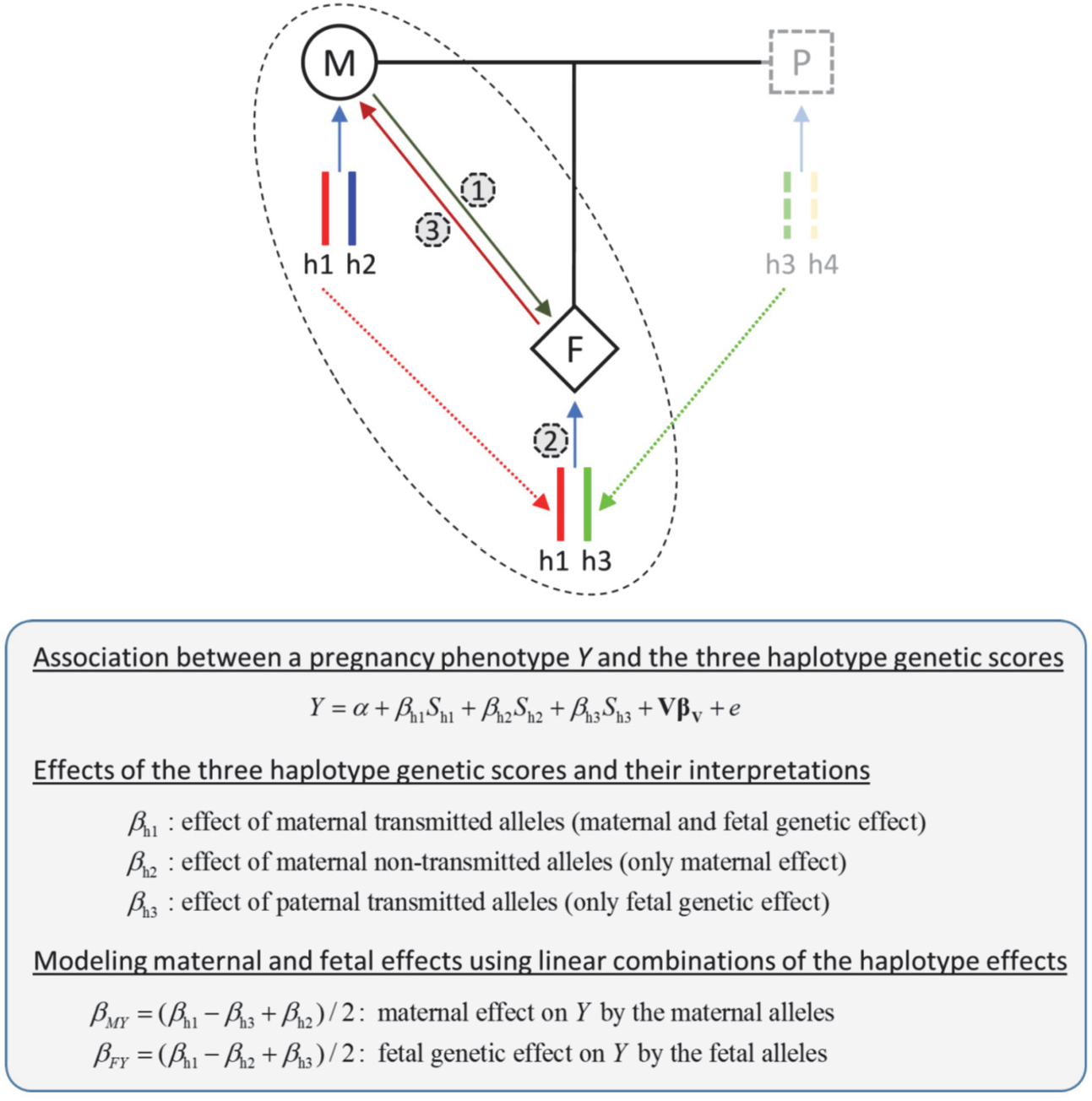
Genetic dissection of maternal and fetal genetic effects using haplotype genetic scores in mother/child pairs. There are three groups of alleles (haplotypes) in a mother (M) / fetus (F) duo: the maternal transmitted alleles (h1) can affect a pregnancy outcome through a maternal (1) or/and fetal genetic effect (2); the maternal non-transmitted alleles (h2) can only affect a pregnancy outcome through maternal effect (1) and the paternal transmitted alleles (h3) only through fetal effect (2) (assuming no paternal effect). The paternal transmitted alleles (h3) could influence a maternal phenotype during pregnancy by fetal drive (3).

## METHODS

### Data sets

We used phenotype and genome-wide single nucleotide polymorphism (SNP) data of 10,734 mother/infant pairs from six birth studies (Supplementary Table 1). These include three case/control data sets collected from Nordic countries (FIN, MOBA and DNBC) for genetic studies of preterm birth(32), a longitudinal birth cohort (ALSPAC)(33) from the UK, and two studies of pregnancy outcomes (HAPO(5) and the GPN(34)) from the US. A detailed description of these data sets can be found in the Supplementary methods.

We focused on investigating the relationships between maternal height, pre-pregnancy BMI, blood pressure, blood glucose levels and pregnancy outcomes including gestational duration (as both quantitative and dichotomous preterm/term trait), birth weight and birth length. Maternal height, pre-pregnancy BMI and the three pregnancy outcomes were available in most of the studies (birth weight and length were not available in the MOBA data used here, and birth length was not available in the DNBC data used here) (Supplementary Table 2). Maternal blood pressure during pregnancy was available in ALSPAC and HAPO. In the HAPO data, blood pressure was measured between 24 and 32 weeks of pregnancy when the mothers underwent an oral glucose tolerance test (OGTT)(5). In ALSPAC, all blood pressure measurements undertaken during antenatal care were extracted from clinical records; women had a median of 13 (interquartile range 11-16) blood pressure measurements(35). We used the values measured between 30 to 36 weeks of pregnancy (as close as possible to when the blood pressure was measured in HAPO). Maternal fasting plasma glucose (FPG) levels during pregnancy were available only in the HAPO study. FPG was measured in over 4000 ALSPAC mothers in a follow-up data collection 18 years after the pregnancy (Supplementary methods and Supplementary Table 2).

As gestational duration is a key determinant of birth weight and birth size, we only included pregnancies with spontaneous deliveries and excluded mother/child pairs without gestational duration information. Pregnancies with known gestational or fetal complications and pre-existing medical conditions were excluded. Detailed inclusion/exclusion criteria are provided in Supplementary methods.

### Genotype data

Genome-wide SNP data were generated using either Affymetrix 6.0 or various Illumina genotyping arrays. Standardized genotype quality control procedures were applied to all data sets. Participants of non-European ancestry were identified and excluded using principal components analysis (PCA) (Supplementary methods and Supplementary Figure 1). Genome-wide imputation was performed using Minimac3(36) and the reference haplotypes from phase 3 of the 1000 Genomes Project(37). In each data set, the haplotype phasing was done in all maternal and fetal samples using Shapeit2(38). This program accommodates mother/child relationship and accurately estimates mother-child allele transmission when phasing mother/child duos together.

### Construction of Genetic scores

We constructed weighted genetic scores to instrument various maternal phenotypes using genome wide association (GWA) SNPs and their estimated effect sizes reported by the most recent large GWA studies (Supplementary methods and Supplementary Table 4). Specifically, 2,130 height associated SNPs and 628 BMI associated SNPs reported by the GIANT consortium(39) were used to build genetic scores for height and BMI, respectively. 831 SNPs associated with blood pressure(40) were used to build genetic scores for blood pressure. For FPG, we used 22 SNPs associated with FPG levels identified in non-diabetic individuals(41). We also constructed a type 2 diabetes (T2D) genetic score using 306 T2D SNPs (excluding SNPs overlapping or in close linkage disequilibrium with the 22 FPG SNPs) (Supplementary methods and Table S3). To examine the effect of fetal growth (as proxied by birth weight) on pregnancy outcomes and maternal blood pressure and FPG during pregnancy, we constructed genetic scores using 86 SNPs associated with birth weight with confirmed fetal effect(42).

For each set of GWA SNPs, we constructed two genotype genetic scores: *S*_mat_ (maternal genotype score), *S*_fet_ (fetal genotype score); and three haplotype genetic scores: *S*_h1_, *S*_h2_ and *S*_h3_ respectively based on the maternal transmitted (h1), maternal non-transmitted (h2) and paternal transmitted alleles (h3) (Figure 2). It follows that, *S*_mat_ = *S*_h1_ + *S*_h2_ and *S*_fet_ = *S*_h1_ + *S*_h3._

### Statistical analyses

#### Phenotype associations and instrumental strength of genetic scores

We first assessed the associations between the four maternal phenotypes (*X*) (i.e. height, BMI, blood pressure and FPG) and each pregnancy outcome (*Y*) (i.e., gestational duration, preterm birth, birth weight and birth length) using regression analyses. Maternal age, fetal sex, maternal height, pre-pregnancy BMI were included as covariates. As gestational duration influences birth weight and length in a non-linear fashion, the first three orthogonal polynomials of gestational duration were included as covariates in the analysis of birth weight and length. These analytical models are described in more detail in the Supplementary methods.

The instrumental strength of the genetic scores was checked by the variance in a maternal phenotype explained (*R*^2^) by the corresponding genetic scores.

#### Association tests between haplotype genetic scores and pregnancy outcomes

Associations between the haplotype genetic scores and the pregnancy outcomes were tested using regression models like those used in the association analysis described above, except that the maternal phenotypes (*X*) were replaced by their corresponding three haplotype genetic scores (*S*_h1_ + *S*_h2_ + *S*_h3_). The associations between these haplotype scores can differentiate between maternal and fetal genetic effects (Figure 2). Specifically, an association of *S*_h2_ (maternal non-transmitted haplotype score) with a pregnancy outcome suggests a maternal (intrauterine phenotypic) effect, whereas an association of *S*_h3_ (paternal transmitted haplotype score) with the pregnancy outcomes suggests fetal genetic effects. As the three haplotype genetic scores were simultaneously tested in a single regression model, the effect size estimates of these haplotypes scores and their associated p-values can be directly compared to assess the directions and relative contributions of the maternal and fetal effects.

#### Modeling of maternal and fetal genetic effects

While *S*_h2_ and *S*_h3_ can be used to draw inference about maternal and fetal genetic effects, this question can be addressed with greater statistical power by also including *S*_h1_, the maternal transmitted haplotype score, in the model. Thus, we modeled the maternal effect and fetal genetic effect as different linear combinations of the regression coefficients of the three haplotype genetic scores (Figure 2 and Supplementary methods)(31). Under the assumptions of additivity between maternal and fetal effect and zero parent-of-origin effect, the total effect (*β*_h1_) of the maternal transmitted haplotype (h1) should be equal to the summation of the maternal effect (*β*_h2_) of the non-transmitted haplotype (h2) and fetal genetic effect (*β*_h3_) of the paternal transmitted haplotype (h3). Thus, (*β*_h1_ - *β*_h3_) and (*β*_h1_ - *β*_h2_) respectively represent the maternal effect and the fetal genetic effect of the maternal transmitted haplotype (h1). Therefore, the average maternal effect (*β*_*MY*_) of the two maternal haplotypes on a birth outcome (*Y*) can be expressed as (*β*_h1_ - *β*_h3_ + *β*_h2_)/2and the average fetal genetic effect (*β*_*FY*_) of the maternal and paternal transmitted haplotypes can be expressed as (*β*_h1_ - *β*_h2_ + *β*_h3_)/2. Since these linear combinations also capture the maternal or fetal genetic effects of the maternal transmitted haplotype (h1), they are more powerful than the methods only using the maternal non-transmitted haplotype (h2) or the paternal transmitted haplotype (h3) as instruments respectively for maternal effect and fetal genetic effect.

#### Estimation of maternal causal effects

The estimated maternal effect 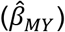 from the haplotype genetic score association analyses can be interpreted as the amount of change in a pregnancy outcome (*Y*) caused by certain amount of difference in a maternal phenotype (*X*) associated with one-unit genetic score. The maternal causal effect 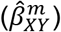 was estimated using the ratio estimate(43) 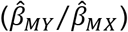, where 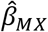 is the estimated maternal effect on the maternal phenotype (Figure 2). As an alternative, we also performed instrumental variable analysis using the two-stage least squares (TSLS) approach(43), with the maternal non-transmitted haplotype score (*S*_h2_) as the genetic instrument for maternal causal effect(15).

#### Estimation of genetically confounded associations

The fetal genetic effect (*β*_*FY*_) reflects that the genetic variants associated with an adult phenotype (*X′*) in the offspring or the corresponding maternal phenotype (*X*) have direct fetal genetic effect on a pregnancy outcome (*Y*). This shared genetic effect can confound the association between a maternal phenotype (*X*) and a pregnancy outcome (*Y*) as well as the association between the pregnancy outcome (*Y*) and the adult phenotype (*X′*) in offspring (Figure 1).

By assuming that all the genetic variants associated with an adult phenotype in offspring (*X′*) or the corresponding maternal phenotype (*X*) have similar effect on a pregnancy outcome *Y* as the fetal genetic effect estimated from the genetic score built on known GWA SNPs 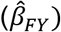, we can approximately estimate the magnitude of these genetically confounded associations (see Supplementary methods for details). Specifically, the genetically confounded association between a maternal phenotype (*X*) and a pregnancy outcome (*Y*) due to the shared genetic effect can be estimated by:

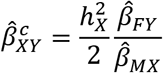

where 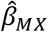 is estimated maternal effect and 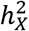 is the heritability (the proportion of additive genetic variance) of the maternal phenotype (*X*).

Similarly, the genetically confounded association between a pregnancy outcome (*Y*) and an adult (late) phenotype (*X′*) in offspring can be estimated by:

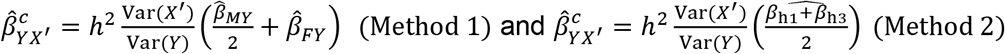

where *h*^2^ is the heritability of the adult phenotype, Var(*X′*) and Var(*Y*) are respectively the variance of the adult phenotype and the variance of the pregnancy outcome. The first method (Method 1) can partition the confounded association into the maternal 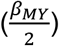 and the fetal component (*β*_*FY*_) (Supplementarly methods).

### Multivariable MR analysis

The genetic scores built on hundreds or thousands of SNPs are likely to be less specific, as they are more likely to be associated with other phenotypic traits(44), which can introduce ambiguities in the interpretation of genetic score associations(45). To circumvent this issue, we performed a two-sample multivariable MR analysis(46, 47) using the MR pleiotropy residual sum and outlier (MR-PRESSO)(48) to detect and correct for variants with horizontal pleiotropic effects(49) in multiple variants MR testing. This analysis studies the maternal or fetal genetic effects by testing whether the effects of the maternal transmitted (h1), maternal non-transmitted (h2) and paternal transmitted (h3) alleles of the GWA SNPs on a pregnancy outcome are proportional to their reported effects on an adult phenotype (*X′*) in the reference GWA studies (Supplementary methods and Figure S2). The allele-specific effect estimates (for the h1, h2 and h3 alleles) of each SNP on a pregnancy outcome were obtained using the same regression methods for haplotype genetic score analysis (Supplementary methods).

We did meta-analyses of the results from all available data sets to generate the overall results. Fixed-effect meta-analysis was used to combine the regression coefficients and standard errors from individual study and we check between study heterogeneity using Cochran’s Q test.

## RESULTS

### Phenotypic associations between maternal phenotypes and pregnancy outcomes

We used 10,734 mother/infant pairs with both genotype and phenotype data in our analyses (Supplementary Table 1). We only included spontaneous deliveries and excluded mother/child pairs without gestational duration information. Distributions of key variables for the maternal phenotypes and pregnancy outcomes are shown in Supplementary Table 2 and Supplementary Figures 5 – 7.

The meta-analysis across the six data sets showed that taller maternal height was associated with longer gestational duration (0.14 day/cm, 95% CI: 0.10-0.18), lower preterm birth risk (OR=0.97 /cm, 95% CI: 0.96-0.98), higher birth weight (15 g/cm, 95% CI: 13.7-16.3) and length (0.068 cm/cm, 95% CI:0.061-0.075). Maternal BMI was positively associated with birth weight (16 kg/m^2^, 95% CI: 13.8-18.2) and birth length (0.047 kg/m^2^, 95 % CI: 0.046-0.048) but was not associated with gestational duration or preterm birth risk (Supplementary Table 3).

Using data from ALSPAC and HAPO, we observed that maternal blood pressure during pregnancy was negatively associated with gestational duration and birth weight. In HAPO, there was a strong positive association between maternal fasting plasma glucose (FPG) and birth weight (192 g/(mmol/L), 95% CI: 116-268) and birth length (0.62 cm/(mmol/L), 95% CI: 0.27- 0.97). However, the association between FPG measured 18 years after pregnancy with either birth weight or length in the ALSPAC data set was close to zero with wide confidence intervals (Supplementary Table 3).

### Associations between genetic scores and maternal phenotypes

We examined the instrumental strength of the genetic scores for the various maternal phenotypes (Supplementary Table 4). The maternal genotype genetic scores (*S*_mat_) were associated with the corresponding maternal phenotypes and explained a substantial fraction of the phenotypic variances with similar contributions from the transmitted (h1) or the non-transmitted haplotype scores (h2) (Supplementary Table 5).

The maternal height genotype score (*S*_mat_) explained >20% of the maternal height variance (Supplementary Table 6) and the maternal BMI genotype score (*S*_mat_) ~5% of the maternal BMI variance (Supplementary Table 7). The blood pressure genotype scores explained over 2% variance in maternal blood pressure (Supplementary Table 8), which is less than half of the reported fraction of variance explained by the same score (5.7%) in published GWAS of non-pregnant women and men (generally of an older age than pregnant women)(40), suggesting that these BP SNPs have a larger effect on blood pressure in older populations, or a weaker effect on maternal blood pressure during pregnancy.

In HAPO, the maternal FPG genetic score built on 22 SNPs explained 8.3% of the FPG variance measured between 24~32 weeks. In ALSPAC, the same score explained 4.1% of the variance of FPG measured 18 years after pregnancy. By contrast, the T2D score (306 SNPs) explained much less FPG variance (Supplementary Table 9).

For each phenotype, we checked the correlations among the various genotype and haplotype genetic scores (Supplementary Table 10). For height, we observed significant correlations between the maternal genotype (*S*_mat_) and the paternal transmitted haplotype score (*S*_h3_) and between the maternal transmitted (*S*_h1_) and non-transmitted scores (*S*_h2_), indicating assortative mating(15, 50, 51) and increased homozygosity of height associated SNPs.

### Maternal causal effects and genetically confounded associations between maternal phenotypes and birth outcomes

We next utilized haplotype genetic scores as genetic instruments to dissect the maternal and fetal genetic effects underlying the observed associations between maternal phenotypes and pregnancy outcomes (Table 1). Detailed meta-analysis of individual data sets can be found in Supplementary Figures 8 – 12. Based on the estimated maternal and fetal genetic effects, we estimated the maternal causal effects and genetically confounded associations between maternal phenotypes and birth outcomes due to shared genetics (Methods).

**Table 1.** Association between haplotype genetic scores and birth outcomes.

| Maternal trait (unit)<br>Haplotype Score Tests<sup>a</sup> | Gestational days |  |  | Preterm birth (log(OR)) |  |  | Birth weight (g) |  |  | Birth length (cm) |  |  |
| --- | --- | --- | --- | --- | --- | --- | --- | --- | --- | --- | --- | --- |
|  | beta | se | p-val | beta | se | p-val | beta | se | p-val | beta | se | p-val |
| **Height (cm)** |  |  |  |  |  |  |  |  |  |  |  |  |
| Maternal trans ($\beta_{h1}$) | 0.038 | 0.046 | 0.41 | -0.027 | 0.011 | **0.016** | 20 | 1.6 | **6.10E-38** | 0.097 | 0.0087 | **1.10E-28** |
| Maternal non-trans ($\beta_{h2}$) | 0.17 | 0.047 | **0.00022** | -0.037 | 0.011 | **0.00097** | 6.3 | 1.6 | **7.60E-05** | 0.022 | 0.0088 | **0.011** |
| Paternal trans ($\beta_{h3}$) | -0.041 | 0.046 | 0.37 | 0.023 | 0.011 | **0.038** | 14 | 1.6 | **3.10E-19** | 0.074 | 0.0085 | **4.70E-18** |
| Maternal effect ($\beta_{MY}$) | 0.13 | 0.04 | **0.0017** | -0.044 | 0.0098 | **9.00E-06** | 6.3 | 1.4 | **5.20E-06** | 0.022 | 0.0076 | **0.0033** |
| Fetal effect ($\beta_{FY}$) | -0.089 | 0.041 | **0.029** | 0.016 | 0.0099 | 0.1 | 14 | 1.4 | **1.10E-23** | 0.074 | 0.0076 | **2.70E-22** |
| **BMI (kg/m<sup>2</sup>)** |  |  |  |  |  |  |  |  |  |  |  |  |
| Maternal trans ($\beta_{h1}$) | -0.059 | 0.17 | 0.73 | -0.065 | 0.041 | 0.11 | 23 | 5.8 | **7.00E-05** | 0.066 | 0.032 | **0.038** |
| Maternal non-trans ($\beta_{h2}$) | -0.11 | 0.17 | 0.52 | 0.015 | 0.041 | 0.71 | 8.8 | 5.8 | 0.13 | 0.079 | 0.032 | **0.013** |
| Paternal trans ($\beta_{h3}$) | 0.027 | 0.17 | 0.88 | 0.049 | 0.04 | 0.23 | -6.5 | 5.7 | 0.26 | 0.027 | 0.032 | 0.39 |
| Maternal effect ($\beta_{MY}$) | -0.098 | 0.15 | 0.51 | -0.048 | 0.036 | 0.18 | 19 | 5 | **0.00016** | 0.06 | 0.028 | **0.031** |
| Fetal effect ($\beta_{FY}$) | 0.043 | 0.15 | 0.77 | -0.018 | 0.036 | 0.6 | 4 | 5 | 0.43 | 0.0077 | 0.028 | 0.78 |
| **BP<sup>b</sup> (mmHg)** |  |  |  |  |  |  |  |  |  |  |  |  |
| Maternal trans ($\beta_{h1}$) | -0.22 | 0.064 | **0.00067** | 0.034 | 0.015 | **0.027** | -6.8 | 2.2 | **0.0016** | -0.018 | 0.012 | 0.13 |
| Maternal non-trans ($\beta_{h2}$) | -0.035 | 0.064 | 0.59 | 0.047 | 0.016 | **0.0023** | -3.1 | 2.2 | 0.16 | -0.018 | 0.012 | 0.13 |
| Paternal trans ($\beta_{h3}$) | -0.016 | 0.064 | 0.8 | 0.005 | 0.015 | 0.74 | -5.9 | 2.1 | **0.0053** | -0.017 | 0.012 | 0.15 |
| Maternal effect ($\beta_{MY}$) | -0.12 | 0.055 | **0.033** | 0.038 | 0.013 | **0.0045** | -2 | 1.9 | 0.27 | -0.0096 | 0.01 | 0.35 |
| Fetal effect ($\beta_{FY}$) | -0.1 | 0.056 | 0.075 | -0.0035 | 0.013 | 0.8 | -4.8 | 1.9 | **0.0094** | -0.0086 | 0.01 | 0.4 |
| **FPG (mmol/L)** |  |  |  |  |  |  |  |  |  |  |  |  |
| Maternal trans ($\beta_{h1}$) | -3.9 | 1.8 | **0.029** | 0.6 | 0.43 | 0.16 | 13 | 59 | 0.82 | -0.067 | 0.32 | 0.84 |
| Maternal non-trans ($\beta_{h2}$) | -3.2 | 1.8 | 0.071 | 0.54 | 0.43 | 0.21 | 270 | 59 | **4.70E-06** | 0.71 | 0.32 | **0.029** |
| Paternal trans ($\beta_{h3}$) | 2.7 | 1.7 | 0.12 | -0.38 | 0.43 | 0.37 | -52 | 59 | 0.38 | -0.13 | 0.32 | 0.69 |
| Maternal effect ($\beta_{MY}$) | -5 | 1.5 | **0.0012** | 0.77 | 0.37 | **0.039** | 170 | 51 | **0.0011** | 0.37 | 0.28 | 0.19 |
| Fetal effect ($\beta_{FY}$) | 0.99 | 1.5 | 0.51 | -0.14 | 0.36 | 0.7 | -150 | 50 | **0.0022** | -0.46 | 0.28 | 0.096 |
| **T2D (log(OR))** |  |  |  |  |  |  |  |  |  |  |  |  |
| Maternal trans ($\beta_{h1}$) | 0.013 | 0.3 | 0.97 | -0.0079 | 0.074 | 0.91 | -14 | 10 | 0.17 | -0.065 | 0.056 | 0.25 |
| Maternal non-trans ($\beta_{h2}$) | 0.05 | 0.31 | 0.87 | 0.013 | 0.072 | 0.86 | 33 | 10 | **0.0012** | 0.059 | 0.057 | 0.3 |
| Paternal trans ($\beta_{h3}$) | 0.83 | 0.31 | **0.0069** | -0.11 | 0.073 | 0.13 | -28 | 10 | **0.0061** | -0.033 | 0.057 | 0.56 |
| Maternal effect ($\beta_{MY}$) | -0.39 | 0.27 | 0.15 | 0.058 | 0.063 | 0.36 | 24 | 8.9 | **0.0064** | 0.015 | 0.049 | 0.76 |
| Fetal effect ($\beta_{FY}$) | 0.4 | 0.27 | 0.14 | -0.067 | 0.062 | 0.28 | -39 | 8.8 | **1.30E-05** | -0.085 | 0.049 | 0.082 |

a: $\beta_{h1}$ , $\beta_{h2}$ and $\beta_{h3}$ are the effects of the three haplotype genetic scores. $\beta_{MY} = (\beta_{h1} - \beta_{h3} + \beta_{h2})/2$ and $\beta_{FY} = (\beta_{h1} - \beta_{h2} + \beta_{h3})/2$ are respectively the maternal and fetal genetic effects modeled by linear combinations of the haplotype effects.
b: BP: mean of the SBP and DBP scores

#### Maternal height

The maternal non-transmitted height genetic score (*S*_h2_) was positively associated with gestational duration (P=0.00022) and negatively associated with preterm birth (P=0.00097) (Table 1). The ratio estimates showed a maternal causal effect of ~1.0 days (95% CI:0.38 to 1.64) longer gestation per 1SD (6.4cm) increase in maternal height. This effect was offset by a weaker and opposite fetal genetic effect of 0.71 days (95% CI:0.07 to 1.35) shorter gestation per the same amount of genetic score associated with 1SD increase in maternal height (Figure 3 and Supplementary Table 11).

**Figure 3.**
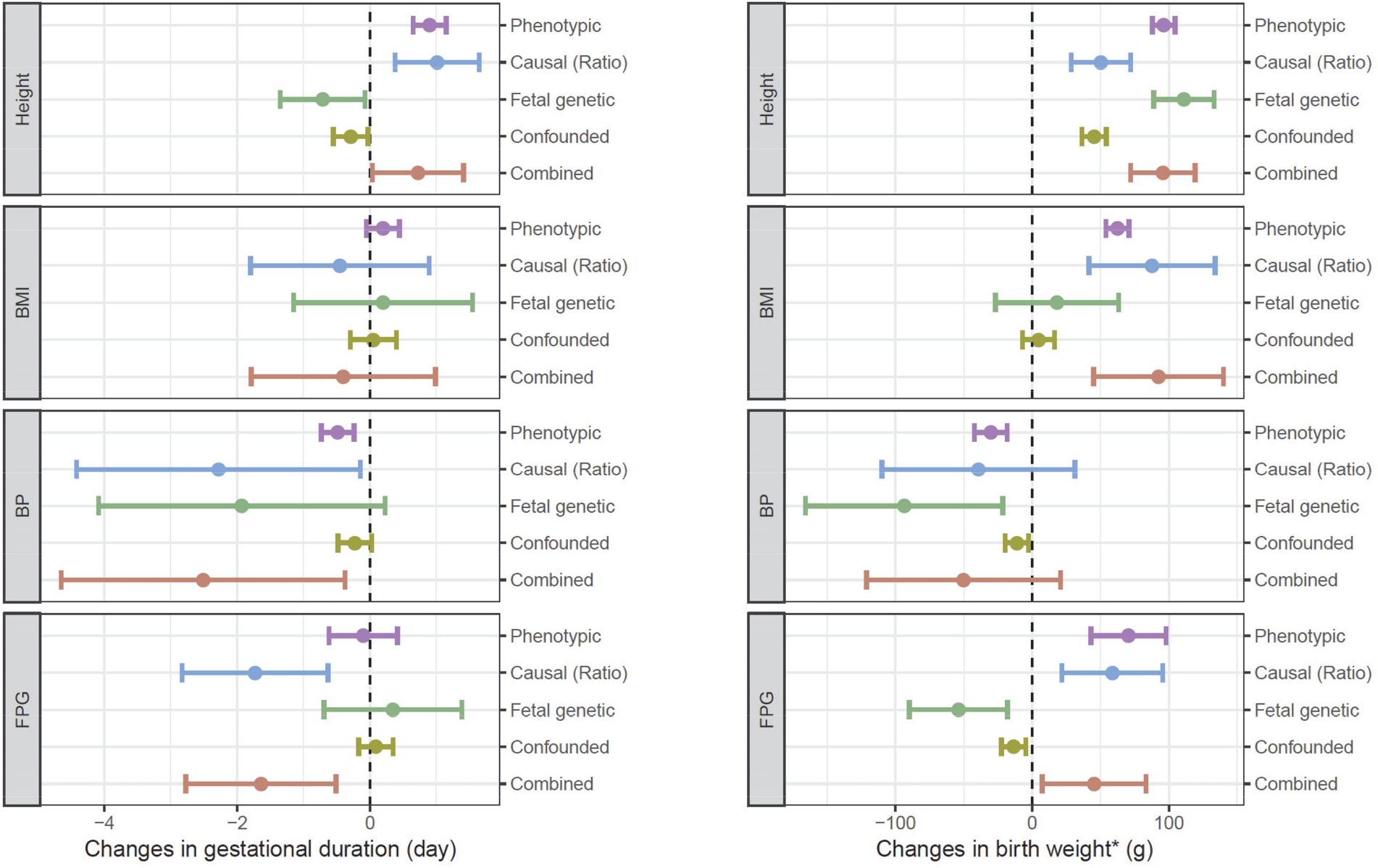
Observed phenotypic associations, estimated maternal causal effects, fetal genetic effects and genetically confounded associations per 1SD change in maternal traits on gestational duration (left) and birth weight (right). The 1SD for maternal traits are: 6.4cm (height), 4.0kg/m^2^ (BMI), 5.8mmHg (BP) and 0.36mmol/L (FPG).

Maternal and paternal transmitted haplotype scores (*S*_hi_ and *S*_h3_) for height were positively associated with birth weight and birth length. The maternal non-transmitted score (*S*_h2_) was also positively associated with birth weight and length but the effect estimates were smaller than the transmitted haplotype scores (Table 1). The larger effects of transmitted alleles indicate height associated SNPs can influence growth in early prenatal development through fetal genetic effect. The estimates for maternal causal and fetal genetic effects were respectively 50 g (95% CI: 29 to 72) and 111 g (95% CI: 89 to 133) for birth weight and 0.18 cm (95% CI: 0.06 to 0.30) and 0.59 cm (95% CI: 0.50 to 0.71) per genetic alleles associated with 1SD (6.4cm) increase in maternal height (Figure 3 and Supplementary Table 11).

#### Maternal pre-pregnancy BMI

BMI haplotype genetic scores were not significantly associated with gestational duration or preterm birth, suggesting minimal maternal and fetal effect of the BMI associated SNPs on gestational duration. Linear hypotheses modeling suggested that BMI associated SNPs have some maternal and no fetal effect on birth weight and length (Table 1). The estimated maternal causal effect on birth weight was 88 g (95% CI: 42 to 134) and on birth length was 0.28 cm (95% CI: 0.02 to 0.53) per 1SD (4.0kg/m^2^) genetically increased BMI (Figure 3 and Supplementary Table 11).

#### Maternal blood pressure

There was a significant association between the maternal non-transmitted blood pressure genetic score and increased preterm birth risk (P = 0.0023) suggesting the association between maternal blood pressure and gestational duration was primarily driven by a maternal effect (Table 1). The estimated causal effect sizes based on the ratio method were −2.3 days (95% CI: −4.4 to −0.09) on gestational duration and OR=2.1 (95% CI: 1.2 to 3.5) in preterm birth risk per 1SD (6mmHg) genetically increased maternal blood pressure (Figure 3 and Supplementary Table 11).

Both maternal transmitted and paternal transmitted blood pressure scores were negatively associated with birth weight (P=0.0016 and 0.0053 for *S*_h1_ and *S*_h3_ respectively, Table 1), suggesting the negative association between maternal blood pressure and birth weight was mainly caused by a fetal genetic effect. The estimated effect size was a reduction of 94 g (95% CI: 21 to 166) in birth weight by alleles associated with 1SD (6mmHg) increase in maternal blood pressure (Figure 3 and Supplementary Table 11). This finding is consistent with Horikoshi et al.(20), which showed negative genetic correlation between birth weight and blood pressure.

#### Maternal fasting plasma glucose

We observed a positive association between maternal non-transmitted FPG genetic score (*S*_h2_) and birth weight (P=4.7E-6), indicating a strong causal effect of increased maternal FPG level on higher birth weight (Table 1). In the HAPO data set, the estimated causal effect size by TSLS based on (*S*_h2_) was 147 g (95% CI: 15 to 279) per 1SD (0.36 mmol/L) increase in maternal FPG and the ratio estimate in all mother/child pairs was 59 g (95% CI: 22 to 96) (Supplementary Table 11). Interestingly, but not unexpectedly, the linear modeling showed a negative fetal genetic effect of FPG increasing alleles on birth weight (−54 g, 95% CI: −90 to −18). This result is in line with the previous epidemiological finding that paternal diabetes is associated with lower birth weight(52). The linear hypothesis modeling also showed a significant negative maternal effect of high maternal FPG on gestational duration (−1.73 days, 95% CI: −2.83 to −0.63) and increased risk for preterm birth (OR = 1.3, 95% CI: 1.0 to 1.7) per 1SD (0.36 mmol/L) increase in maternal FPG (Figure 3 and Supplementary Table 11).

To further understand whether the observed associations were driven by the SNPs that influence normal FPG levels or by the SNPs associated with pathological hyperglycemia (i.e. T2D), we tested associations between the T2D haplotype scores and pregnancy outcomes. Compared to the FPG scores, the maternal non-transmitted T2D score (*S*_h2_) was less significantly associated with both birth weight and gestational duration; but the negative associations between the paternal transmitted score (*S*_h3_) and birth weight and gestational duration were more apparent (Table 1). This indicates that the maternal causal effect is mainly driven by the quantitative level of FPG (indexed by the FPG scores); while the fetal genetic effect can be mediated through mechanisms other than normal control of blood glucose levels (indexed by the T2D SNPs), which is in agreement with the fetal insulin hypothesis(19).

As shown In Figure 3 and Supplementary Figure 13, the combinations of the estimated maternal causal effects and the genetically confounded associations due to genetic transmission were consistent with the observed phenotypic associations between maternal phenotypes and birth outcomes. The maternal causal effects were usually more dominant than the genetically confounded associations in shaping the phenotypic associations between maternal phenotypes and birth outcomes. In some cases, the maternal causal effects and the genetically confounded associations pointed to opposite directions (e.g. between height and gestational duration; between FPG and birth weight) and in these situations the estimated maternal causal effects could be larger than the observed phenotypic associations.

### Genetically confounded associations between birth outcomes and adult phenotypes in offspring

We estimate the magnitude of genetically confounded associations between the birth outcomes and adult phenotypes (in the offspring) based on the hypothesis that the association was driven by the variants that were associated with an adult phenotype were also associated with a birth outcome (Methods). The two methods (Method 1 and Method 2) generated similar results (Supplementary Table 12). As shown in Figure 4, 1 SD change in birth weight (gestational age adjusted) was estimated to be associated respectively with 0.20 SD (95% CI: 0.17 to 0.24) and 0.076 SD (95% CI: 0.014 to 0.138) differences in adult height and BMI. 1 SD increase in both gestational duration and birth weight were estimated to be associated with 0.05 SD decrease in adult blood pressure and 0.025-0.03 SD decrease in adult FPG level, and birth weight was also estimated to be negatively associated with susceptibility to T2D (OR = 0.95, 95% CI: 0.92 to 0.99). The genetically confounded associations between birth outcomes and adult phenotypes (Supplementary Figure 14 and 15) were mainly driven by the shared genetic effects in offspring; however, the maternal effects could substantially confound the associations and point to opposite directions to the confounded associations by fetal genetic effects in certain cases (e.g. the confounded associations between gestational duration and height, between birth weight and FPG or T2D risk). We compared the genetically confounded associations between birth weight and adult phenotypes with the estimated phenotypic associations from genetic correlation(20), the observed associations in the ALSPAC data and the reported associations from an epidemiological meta analysis(53) (Figure 5). The estimated genetically confounded associations using our approach were similar to those estimated from genetic correlations(20) and were largely consistent with the observed or reported associations between birth weight and adult phenotypes with the exception of body height, for which the observed association (from the ALSPAC data) was significantly stronger than the estimated confounded association due to shared genetics.

**Figure 4.**
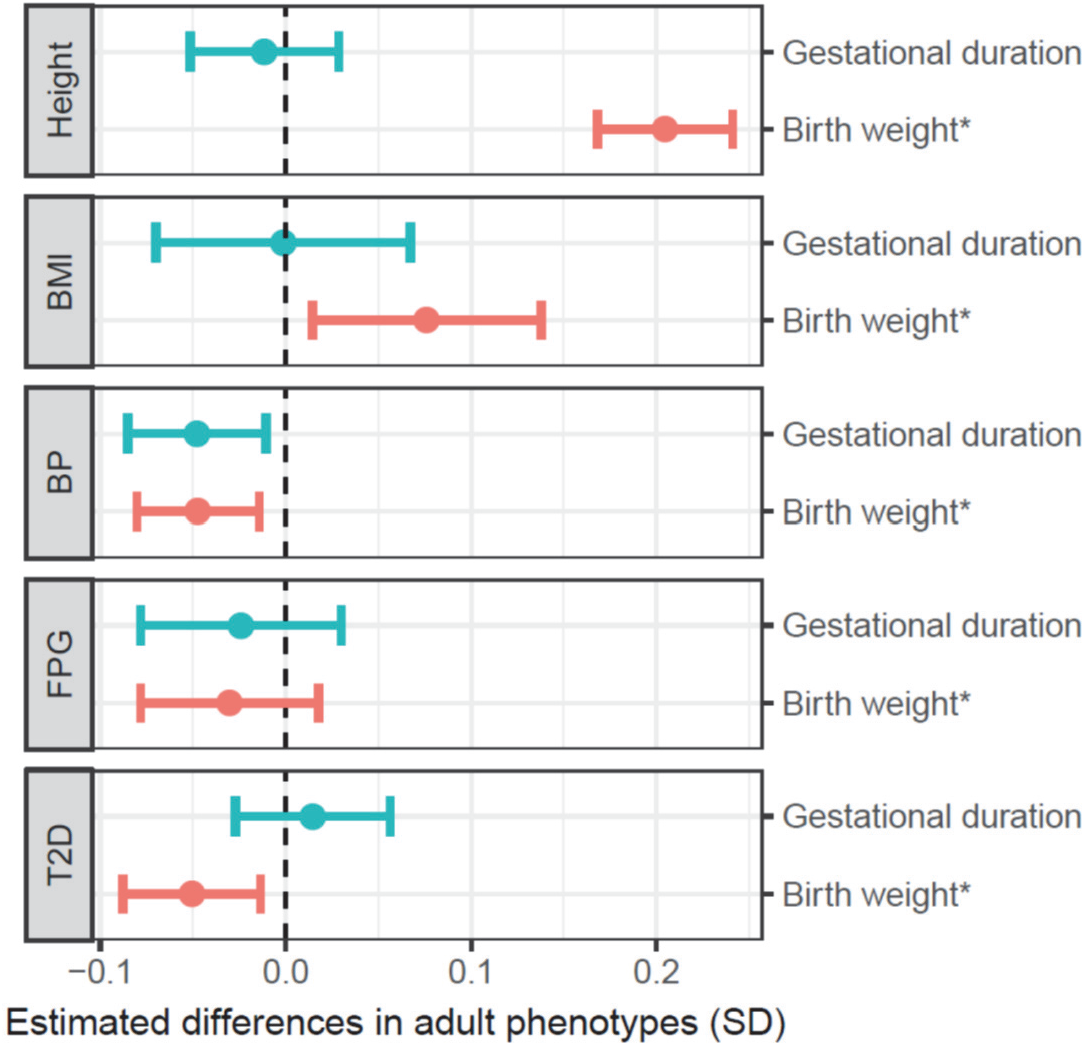
Estimated differences in adult phenotypes (in SD) per 1SD difference in birth weight and gestational duration. The birth weight* was adjusted by gestational duration; 1SD=426g. 1SD of gestational duration was 11.4 days.

**Figure 5.**
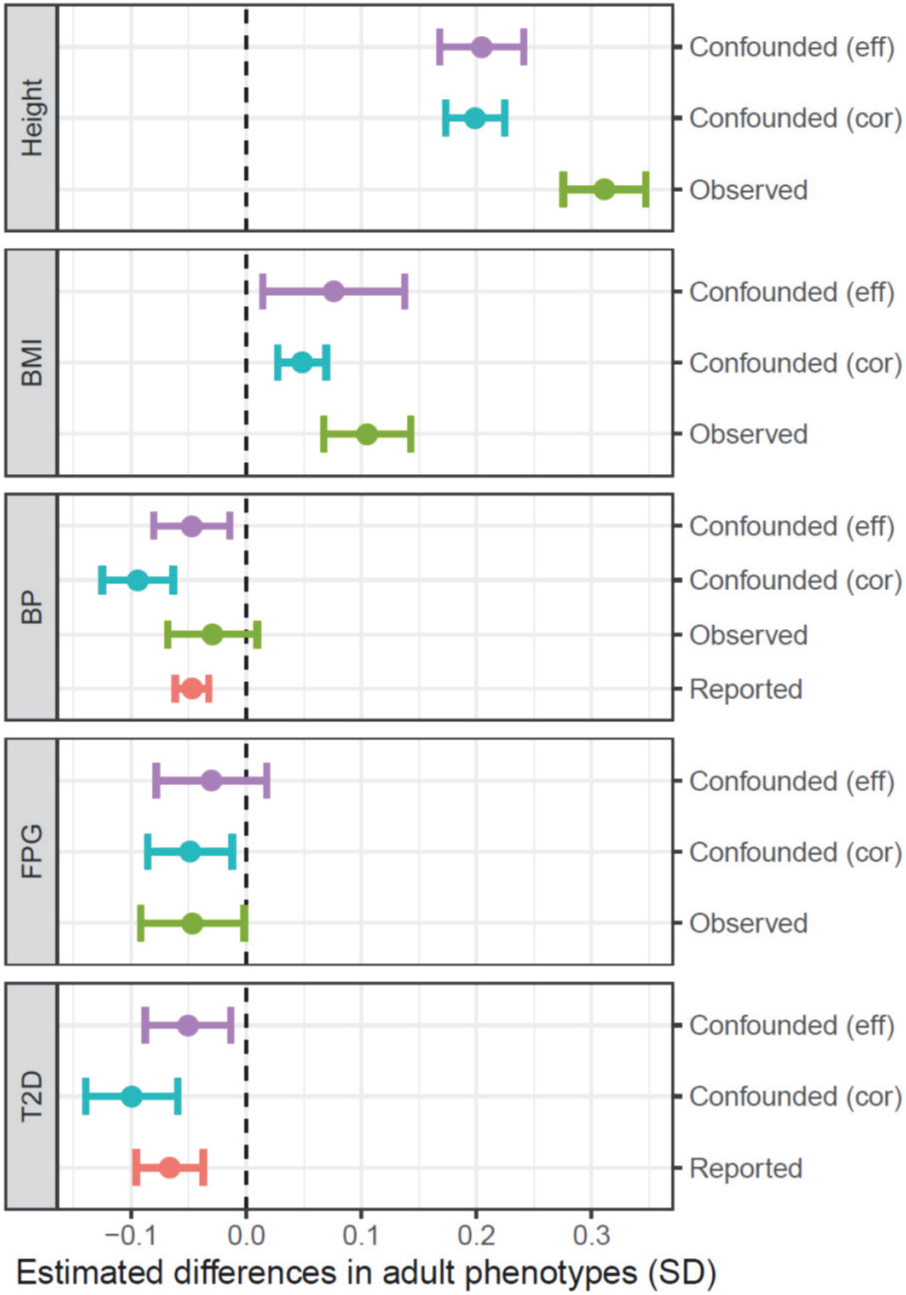
Comparing the magnitudes of genetically confounded associations (differences in adult phenotypes (in SD) per 1SD difference in birth weight) with observed and reported phenotypic associations. **<u>Confounded (eff):</u>** genetically confounded associations estimated based on the fetal genetic effect on birth weight (adjusted by gestational duration) of the variants associated with an adult phenotype. **<u>Confounded (cor)</u>**: genetically confounded associations between birth weight (unadjusted by gestational duration) and adult phenotype based on the reported genetic correlations (Ref: 20). **<u>Observed</u>**: observed phenotypic associations between gestational duration adjusted birth weight and offspring height, BMI, BP and FPG measured at age 17 in ASLPAC. **<u>Reported</u>**: the reported associations between birth weight (unadjusted by gestational duration) and BP and T2D susceptibility from a recent epidemiological study (Ref: 53). 1SD in the gestational duration adjusted and unadjusted birth weight were 426g and 484g, respectively.

### Causal effect of fetal growth on gestational duration, maternal systolic blood pressure and FPG

To test the possible fetal drive of fetal growth on birth outcomes and maternal pregnancy phenotypes, we constructed genetic scores using 86 SNPs associated with birth weight with confirmed fetal effect(28) and tested their associations with birth outcomes as well as maternal blood pressure and FPG during pregnancy (Table 2). The fetal genetic score for birth weight was significantly associated with gestational age adjusted birth weight with an *R*^2^ = 3.1%. The paternal transmitted haplotype score (*S*_h3_) consistently has larger effect than the maternal transmitted score (*S*_h1_) (Supplementary Table 13) probably due to a negative maternal effect of the same alleles on birth weight and birth length as shown by the negative effect of the non-transmitted haplotype score (*S*_h2_) (Table 2A). The maternal and paternal transmitted birth weight haplotype scores (*S*_h1_ and *S*_h3_) were also associated with shorter gestational duration and increased preterm birth risk (Table 2A). Using the paternal transmitted haplotype score as an instrument, the estimated causal effects were 3 days (95% CI: 1.5 to 5.0) shorter gestation and an approximate doubling of the preterm birth risk per 1SD changes in fetal growth rate (Table 2A and Supplementary Figure 16). In addition, we also observed significant associations between paternal transmitted birth weight score (*S*_h3_) and maternal SBP (P=0.022) measured during pregnancy and the estimated causal effects (Ratio method) were 1.4 mmHg (95% CI: 0.2 to 2.6) increase in SBP per 1SD changes in fetal growth rate (Table 2B and Supplementary Figure 17).

**Table 2.** Associations between birth weight genetic scores with birth outcomes and maternal blood pressures and glucose levels.

| Genetic score association & Causal estimation<sup>a</sup> | Gestational days |  |  | Preterm birth (log(OR)) |  |  | Birth weight (g) |  |  | Birth length (cm) |  |  |
| --- | --- | --- | --- | --- | --- | --- | --- | --- | --- | --- | --- | --- |
|  | beta | se | p-val | beta | se | p-val | beta | se | p-val | beta | se | p-val |
| Maternal trans ($\beta_{h1}$) | -0.0056 | 0.0027 | **0.035** | 0.0024 | 0.00064 | **0.00018** | 0.79 | 0.087 | **1.80E-19** | 0.0018 | 0.00048 | **0.00022** |
| Maternal non-trans ($\beta_{h2}$) | 0.001 | 0.0027 | 0.7 | -0.00058 | 0.00065 | 0.37 | -0.19 | 0.088 | **0.029** | -0.00022 | 0.00049 | 0.66 |
| Paternal trans ($\beta_{h3}$) | -0.0099 | 0.0026 | **0.00018** | 0.0019 | 0.00064 | **0.003** | 1.3 | 0.087 | **1.70E-48** | 0.0031 | 0.00048 | **2.30E-10** |
| Maternal effect ($\beta_{MY}$) | 0.0026 | 0.0023 | 0.26 | -5.80E-05 | 0.00055 | 0.92 | -0.33 | 0.076 | **1.10E-05** | -0.00073 | 0.00042 | 0.082 |
| Fetal effect ($\beta_{FY}$) | -0.0083 | 0.0023 | **0.00029** | 0.0025 | 0.00056 | **9.20E-06** | 1.1 | 0.075 | **1.00E-50** | 0.0025 | 0.00042 | **1.30E-09** |
| Causal (TSLs) | -2.92 | 0.805 | **0.00029** | 0.837 | 0.267 | **0.0017** | NA |  |  | 1.05 | 0.124 | **2.40E-17** |
| Causal (Ratio) | -3.24 | 0.889 | **0.00027** | 0.624 | 0.215 | **0.0036** |  |  |  | 1 | 0.172 | **5.10E-09** |

| Genetic score association & Causal estimation<sup>a</sup> | SBP<sup>b</sup> (mmHg) |  |  | DBP<sup>b</sup> (mmHg) |  |  | FPG<sup>c</sup> (mmol/L) |  |  |
| --- | --- | --- | --- | --- | --- | --- | --- | --- | --- |
|  | beta | se | p-val | beta | se | p-val | beta | se | p-val |
| Maternal trans ($\beta_{h1}$) | -0.0026 | 0.0019 | 0.17 | -0.00021 | 0.0013 | 0.87 | -0.00011 | 0.00022 | 0.61 |
| Maternal non-trans ($\beta_{h2}$) | 0.00057 | 0.0019 | 0.77 | 0.0006 | 0.0013 | 0.65 | -0.00025 | 0.00022 | 0.26 |
| Paternal trans ($\beta_{h3}$) | 0.0043 | 0.0019 | **0.022** | 0.0021 | 0.0013 | 0.1 | 8.90E-05 | 0.00022 | 0.69 |
| Maternal effect ($\beta_{MY}$) | -0.0032 | 0.0016 | 0.053 | -0.00083 | 0.0011 | 0.47 | -0.00022 | 0.0002 | 0.26 |
| Fetal effect ($\beta_{FY}$) | 0.00054 | 0.0016 | 0.74 | 0.00065 | 0.0011 | 0.57 | 1.10E-04 | 0.00019 | 0.55 |
| Causal (TSLs) | 1.2 | 0.551 | **0.03** | 0.641 | 0.378 | 0.09 | 0.0203 | 0.0565 | 0.72 |
| Causal (Ratio) | 1.41 | 0.621 | **0.023** | 0.69 | 0.426 | 0.1 | 0.029 | 0.0724 | 0.69 |

a: The effect sizes of genetic score association were given by per unit (g) change in genetic scores; the causal effect sizes were based on per 1SD (426g) change in age adjusted birth weight.
b: Maternal blood pressures in ALSPAC and HAPO
c: Fasting plasma glucose (FPG) during pregnancy measured in HAPO only

## DISCUSSION

In this report, we utilized a haplotype-based genetic score approach to explicitly model the maternal and fetal genetic effects (Figure 2). From the estimated maternal and fetal genetic associations (Table 1), we estimated maternal causal effects and the genetically confounded associations between maternal phenotypes and birth outcomes (Figure 3) as well as the genetically confounded associations between birth outcomes and adult phenotypes (Figure 4). We also tested whether fetal growth (as indicated by gestational age adjusted birth weight) could influence gestational duration, maternal blood pressure and FPG levels during pregnancy (Table 2). Our results revealed complex maternal and fetal genetic effects in shaping the associations between maternal phenotypes and birth outcomes and their associations with adult phenotypes (Figure 6).

**Figure 6.**
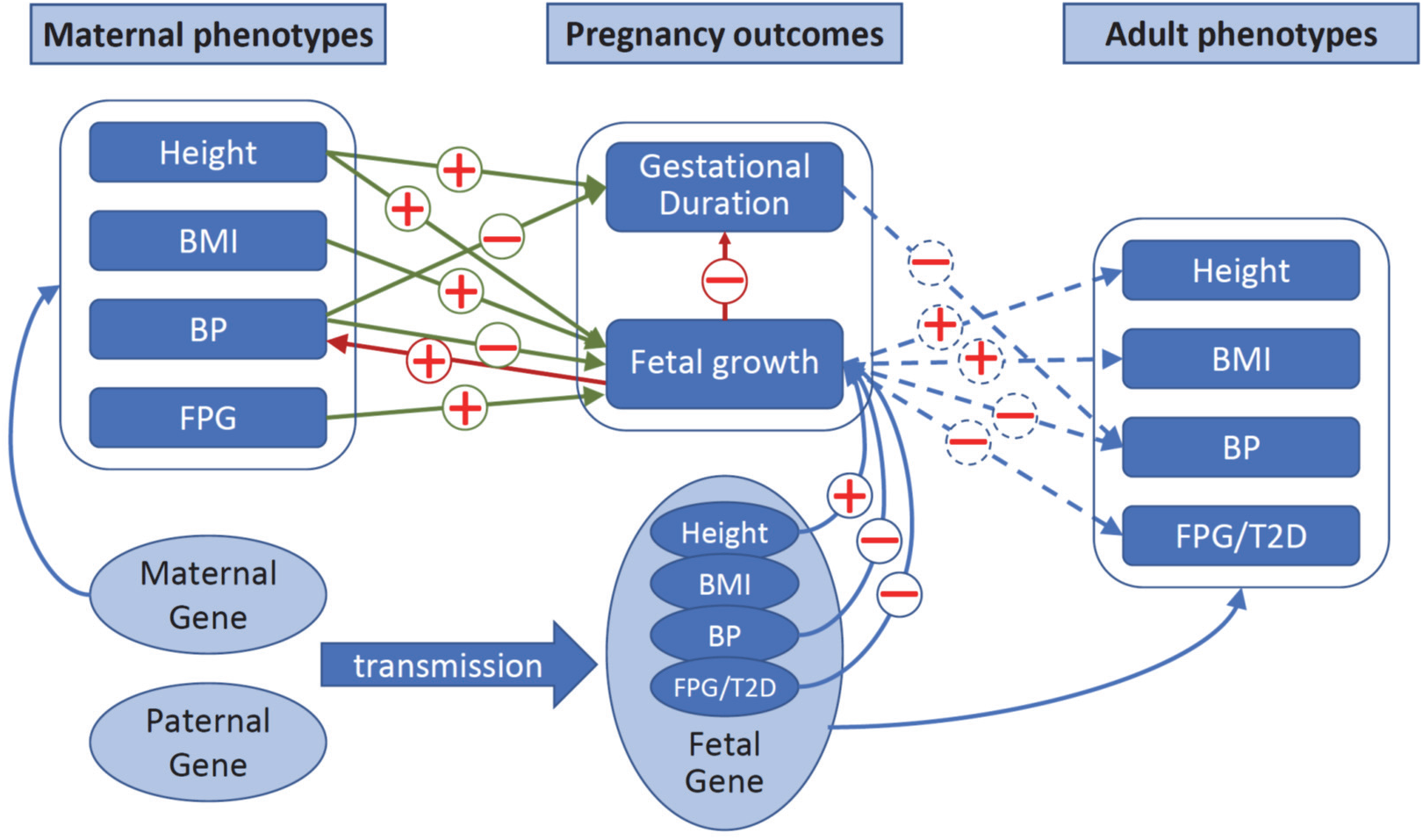
Estimated maternal and fetal genetic effects underlying the associations between maternal phenotypes and pregnancy outcomes and their associations with adult phenotypes. Blue arrows: maternal or fetal genetic effects. Green arrows: maternal causal effects. Red arrows: “fetal drive”. Dashed arrows: Genetically confounded associations between birth outcomes and late adult phenotypes due to shared genetics.

These findings have several implications. First, maternal size and fetal growth are important factors in defining the duration of gestation. This is demonstrated not only by the significant causal effect of maternal height (size of the mother) on gestational duration, but also by the observation that the maternal or fetal genetic effects on fetal growth are usually associated with opposing effects on gestational duration. For example, rapid fetal growth either due to high maternal FPG or direct fetal genetic effects of growth promoting alleles shortens gestational duration. Whether this “trade-off” between fetal growth and gestational duration is due to physical(54) or metabolic(55) constraints will require further investigation. Alleles associated with blood pressure have negative impacts on both birth weight and gestational duration, but mainly through either fetal genetics or a maternal effect respectively.

Secondly, fetal growth (as evaluated by gestational duration-adjusted birth weight and length) is influenced by both maternal and fetal effects. In addition to the strong causal effects of maternal FPG on birth weight, the many alleles associated with body height, blood pressure, FPG level or T2D susceptibility can influence fetal growth through fetal genetic effects. The alleles associated with body height are positively associated with birth size, while the alleles associated with higher metabolic risks (e.g. high blood pressure, blood glucose level and higher risk of T2D) reduce birth weight, which also suggests that lower birth weight (or more precisely small for gestational age status) might be a predictor of the load of genetic metabolic risks. Our results show that the shared genetic effects largely explain the life-course associations between birth weight and many cardiometabolic phenotypes (Figure 5) – a result that is consistent with the reported inverse genetic correlations between birth weight and late-onset metabolic disorders(20) and both support a strong genetic rather than an environmental effect underlying the life-course association between birth weight and later metabolic risks. Compared with the previous analyses based on genetic correlations using genome-wide SNPs, our approach estimated the life-course associations by extrapolating the effects of the top GWA SNPs associated with the adult phenotypes and therefore has a specific mechanistic implication – the life-course associations were mainly driven by genetic variants with large effects on adult phenotypes rather than by the variants with large effects on fetal growth. In addition, our method can partition the genetically confounded associations to either maternal effects or shared genetic effects in offspring (Supplementary Methods).

The results from this study also reveal a major theme in human pregnancy – both maternal effects and direct fetal genetic effects jointly shape the observed associations between maternal phenotypes and pregnancy outcomes. The same genetic variants can influence different birth outcomes or the same birth outcomes through both maternal and fetal effects, and these two types of effects can be antagonistic, as exemplified by the opposing maternal and fetal effects of the FPG associated alleles on birth weight, or the effects of height associated alleles on gestational duration. These complex mechanisms can be further complicated by “fetal drive” – as shown by the associations between paternal transmitted birth weight genetic score and gestational duration as well as maternal blood pressure, i.e. fast fetal growth shortens gestational duration and increases maternal blood pressure (Table 2).

Our study had a number of limitations. First, there was some incomplete and heterogeneous phenotype data. Most maternal phenotypes (e.g. age, height and BMI) and birth outcomes (e.g. gestational duration and birth weight) were available from the study data sets; however, FPG level during pregnancy was only available in HAPO and blood pressure data were only available from HAPO and ALSPAC. Another limitation is that due to incomplete data, we were not able to include important environmental or socioeconomic factors in the analysis. However, we argue that the genetic scores used in our study are not known to be associated with these factors, and therefore our analyses are robust to potential confounding due to these environmental factors. Biological pleiotropy is always an issue in causal inference using genetic instruments, especially when a large number of genetic variants are used(45, 56). We used the MR-PRESSO method to detect and remove SNPs with horizontal pleiotropic effects (Supplementary methods). The results (Supplementary Table 14 and 15) were essentially the same as the corresponding haplotype genetic score associations (Table 1 and Table 2). There is some evidence of pleiotropy (MR-PRESSO global test P < 0.05), suggesting the identified maternal effects may be not exclusively mediated by the targeted maternal phenotypes and the fetal effects of the SNPs on a birth outcome are not always proportional to their report effects on the adult phenotype (Supplementary Figure 2). After excluding the outliers, the corrected estimates and the p-values did not change substantially, and all the MR-PRESSO distortion test p-values were nonsignificant (Supplementary Table 14, 15 and Supplementary Figure 18).

To conclude, our study revealed that many SNPs associated with maternal height, blood pressure, blood glucose levels (or T2D susceptibility) can have various maternal and fetal genetic effects on gestational duration and fetal growth. These maternal and fetal genetic effects can largely explain the observed associations between the studied maternal phenotypes and birth outcomes as well as the life-course associations between these birth outcomes and adult phenotypes. We show that rapid fetal growth might reduce gestational duration and increase maternal blood pressure. These findings provide novel insights into the mechanisms behind the observed associations between maternal phenotype and birth outcomes and their life course impacts on later-life health.

## ACKNOWLEDGEMENTS

FIN: We thank the participants in the Finnish birth cohort as well as the research group who collected the data.

MoBa: We are grateful to all the participating families in Norway who take part in this cohort study.

ALSPAC: We are extremely grateful to all the families who took part in this study, the midwives for their help in recruiting them, and the whole ALSPAC team, which includes interviewers, computer and laboratory technicians, clerical workers, research scientists, volunteers, managers, receptionists and nurses.

DNBC: We are very grateful to all DNBC families who took part in the study. We would also like to thank everyone involved in data collection and biological material handling.

HAPO: would like to acknowledge the participants and research personnel at the participating HAPO field centers.

GPN: We thank the infants and their parents who agreed to take part in this study and the medical and nursing colleagues who collected that data.

dbGaP: We thank dbGAP for depositing and hosting the phenotype and genotype data of the DNBC, HAPO and GPN data sets.

