## Supplementary Material for "Haplotype genetic score analysis in 10,734 mother/infant pairs reveals complex maternal and fetal genetic effects underlying the associations between maternal phenotypes, birth outcomes and adult phenotypes" (from another version of this preprint: version 1, 2019-08-15)

### Title Page

#### Title

### Supplementary Methods

#### Data sets

We used phenotype and genomewide SNP data of 10,734 mother/infant pairs from six birth studies collected from European countries and United States (Table S1).

##### *FIN*

The Finnish dataset (FIN) was collected for a genetic study of spontaneous preterm birth[1]. Briefly, whole blood samples were collected from more than 1,600 mother/child pairs from the Helsinki (southern Finland) University Hospitals between 2004 and 2014. All the studied samples are of Finnish descent. Crown-rump length at the first ultrasound screening between 10+ and 13 weeks was used to determine the gestational age. 2,962 blood samples from mothers and children were genotyped. After genotype quality control (QC) procedure and applying the phenotype-based inclusion/exclusion criteria, 1,170 mother/child pairs were selected and used in the analysis. The study was approved by the Ethics Committee of Oulu University Hospital and that of Helsinki University Central Hospital. Written informed consent was given by all participants.

##### *MoBa*

The Mother Child dataset of Norway (MoBa) is a nationwide Norwegian pregnancy study administered by the Norwegian Institute of Public Health. The study includes more than 114,000 children, 95,000 mothers and 75,000 fathers recruited from 1999 through 2008[2]. Gestational age was estimated by ultrasound at gestational weeks 17–19. In the few cases without ultrasound dating, gestational age was estimated using the date of the last menstrual period. For the current study, we used the mother-child pairs that were selected from Version 4 of the MoBa dataset, which included a total of 71,669 pregnancies[3]. Singleton live-born spontaneous pregnancies with mothers in the age group 20–34 years were selected. Random sampling was done from two gestational age ranges 154-258 days (cases) and 273–286 days (controls). Pregnancies involving pre-existing medical conditions, pregnancies with complications as well as pregnancies conceived by in vitro fertilization, were excluded from the study. In total, blood samples from 3,120 mothers and children were genotyped[3]. 1,009 mother/child pairs that passed QC and inclusion/exclusion criteria were included in the analysis. All parents gave informed, written consent. The study was approved by The Regional Committee for Medical Research Ethics in South-Eastern, Norway.

*DNBC*

The Danish National Birth Cohort (DNBC) followed over 100,000 pregnancies between 1996 and 2003 with extensive epidemiologic data on health outcomes in both mother and child[4]. The current study used the data downloaded from the Database of Genotypes and Phenotypes (dbGaP) (phs000103.v1.p1), which contains data from a genome-wide case/control study using approximately 1,000 preterm mother-child pairs (gestational age between 22–37 weeks) from the DNBC, along with 1,000 control pairs in which the child was born at ~40 weeks gestation. Gestational duration in this dataset was determined by a consensus algorithm combining all available information from multiple sources: self-reported date of last menstrual period, self-reported delivery date, and gestational age at birth registered in the Medical Birth Register and the National Patient Register. Of the 3,886 samples with genotype data, we identified 1,739 mother/child pairs and used them in the analysis. The study protocol was approved by the Danish Scientific Ethical Committee and the Danish Data Protection Agency.

*HAPO*

The Hyperglycemia and Adverse Pregnancy Outcome (HAPO) Study is a multicenter, international study in which high quality phenotypic data related to fetal growth and maternal glucose metabolism has been collected from 25,000 pregnant women of varied racial and socio-demographic backgrounds using standardized protocols that were uniform across centers. For the current study, we utilized phenotype and genotype data of 1,500 infants and their mothers of European descent download from dbGaP (phs000096.v2.p1). Gestational duration in this dataset was determined by last menstrual period or ultrasound estimation from 6-24 weeks. Of the 2,866 samples with genotype data, we identified 1,089 mother/child pairs and used them in the analysis.

*GPN*

The Genomic and Proteomic Network for Preterm Birth Research (GPN) study is a multicenter observational genome-wide association study (GWAS) designed to determine the genetic predisposition to idiopathic preterm birth. Phenotype data and genotype data from 743 spontaneous preterm births (20 to less than 34 weeks gestation), and 752 controls (39 to less than 42 weeks gestation) of diverse ethnic background (White, Hispanics, African Americans, and Others) were collected. For this current study, we identified 343 mother/child pairs of European descent from the data downloaded from dbGaP (phs000714.v1.p1).

##### *ALSPAC*

The Avon Longitudinal Study of Parents and Children (ALSPAC) is a prospective birth cohort study. 14,541 pregnant women resident in the former county of Avon (situated around the city of Bristol in the South West of England) with expected dates of delivery 1st April 1991 to 31st December 1992 were recuited[5,6]. The children arising from these women, and their partners were followed up intensively over nearly three decades. Genotype data of the mothers and children were generated using the Illumina HumanHap550 quad (children) and Illumina human660W quad (mothers). This resulted in a dataset of 17,842 participants (either mothers or offspring), containing 6,305 mother-offspring pairs, each with 465,740 SNPs genotyped. From this data set, 5,384 mother-offspring pairs who passed genotype QC and inclusion/exclusion criteria were included in the analysis. Please note that the study website contains details of all the data that is available through a fully searchable data dictionary and variable search tool (http://www.bristol.ac.uk/alspac/researchers/our-data/).

#### Phenotype data

##### *Phenotype measures*

In this study, we focused on maternal height, pregnancy BMI, blood pressure and blood glucose levels measured during pregnancy and pregnancy outcomes including gestational duration (and preterm birth as a binary trait), birth weight and birth length. Maternal height, weight and most of the birth outcomes were available in most of the data sets (except for birth weight and length were not available from MoBa and birth length was not available in the DNBC data set used in this study).

Maternal blood pressure data was available in ALSPAC and DNBC. In the ALSPAC study, all blood pressure measurements taken during antenatal care were extracted from medical records with each woman having a median (interquartile range) of 13 (11, 16) measurements between 8 weeks and delivery[7]. In the HAPO study, blood pressure was measured once between 24 and 32 (mean 28) weeks when the mothers underwent a 75-g oral glucose tolerance test.

In the HAPO study, fasting plasma glucose (FPG) levels in the mothers were measured between 24 and 32 weeks gestation as part of the standard oral glucose tolerance test. In the ALSPAC study, FPG was measured in some mothers in a follow-up data collection 18 years after the pregnancy.

*Inclusion/exclusion criteria*

We studied singleton pregnancies with spontaneous live births – including spontaneous deliveries with or without premature rupture of membranes (PROM). C-sections after spontaneous onset of labor were retained. Medically indicated deliveries or C-sections and repeat C-sections were excluded. Mother/child pairs without gestational duration information were excluded. Pregnancies with known gestational or fetal complications (e.g. placental abnormities, chorioamnionitis, preeclampsia, and congenital anomalies) and pregnancies involving pre-existing medical conditions (i.e. hypertension or diabetes) or maternal risk exposure (e.g. drug use during pregnancy) known be associated with preterm birth were also excluded. Pregnancies with gestational diabetes and gestational hypertension were not excluded for the investigation of maternal physiological changes during pregnancy.

#### Genotype data

The genomewide SNP data of the data sets were generated using either Affymetrix 6.0 or various Illumina arrays. After genotype calling, similar genotype QC procedures were applied. An overview of the data procedure was shown in Figure S1.

*Genotype calls*

For the FIN dataset, genotyping was conducted using Affymetrix 6.0 (Affymetrix, California, United States) and various other Illumina arrays (Illumina, California, United States). For the Affymetrix SNP Array 6.0, genotype calls were determined using the CRLMM algorithm[8,9] among chips that passed the vendor-suggested QC (Contrast QC > 0.4). For the Illumina chips, the genotype calling was conducted using Illumina's genotyping module v1.94 in the GenomeStudio v2011.1.

The samples from the MoBa dataset were genotyped using the Illumina Human660W-Quadv1_A bead chip (Illumina Inc.) and the genotype calls were determined using CRLMM algorithm.

The DNBC samples were genotyped using Human660W-Quad bead arrays from Illumina. The raw genotype intensity data (*.idat) files were obtained from dbGaP and we performed genotype calls using CRLMM algorithm.

The HAPO samples of European descent were genotyped using Human610-Quad array. We obtained the raw genotype intensity data (*.idat) files performed genotype calls using CRLMM algorithm.

The processed genotype calls in plink format of the GPN data set was obtained from dbGaP (phs000714.v1.p1). Data from participants of apparent duplications with others (IBD>0.8), with sex discrepancies (between known sex and genetically inferred sex), and children with high Mendelian errors (>10%) were deleted.

The ALSPAC genotype data were generated using the Illumina HumanHap550 quad (children) and Illumina human660W quad (mothers). The cleaned genotype calls of 465,740 SNPs of 17,842 subjects were obtained from ALSPAC.

##### *Genotype QC procedures*

We performed similar genotype QC across of all the dataset (Figure S1). We first performed sample-level QC based on call rate, overall heterogeneity, sex discrepancies; we checked the pedigree relationship based IBD analysis; Genetic ancestry was assessed by principal components analysis (PCA) anchored by 1000 Genomes reference samples. Individuals with non-European ancestry were excluded. We then perform marker level QC: SNPs with low call rate (<98%), low minor allele frequency (<0.01) or significant deviation from Hardy-Weinberg Equilibrium ($P < 5\times10^{-6}$) were excluded.

##### *Genomewide phasing, imputation and inference of allelic transmission*

The genotype data of the mother and infants was phased together using Shapeit2 software[10] and then the estimated haplotypes were used to impute untyped SNPs using the reference haplotypes extracted from the Phase III 1000 Genomes Project[11] by Minimac3[12]. The genotype data of the FIN data set was generated using multiple platforms. The pre-phasing and imputation were done separately by each platform. To fully utilize the mother/child pair relationship across all the sample, we merged the imputed data across all the samples and conducted re-phasing using Shapeit2 (Figure S1).

The haplotype phasing information in mother/infant pairs was used to infer allele transmission between mothers and their infants. When phasing in mother/infant duos, the Shapeit2 automatically accommodates the mother/child relationship and impose constraints on each individual's haplotypes based on allele transmission between a mother and her child[10]. We did not use the duoHMM argument (for estimating haplotypes in general pedigrees[13]) when running the Shapeit2 as we only included mother/infant duos from the data sets and excluded other close related samples. Without invoking the duoHMM argument, the first phased haplotype of a mother/child pair reported by Shapeit2 is the transmitted haplotype.

#### Genetic scores

##### *Selection of GWA SNPs*

Reported GWA SNPs robustly associated with body height, BMI, blood pressure, fasting plasma glucose (FPG) levels and type 2 diabetes susceptibility (T2D) to construct weighted genetic scores to instrument various maternal phenotypes (Table S4). In addition, we also constructed genetic scores for birth weight.

**Height and BMI**: The GWA SNPs associated with height and BMI reported by the 2018 GIANT and UK BioBank Meta Analysis[14] were download from the GIANT website (https://portals.broadinstitute.org/collaboration/giant/). From the top 3,290 height SNPs based on conditional & joint COJO analysis (COJO), we selected 2,390 SNPs with primary P-value < 1E-8 and among these 2,382 SNPs were diallelic and polymorphic in the 1000 Genomes European reference samples. Further, we excluded 141 SNPs because they were in substantial LD ($r^2 > 0.2$) with other more significant SNPs. As a result, 2,214 SNPs and their reported beta values were used to construct height genetic scores. Following a similar procedure, we selected 643 SNPs from the reported top 941 BMI SNPs to construct BMI genetic scores.

**Blood pressure**: For blood pressure, we extracted the 885 SNPs and their estimated effect sizes from the most recent GWAS[15] (Supplementary Table 24 of the original report). Among this list 5 SNPs were not polymorphic in the 1000 Genomes European reference samples, one SNP had zero effect and one was in close LD with another more significant SNP, were excluded. The final list included 878 SNPs. These SNPs together with their estimated effects on systolic (SBP), diastolic blood pressure (DBP) were used to build genetic scores for SBP and DBP, respectively. For simplicity, we also build a score for average blood pressure (BP) using the mean estimated effects of SBP and DBP.

**Blood glucose and T2D susceptibility**: 23 GWA SNPs associated with fasting glucose levels (FPG) identified from 96,496 non-diabetic individuals[16] and their estimated effect sizes accounted for BMI were used to construct genetic scores for blood glucose levels. In addition, we also constructed genetic scores for susceptibility to Type 2 diabetes (T2D) using the SNPs reported by a recent GWA study in 898,130 European-descent[17]. Among the 403 reported SNPs (Supplementary Table 4 of the original report), 21 SNPs were excluded either due to non-polymorphic or LD redundancy. To contrast with the genetic regulation of normal glucose levels, we also excluded 13 SNPs which were in close LD with any of those 23 GWA SNPs associated with normal fasting glucose levels. The final list of 369 SNPs and the log transformed BMI adjusted odds ratios (ORs) were used to construct T2D genetic scores.

**Birth weight**: We selected GWA SNPs associated with birth weight through fetal genetic effect from a recent GWA study using a large number of European samples[18]. This study used a structural equation modelling (SEM) method[19] to separate maternal from fetal effects. From the total 209 genome-wide significant SNPs (Supplementary Table 6 of the original report), we used 106 SNPs with confirmed fetal genetic effects (SEM classification: "Fetal Only" or "Fetal and Maternal"). After removing 10 SNPs due to LD redundancy or co-locating with other SNP, 96 SNPs and their fetal effects estimated by the SEM method were used to construct birth weight genetic score and to model fetal genetic effect on fetal growth.

##### *Construction of genetic scores*

The weighted genetic score was constructed as $S = \sum b_i G_i$, where $G_i = 0, 1 \text{ or } 2$ (for genotype) or $G_i = 0 \text{ or } 1$ (for haplotype genetic score), which indicates the number of reference alleles for a specific SNP, and $b_i$ is the estimated allelic effect reported by the GWA studies. For a set of GWA SNPs associated with a particular maternal (adult) phenotype, we constructed two genotype genetic scores: $S_{\text{mat}}$ (maternal genotype score), $S_{\text{fet}}$ (fetal genotype score); and three haplotype genetic scores: $S_{\text{h1}}$, $S_{\text{h2}}$ and $S_{\text{h3}}$ respectively based on the maternal transmitted (h1), maternal non-transmitted (h2) and paternal transmitted alleles (h3) (Figure 1). Obviously, the maternal genotype score is the summation of the two maternal (transmitted and non-transmitted) haplotype scores ($S_{\text{mat}} = S_{\text{h1}} + S_{\text{h2}}$); the fetal genotype score is the summation of the two transmitted haplotype scores ($S_{\text{fet}} = S_{\text{h1}} + S_{\text{h3}}$)

#### Statistical analyses

##### *Phenotypic associations*

We first assessed statistical associations between the maternal phenotypes ($X$) and the pregnancy outcomes ($Y$) using regression models with a common form: $Y \sim X + \text{Cov}$, where $\text{Cov}$ is a list of appropriate covariates. Specifically, the regression models are:

- $\text{gday} \sim \text{age1} + \text{age2} + \text{ht} + \text{bmi} + \text{sex}$
- $\text{pre} \sim \text{age1} + \text{age2} + \text{ht} + \text{bmi} + \text{sex}$
- $\text{bwt} \sim \text{age1} + \text{age2} + \text{ht} + \text{bmi} + \text{sex} + \text{gday1} + \text{gday2} + \text{gday3}$
- $\text{blen} \sim \text{age1} + \text{age2} + \text{ht} + \text{bmi} + \text{sex} + \text{gday1} + \text{gday2} + \text{gday3}$

In the above models, **gday** is the gestational duration measured in days, **pre** is the binary preterm birth status, **sex** is the fetal sex, **ht** and **bmi** are maternal prepregnant height and BMI and **age1** and **age2** are the first two polynomials of maternal age to capture the nonlinear effect of maternal age on pregnancy outcomes. Since birth weight (**bwt**) and birth length (**blen**) were primarily defined by gestational duration (**gday**), the first three polynomials (**gday1**, **gday2** and **gday3**) of gestational duration were included in regression models for **bwt** and **blen** to account for the strong effects of gestational duration on these birth outcomes.

In the HAPO and ALSPAC data sets, we also tested the impacts of maternal blood fasting glucose levels (**fpg**) and blood pressures (**sbp** and **dbp**) on the pregnancy outcomes using similar regression models by adding these variables as additional independent variable in the above regression models.

##### *Associations between genetic scores and maternal phenotypes*

Associations between the various genetic scores and their corresponding maternal phenotypes were examined using regression analyses. In the analysis of maternal height (**ht**) and maternal pre-pregnancy BMI (**bmi**), the first two polynomials of maternal age (**age1** and **age2**) were included as covariates. In the analysis of maternal blood pressure measures (**sbp** and **dbp**) and maternal fasting glucose levels (**fpg**), maternal height (**ht**), maternal BMI (**bmi**) and fetal sex (**sex**) were also included as covariates, as these factors were significantly associated with the maternal blood pressure (**sbp** and **dbp**) and maternal fasting glucose levels (**fpg**) measured during pregnancy.

- $\mathrm{ht} \sim \mathrm{scores} + \mathrm{age1} + \mathrm{age2}$
- $\mathrm{bmi} \sim \mathrm{scores} + \mathrm{age1} + \mathrm{age2}$
- $\mathrm{bp} \sim \mathrm{scores} + \mathrm{age1} + \mathrm{age2} + \mathrm{height} + \mathrm{bmi} + \mathrm{sex}$
- $\mathrm{fpg} \sim \mathrm{scores} + \mathrm{age1} + \mathrm{age2} + \mathrm{height} + \mathrm{bmi} + \mathrm{sex}$

##### *Associations between genetic scores and pregnancy outcomes*

To investigate the maternal or fetal genetic effects of a genetic score on a birth outcome, we tested associations between the three haplotype genetic scores and the pregnancy outcomes using regression models like those used in the observational association analyses, except the maternal phenotype under consideration was replaced by the three haplotype genetic scores ($S_{\mathrm{h1}} + S_{\mathrm{h2}} + S_{\mathrm{h3}}$). Hence the regression models for haplotype genetic score analysis take the form: $Y \sim S_{\mathrm{h1}} + S_{\mathrm{h2}} + S_{\mathrm{h3}} + \mathrm{Cov}$. For example, the following regression models were used to examine the association between height genetic scores and birth outcomes:

- $\text{gday} \sim S_{\text{h1}} + S_{\text{h2}} + S_{\text{h3}} + \text{age1} + \text{age2} + \text{bmi} + \text{sex}$
- $\text{pre} \sim S_{\text{h1}} + S_{\text{h2}} + S_{\text{h3}} + \text{age1} + \text{age2} + \text{bmi} + \text{sex}$
- $\text{bwt} \sim S_{\text{h1}} + S_{\text{h2}} + S_{\text{h3}} + \text{age1} + \text{age2} + \text{bmi} + \text{sex} + \text{gday1} + \text{gday2} + \text{gday3}$
- $\text{blen} \sim S_{\text{h1}} + S_{\text{h2}} + S_{\text{h3}} + \text{age1} + \text{age2} + \text{bmi} + \text{sex} + \text{gday1} + \text{gday2} + \text{gday3}$

The associations between these haplotype-based scores ($S_{\text{h1}}$, $S_{\text{h2}}$ and $S_{\text{h3}}$), i.e. the estimates of the regression coefficients ($\hat{\beta}_{\text{h1}}$, $\hat{\beta}_{\text{h2}}$, and $\hat{\beta}_{\text{h3}}$) and their associated p-values can explicitly model maternal effect vs fetal genetic effect (Figure 2)[20,21]. In particular, an association between $S_{\text{h2}}$ (maternal non-transmitted haplotype score) with a pregnancy outcome is a clear sign of maternal effect; whereas an association with $S_{\text{h3}}$ (paternal transmitted haplotype score) indicates direct fetal genetic effect on a pregnancy outcome.

##### *Modeling of maternal and fetal genetic effects using linear hypotheses*

In addition, we modeled the maternal effect and the fetal effect as the linear combinations of the regression coefficients ($\beta_{\text{h1}}$, $\beta_{\text{h2}}$, $\beta_{\text{h3}}$) of the three haplotype genetic scores (Figure 2). Under the assumptions of additivity between maternal and fetal effect and zero parent of origin effect, the effect ($\beta_{\text{h1}}$) of the maternal transmitted haplotype (h1) should equal to the summation of the maternal effect ($\beta_{\text{h2}}$) of the non-transmitted haplotype (h2) and fetal genetic effect ($\beta_{\text{h3}}$) of the paternal transmitted haplotype (h3). Thus, ($\beta_{\text{h1}} - \beta_{\text{h3}}$) and ($\beta_{\text{h1}} - \beta_{\text{h2}}$) respectively represent the maternal effect and the fetal genetic effect of the maternal transmitted haplotype (h1). Therefore, we can model the maternal and fetal genetic effect on a pregnancy outcome as:

- <u>Maternal effect</u>: $\beta_{MY} = (\beta_{\text{h1}} + \beta_{\text{h2}} - \beta_{\text{h3}})/2$ models the joint effect of the two maternal haplotypes ($\beta_{\text{h1}} + \beta_{\text{h2}}$) and excludes the possible fetal effect of the transmitted haplotype h1 by subtracting $\beta_{\text{h3}}$.
- <u>Fetal genetic effect</u>: $\beta_{FY} = (\beta_{\text{h1}} - \beta_{\text{h2}} + \beta_{\text{h3}})/2$ models the joint effect of the maternal and the paternal transmitted haplotypes ($\beta_{\text{h1}} + \beta_{\text{h3}}$) and excludes the possible maternal effect of the transmitted haplotype h1 by subtracting $\beta_{\text{h2}}$.

Modeling of maternal and fetal genetic effect using the linear combinations of the regression coefficients of the three haplotype genetic scores are more powerful (hence smaller standard error) than a single haplotype score (e.g. $S_{\text{h2}}$ for maternal effect and $S_{\text{h3}}$ for fetal effect) when the assumptions of additivity between maternal and fetal effect and zero parent of origin effect ($\beta_{\text{h1}} = \beta_{\text{h2}} + \beta_{\text{h3}}$) hold.

*Estimation of maternal causal effects*

The effect size estimates of the maternal non-transmitted haplotype score ($\hat{\beta}_{\mathrm{h2}}$) or the linear combination ($\hat{\beta}_{MY} = (\beta_{\mathrm{h1}} + \widehat{\beta_{\mathrm{h2}}} - \beta_{\mathrm{h3}})/2$) for maternal effect can be interpreted as a two-sample Mendelian randomization estimates of maternal causal effect. The first estimate ($\hat{\beta}_{\mathrm{h2}}$) based on the maternal non-transmitted alleles (h2), exclusively instruments the maternal effect. The second estimate ($\hat{\beta}_{MY}$) based on the linear combination of the regression coefficients of the three haplotype scores, although less specific (because it includes the transmitted alleles and needs the assumptions of additivity and zero parent of origin effect), is more powerful because it also considers the maternal effect of the transmitted alleles in instrumenting the maternal effect. These estimates evaluate the change in a pregnancy outcome caused by *certain amount* of change in a maternal phenotype caused by one unit change in the genetic score (which can cause exactly one unit change in the adult phenotype).

For a maternal phenotype (e.g. maternal pre-pregnancy height), the weighted genetic score was constructed using the GWA SNPs and their estimated effect sizes of a "same" adult phenotype (e.g. adult height). In the reference study, one unit genetic score should associated exactly with one unit difference in the adult phenotype; however, it may not associated with the same amount of difference in the corresponding maternal phenotype because of the heterogeneity due to different sex, age, etc from the reference study. For example, one unit (mmHg) SBP and DBP genetic score only associated with 0.27mmHg and 0.32mmHg changes in maternal SBP and DBP in this study (Table S5). Therefore, the effect size estimate ($\hat{\beta}_{\mathrm{h2}}$ or $\hat{\beta}_{MY}$) needs to be adjusted by their effect on the maternal phenotype ($\beta_{MX}$) to reflect the real magnitude of the causal effect (i.e. amount of change in a pregnancy outcome caused by one unit change in the maternal phenotype). We estimated the maternal causal effect by the ratio estimate[22]:

$$\hat{\beta}_{XY}^{m} = \hat{\beta}_{MY}/\hat{\beta}_{MX}$$

The standard error of can be obtained based on the asymptotic normal approximation[23]:

$$\mathrm{se}(\hat{\beta}_{XY}^{m}) = \sqrt{\frac{\mathrm{se}(\hat{\beta}_{MY})^2}{\hat{\beta}_{MX}{}^2} + \frac{\hat{\beta}_{MY}{}^2\,\mathrm{se}(\hat{\beta}_{MX})^2}{\hat{\beta}_{MX}{}^4}}$$

As an alternative, we also utilized the maternal non-transmitted haplotype score ($S_{\mathrm{h2}}$) as genetic instrument[21] and estimated the maternal causal effects using the two-stage least squares analysis (TSLS)[22].

*Estimation of genetically confounded associations due to shared genetic effects*

The effect size estimates of the paternal transmitted haplotype score ($\hat{\beta}_{\mathrm{h3}}$) or the linear combination ($\hat{\beta}_{FY} \; = \; (\beta_{\mathrm{h1}} - \widehat{\beta_{\mathrm{h2}}} + \beta_{\mathrm{h3}})/2$) for fetal genetic effect reflect the genetic variants associated with an adult phenotype have (shared) genetic effect on a pregnancy outcome. Quantitatively, they describe the amount of change in a pregnancy outcome caused by the genetic alleles that are associated with one unit difference in an adult phenotype. This shared genetic effect of the alleles associated with an adult phenotype $X'$ (or the corresponding maternal phenotype $X$) on a pregnancy outcome ($Y$) can confound the association between a maternal phenotype ($X$) and a pregnancy outcome ($Y$) as well as the association between the pregnancy outcome ($Y$) and the adult phenotype ($X'$) in offspring (Figure 2). In the following, we provided the mathematics for estimating these confounding associations from the estimated shared genetic effect ($\hat{\beta}_{FY}$).

Considering a general scenario (Figure S2), in which two random variables ($X$ and $Y$) are influenced by a confounder $Z$. The correlation between $X$ and $Y$ due to confounding is:

$$\rho'_{XY} = \rho_{ZX}\rho_{ZY} = \rho_{ZX}^2\rho_{ZY}/\rho_{ZX}$$

If expressing the correlations using regression coefficients following ($\rho_{XY} = \beta_{XY}\frac{\mathrm{sd}(X)}{\mathrm{sd}(Y)}$), the formula can be written as:

$$\beta'_{XY}\frac{\mathrm{sd}(X)}{\mathrm{sd}(Y)} = \rho_{ZX}^2\left(\beta_{ZY}\frac{\mathrm{sd}(Z)}{\mathrm{sd}(Y)}\right)\Big/\left(\beta_{ZX}\frac{\mathrm{sd}(Z)}{\mathrm{sd}(X)}\right)$$

which gives:

$$\beta'_{XY} = \rho_{ZX}^2\,\frac{\beta_{ZY}}{\beta_{ZX}} \text{ or } \beta'_{XY} = \rho_{ZY}^2\frac{\mathrm{Var}(Y)}{\mathrm{Var}(X)}\frac{\beta_{ZX}}{\beta_{ZY}}$$

By applying this general conclusion to the two confounding scenarios due to shared genetic effect (Figure S3), we can estimate the magnitudes of the genetically confounded associations.

**Scenario 1**: Genetically confounded association ($\beta_{XY}^c$) between a maternal phenotype ($X$) and a pregnancy outcome ($Y$) (Figure S3A). In this case, the confounder is set of the transmitted alleles ($Z$) that are associated with a maternal phenotype ($X$) and the genetically confounded association can be written as:

$$\beta_{XY}^c = \frac{h_X^2}{2}\frac{\beta_{FY}}{\beta_{MX}}$$

where $\rho_{ZX}^2$ is replaced by the half of the heritability of $X$ ($h_X^2$) and $\beta_{MX}$ and $\beta_{FY}$ are respectively the regression coefficient of genetic effects on a maternal phenotype ($X$) and a pregnancy outcome ($Y$). Assuming all the genetic variants (including those unidentified ones) associated with a maternal phenotype ($X$) have similar effects as those estimated from the genetic score built on the known GWA SNPs ($\hat{\beta}_{MX}$ and $\hat{\beta}_{FY}$ for maternal and fetal genetic effects), then we can estimate the magnitude of the genetically confounded association by:

$$\hat{\beta}_{XY}^{c} = \frac{h_X^2}{2}\frac{\hat{\beta}_{FY}}{\hat{\beta}_{MX}}$$

As the ratio of two normally distributed variables (i.e. $\hat{\beta}_{FY}$ and $\hat{\beta}_{MX}$), there is no simple analytical solution to obtain the distribution of this estimate ($\hat{\beta}_{XY}^{c}$). We calculated the standard error (se) using a normal approximation (the formula is similar to the ratio estimate above):

$$\mathrm{se}\left(\hat{\beta}_{XY}^{c}\right) = \frac{h_X^2}{2}\sqrt{\frac{\mathrm{se}\left(\hat{\beta}_{ZY}\right)^2}{\hat{\beta}_{ZX}{}^2} + \frac{\hat{\beta}_{ZY}{}^2\,\mathrm{se}\left(\hat{\beta}_{ZX}\right)^2}{\hat{\beta}_{ZX}{}^4}}$$

The heritability of a maternal phenotypes ($X$) is not always same as the corresponding adult phenotype. We therefore calculated the heritability of a maternal phenotype ($h_X^2$) from the heritability of the corresponding adult phenotype ($h^2$, Table S4) by:

$$h_X^2 = h^2\frac{R_{\mathrm{mat}}^2}{R_{\mathrm{rep}}^2}$$

where $R_{\mathrm{rep}}^2$ and $R_{\mathrm{mat}}^2$ are respectively the variance explained by the same genetic score in the adult phenotype (as reported by the reference GWA study).and in a maternal phenotype (as observed in this current study). The intuition behind this calculation is that we assume a same set of genetic variants influence both the adult phenotype and the maternal phenotype and their cumulative additive genetic effects on these traits are proportional to the cumulative additive genetic effects of the GWA SNPs (variance explained by the genetic score).

**Scenario 2**: Genetically confounded association ($\beta_{YX'}^{c}$) between a pregnancy outcome ($Y$) and an adult (late) phenotype ($X'$) in offspring (Figure S3B). In this case, the confounder includes the maternal as well as the paternal transmitted alleles that are associated with adult phenotype ($X'$). The maternal transmitted alleles can be associated with the pregnancy outcome ($Y$) by both maternal effect ($\beta_{MY}$) and fetal genetic effect ($\beta_{FY}$). The genetically confounded association between a pregnancy outcome ($Y$) and adult phenotype ($X'$) is given by:

$$\beta^{c}_{YX'} = h^2\left(\frac{\beta_{MY}}{2} + \beta_{FY}\right)/\beta_{FX'} \quad \text{(Method 1)}$$

Where $h^2$ is the heritability of the adult phenotype ($X'$), $\beta_{MY}$ and $\beta_{FY}$ are the are the regression coefficients of maternal effect and fetal genetic effect on a pregnancy outcome ($Y$) and $\beta_{FX'}$ is the regression coefficient of genetic effect on an adult phenotype ($X'$) in offspring. This estimate partitions the genetically confounded association into the maternal ($\frac{\beta_{MY}}{2}$) and the fetal component ($\beta_{FY}$).

Considering that the maternal non-transmitted alleles (h2) has no effect on the confounding, we can also use a simple alternative to estimate the genetically confounded association:

$$\beta^{c}_{YX'} = h^2\left(\frac{\beta_{\mathrm{h1}} + \beta_{\mathrm{h3}}}{2}\right)/\beta_{FX'} \quad \text{(Method 2)}$$

If the assumptions of additivity and zero parent of origin effect ($\beta_{\mathrm{h1}} = \beta_{\mathrm{h2}} + \beta_{\mathrm{h3}}$) hold, this formula is mathematically equivalent to the previous one. This simple formula uses the combination of two regression coefficients ($\beta_{\mathrm{h1}} + \beta_{\mathrm{h3}}$) and does not require the assumption of zero parent of origin effect. It is theoretically more accurate and precise than the previous one in statistical inference; however, it cannot dissect the maternal and fetal contributions of the confounding.

Although we did not directly estimate $\beta_{FX'}$ (because the adult phenotype is missing), the magnitude of the association should always equal to one ($\beta_{FX'} = 1$) if the reference GWA study (from which we extracted the GWA SNPs and effect size estimates to construct the genetic scores) is representative (i.e. homogenous) of our study population. Again, if assuming all the genetic variants associated with the adult phenotype ($X'$) have similar maternal and fetal genetic effects as those estimated from the genetic score built on known GWA SNPs, then we can estimate the genetically confounded association between a pregnancy outcome ($Y$) and an adult (late) phenotype ($X'$) in offspring by:

$$\hat{\beta}^{c}_{YX'} = h^2\frac{\mathrm{Var}(X')}{\mathrm{Var}(Y)}\left(\frac{\hat{\beta}_{MY}}{2} + \hat{\beta}_{FY}\right) \quad \text{(Method 1)}$$

or

$$\hat{\beta}^{c}_{YX'} = h^2\frac{\mathrm{Var}(X')}{\mathrm{Var}(Y)}\left(\frac{\widehat{\beta_{\mathrm{h1}} + \beta_{\mathrm{h3}}}}{2}\right) \quad \text{(Method 2)}$$

By assuming the variance of the adult phenotype ($\mathrm{Var}(X')$) and its heritability ($h^2$), and the variance of the pregnancy outcome ($\mathrm{Var}(Y)$) as known population parameters, the estimates ($\hat{\beta}^{c}_{YX'}$) follow normal distributions as with mean ($\frac{\hat{\beta}_{MY}}{2} + \hat{\beta}_{FY}$ or $\frac{\widehat{\beta_{\mathrm{h1}}+\beta_{\mathrm{h3}}}}{2}$) and se scaled by $h^2 \frac{\mathrm{Var}(X')}{\mathrm{Var}(Y)}$.

We compared our estimated genetically confounded association ($\beta^{c}_{YX'}$) between a pregnancy outcome ($Y$) and an adult (late) phenotype ($X'$) (referred as the **Confounded (eff)**) with:

1) The **confounded (cor)** phenotypic associations between birth weight and adult phenotypes based the genetic correlations reported by Horikoshi et al.[24] We estimated the phenotypic correlation using $R_{\mathrm{p}} = R_{\mathrm{g}}\sqrt{h_1^2 h_2^2}$, where $R_{\mathrm{g}}$ is the reported genetic correlation and $\sqrt{h_1^2 h_2^2}$ is the square root of the product of the heritability of an adult phenotype ($h_1^2$) and birth weight ($h_2^2$) (also see Table S4).

2) The **observed** phenotypic associations from the ALSPAC data set. We calculated the regression coefficients between birth weight and offspring height, BMI, BP and FPG measured at age 17 in a subset of ALPAC samples. Gestational age and sex were included as covariates.

3) The **reported** associations between birth weight and BP and T2D susceptibility from a recent epidemiological meta study[25]. We extracted the “unstratified risk” of T2D and changes in SBP and DBP per 1kg increase in birth weight from Table 1 of the reference[25] and covered the values to changes in standard deviation (of an adult phenotype) per 1SD (484g) increase in birth weight.

##### *Multivariable Mendelian Randomization analysis*

The analyses of associations between the haplotype genetic scores and birth outcomes provided a powerful way to illustrate the different maternal and fetal effects of the SNPs known to be associated with an intermediate phenotype. However, the genetic scores built on hundreds or thousands of SNPs were less specific, as they are more likely to be associated with other phenotypic traits, therefore introducing ambiguities in the interpretation of genetic score associations[26]. To circumvent this issue, we performed multivariable Mendelian Randomization (MR) analysis using the recently developed Mendelian randomization pleiotropy residual sum and outlier (MR-PRESSO) approach[27]. This method can detect and correct for SNPs with horizontal pleiotropic effects (outliers) in multivariable MR test. Using this approach, we tested whether the effects of the maternal transmitted (h1), maternal non-transmitted (h2) and paternal transmitted (h3) alleles of the GWA SNPs on a pregnancy outcome are proportional to their reported effects on an intermediate phenotype by large GWA studies. The allele-specific effect estimates (for the h1, h2 and h3 alleles) of a SNP were obtained using the same regression method as those in testing the associations between haplotype genetic scores and pregnancy outcomes ($Y \sim S_{\mathrm{h1}} + S_{\mathrm{h2}} + S_{\mathrm{h3}} + \mathrm{Cov}$), except for the haplotype genetic scores were replace by the haplotypes of a SNP.

In two-sample MR analysis[28], the non-zero slope of the regression line between the reported effect on an exposure and the estimated effect on an outcome has usually been interpreted as *causal effect*. In the current context, the allele-specific effects by the maternal transmitted (h1), maternal non-transmitted (h2) and paternal transmitted (h3) alleles on a pregnancy outcome were used to index the maternal effect (h2), fetal genetic effect (h3), or combination of both (h1). In this case, a non-zero "casual estimate"" (i.e. slope of the regression line between the reported effects of the SNPs on a maternal phenotype and the estimated allele-specific effect on a birth outcome) may or may not imply causal effect (Figure S4). Using the height GWA SNPs as an example, the "causal estimate" of maternal height on gestational duration by the maternal non-transmitted alleles (h2) represents a maternal effect, and it is very likely that the causal path is mediated by maternal height. However, the "estimated effect" on birth weight by the paternal transmitted alleles (h3) indicates a fetal genetic effect and the estimated effect sizes on birth weight are even proportional to their effects on adult height, which most probably reflects that the pleiotropic effects of the height associated SNPs on both birth weight and adult height act though a common pathway[29]. A significant MR-PRESSO global test indicates that the allele-specific effect sizes of the variants on an outcome are not always proportional to their effects on an intermediate phenotype. In cases of causal inferences (e.g. utilizing non-transmitted alleles to infer causal effect of a maternal trait on a birth outcome, Figure S4A), this is sign of horizontal pleiotropy – meaning the identified maternal effect may not exclusively be mediated by the intermediate maternal phenotype. In cases of genetic correlation between a birth outcome and an adult phenotype (e.g. the fetal effects of height associated SNPs on birth weight, Figure S4B), this suggests the genetic effects of the SNPs on a birth outcome are not always proportional to their reported effects on the adult phenotype, which might suggest involvement of different molecular pathways.

##### *Meta-analysis*

Fixed-effect meta-analysis was used to combine the results from different studies. Between-study heterogeneity was tested by Cochran's Q test.

### Supplementary Tables

#### Table S1. Data sets and number of samples

| Data set | Phenotype[a] | Genotype[b] | | Filtered[c] | | | |
|---|---|---|---|---|---|---|---|
| | Record | Typed | Passed | Record | Mother | Infant | Duo |
| FIN | 1644 | 2962 | 2710 | 1371 | 1322 | 1217 | 1170 |
| MoBa | 2035 | 3120 | 2979 | 1933 | 1804 | 1134 | 1009 |
| DNBC | 2068 | 3886 | 3799 | 2038 | 1912 | 1865 | 1739 |
| HAPO | 1507 | 2866 | 2743 | 1266 | 1203 | 1152 | 1089 |
| GPN | 1495 | 2852 | 1360 | 440 | 419 | 364 | 343 |
| ALSPAC | 15443 | 17842 | 17435 | 9806 | 7603 | 7587 | 5384 |
| Total | | | | 16854 | 14263 | 13319 | **10734** |

a: Number of phenotype records (pregnancies)

b: Number of genotyped subjects (mothers and infants) and subjects passed genotype QC

c: Number of phenotype records (pregnancies), and number of mothers, infants and mother/infant duos after applying genotype QC and phenotype filtering. 10,734 was the total number of mother/infant duos used in the analyses.

#### Table S2. Descriptive statistics of maternal traits and birth outcomes

| Trait | FIN | | | MoBa | | | DNBC | | | HAPO | | | GPN | | | ALSPAC | | | Total | | |
|---|---|---|---|---|---|---|---|---|---|---|---|---|---|---|---|---|---|---|---|---|---|
| | N | mean | sd | N | mean | sd | N | mean | sd | N | mean | sd | N | mean | sd | N | mean | sd | N | mean | sd |
| ***Maternal traits*** | | | | | | | | | | | | | | | | | | | | | |
| Height (cm) | 1170 | 166.7 | 5.8 | 975 | 167.9 | 5.8 | 1653 | 168.9 | 6.1 | 1089 | 164.5 | 6.2 | 342 | 164.9 | 7.8 | 4993 | 164.4 | 6.6 | 10222 | 165.7 | 6.4 |
| BMI (kg/m2) | 1168 | 23.0 | 3.7 | 958 | 24.0 | 4.2 | 1625 | 23.5 | 4.2 | 1051 | 24.2 | 4.7 | 337 | 24.2 | 5.3 | 4759 | 22.9 | 3.7 | 9898 | 23.3 | 4.0 |
| SBP (mmHg) | | | | | | | | | | 1089 | 108.0 | 9.7 | | | | 4923 | 113.1 | 6.8 | 6012 | 112.1 | 7.4 |
| DBP (mmHg) | | | | | | | | | | 1089 | 71.1 | 7.8 | | | | 4943 | 66.8 | 4.7 | 6032 | 67.6 | 5.4 |
| FPG (mmol/L) | | | | | | | | | | 1088 | 4.5 | 0.4 | | | | 2452 | 5.2 | 0.4 | 3540 | 5.0 | 0.4 |
| ***Birth outcomes*** | | | | | | | | | | | | | | | | | | | | | |
| Gestational days | 892 | 282.0 | 6.3 | 522 | 280.4 | 3.6 | 977 | 282.6 | 3.0 | 1034 | 281.2 | 7.2 | 197 | 276.4 | 4.2 | 4967 | 279.8 | 7.9 | 8589 | 280.5 | 7.0 |
| | 255 | 237.9 | 13.4 | 487 | 248.2 | 11.4 | 748 | 240.1 | 15.2 | | | | 146 | 212.0 | 20.3 | 212 | 242.7 | 13.5 | 1848 | 240.0 | 14.3 |
| Birth weight (g) | 892 | 3578.0 | 434.3 | | | | 973 | 3712.6 | 456.5 | 1034 | 3463.3 | 487.0 | 197 | 3505.2 | 366.0 | 4909 | 3514.3 | 455.7 | 8005 | 3538.7 | 455.5 |
| | 255 | 2348.8 | 477.1 | | | | 739 | 2459.5 | 645.8 | | | | 146 | 1641.4 | 522.1 | 208 | 2543.1 | 536.8 | 1348 | 2362.9 | 587.4 |
| Birth length (cm) | 892 | 50.4 | 1.9 | | | | | | | 1034 | 50.7 | 2.2 | 192 | 34.7 | 1.4 | 4004 | 51.0 | 2.1 | 6122 | 50.4 | 2.1 |
| | 253 | 45.0 | 2.9 | | | | | | | | | | 146 | 28.4 | 3.5 | 152 | 47.3 | 2.5 | 551 | 41.3 | 3.0 |
| ***Male/Female*** | 606/564 | | | 503/506 | | | 908/831 | | | 550/539 | | | 189/154 | | | 2690/2694 | | | | | |

a: In HAPO, maternal FPG was measured between 24~32 weeks of gestation. In ALSPAC, the FPG was measured 18 years after pregnancy.

b: Descriptive statistics of pregnancy outcomes were calculated in term (>=38 weeks) and preterm (<37 weeks) infants separately. All infants in the HAPO data set were term.

#### Table S3. Association between maternal phenotypes and birth outcomes

| Data set | Gestational days | | | Preterm birth (log(OR)) | | | Birth weight (g) | | | Birth length (cm) | | |
|---|---|---|---|---|---|---|---|---|---|---|---|---|
| | beta | se | p-val | beta | se | p-val | beta | se | p-val | beta | se | p-val |
| ***Height* (cm)** | | | | | | | | | | | | |
| ALSPAC | 0.13 | 0.025 | **3.2E-07** | -0.045 | 0.012 | **0.00012** | 16 | 0.91 | **1.7E-66** | 0.074 | 0.0048 | **2.2E-52** |
| FIN | 0.15 | 0.1 | 0.13 | -0.018 | 0.012 | 0.15 | 14 | 1.9 | **1.6E-13** | 0.065 | 0.0087 | **1.9E-13** |
| MoBa | 0.24 | 0.1 | **0.017** | -0.031 | 0.011 | **0.0069** | | na | | | na | |
| DNBC | 0.37 | 0.094 | **0.00011** | -0.029 | 0.0085 | **0.00053** | 15 | 1.7 | **4.7E-17** | | | |
| HAPO | 0.1 | 0.04 | **0.013** | | | | 18 | 2.2 | **5.0E-17** | 0.08 | 0.0098 | **7.6E-16** |
| GPN | 0.43 | 0.25 | 0.084 | -0.024 | 0.015 | 0.12 | 7.6 | 2.3 | **0.00089** | 0.011 | 0.012 | 0.37 |
| meta | 0.14 | 0.02 | **2.2E-12** | -0.03 | 0.005 | **2.2E-09** | 15 | 0.67 | **1.5E-111** | 0.068 | 0.0037 | **1.6E-75** |
| p_het | 0.097 | | | 0.6 | | | **0.0054** | | | **2.1E-05** | | |
| ***BMI* (kg/m²)** | | | | | | | | | | | | |
| ALSPAC | 0.099 | 0.045 | **0.029** | -0.042 | 0.023 | 0.065 | 20 | 1.6 | **2.9E-33** | 0.052 | 0.0085 | **8.6E-10** |
| FIN | -0.081 | 0.16 | 0.61 | 0.017 | 0.019 | 0.38 | 9.3 | 3 | **0.0018** | 0.036 | 0.014 | **0.0086** |
| MoBa | -0.12 | 0.14 | 0.38 | -0.0025 | 0.016 | 0.88 | | na | | | na | |
| DNBC | -0.012 | 0.14 | 0.93 | -0.0077 | 0.012 | 0.52 | 13 | 2.5 | **1.3E-07** | | na | |
| HAPO | 0.035 | 0.054 | 0.51 | | | | 18 | 2.9 | **8.7E-10** | 0.039 | 0.013 | **0.0024** |
| GPN | -0.25 | 0.36 | 0.48 | -0.0038 | 0.022 | 0.86 | 7.8 | 3.2 | **0.016** | 0.054 | 0.018 | **0.0026** |
| meta | 0.049 | 0.032 | 0.12 | -0.006 | 0.0075 | 0.42 | 16 | 1.1 | **1.0E-47** | 0.047 | 0.006 | **3.9E-15** |
| p_het | 0.5 | | | 0.41 | | | **0.0012** | | | 0.68 | | |
| ***SBP* (mmHg)** | | | | | | | | | | | | |
| ALSPAC | -0.074 | 0.021 | **0.00038** | | | | -4.1 | 0.98 | **2.9E-05** | -0.014 | 0.0052 | **0.0057** |
| HAPO | 0.0059 | 0.027 | 0.83 | | na | | -0.74 | 1.5 | 0.61 | 0.006 | 0.0066 | 0.36 |
| meta | -0.044 | 0.017 | **0.0073** | | | | -3 | 0.81 | **0.00018** | -0.0065 | 0.0041 | 0.11 |
| p_het | **0.019** | | | | | | 0.055 | | | **0.015** | | |
| ***DBP* (mmHg)** | | | | | | | | | | | | |
| ALSPAC | -0.16 | 0.031 | **1.6E-07** | | | | -7.4 | 1.4 | **2.9E-07** | -0.018 | 0.0075 | **0.02** |
| HAPO | -0.037 | 0.035 | 0.28 | | na | | -4.1 | 1.9 | **0.027** | 0.00075 | 0.0084 | 0.93 |
| meta | -0.11 | 0.023 | **3.3E-06** | | | | -6.2 | 1.1 | **6.0E-08** | -0.0094 | 0.0056 | 0.094 |
| p_het | **0.008** | | | | | | 0.16 | | | 0.11 | | |
| ***FPG* (mmol/L)[a]** | | | | | | | | | | | | |
| ALSPAC | 0.21 | 0.53 | 0.69 | -0.058 | 0.26 | 0.82 | 28 | 19 | 0.15 | -0.1 | 0.1 | 0.32 |
| HAPO | -0.28 | 0.73 | 0.7 | | na | | 192 | 39 | **5.7E-07** | 0.62 | 0.18 | **0.00048** |
| meta | 0.043 | 0.43 | 0.92 | -0.058 | 0.26 | 0.82 | 61 | 17 | **0.00042** | 0.082 | 0.089 | 0.36 |
| p_het | 0.59 | | | 1 | | | **0.00011** | | | **0.00043** | | |

a: In HAPO, maternal FPG was measured between 24~32 weeks of gestation. In ALSPAC, the FPG showed here was measured 18 years after pregnancy.

b: The shaded rows show the meta-analysis results. p_het: P value for heterogeneity test.

#### Table S4. GWA SNPs used to build the genetic scores

| Trait | Total[a] | Selected[b] | Used[c] | Unit[d] | Reference[e] | $H^2$ [f] |
|---|---|---|---|---|---|---|
| Height | 3290 | 2214 | 2130 | SD* | [14] | 0.80 [30] |
| BMI | 941 | 643 | 628 | SD* | [14] | 0.6 [31] |
| BP | 885 | 878 | 831 | mmHg | [32] | 0.5 [33] |
| FPG | 23 | 23 | 22 | mmol/L | [16] | 0.66 [34] |
| T2D | 403 | 369 | 306 | Log(OR) | [17] | 0.45 [35] |
| Birth weight | 106 | 96 | 86 | SD* | [36] | 0.3 [37] |

a: Total number of GWA SNPs with genome-wide significance (P<1E-8 (height, BMI) or P<5E-8 (BP, FPG and T2D).

b: Number of selected GWA significant SNPs (polymorphic in 1000 Genomes European samples and not in close LD with other more significant SNPs).

c: SNPs that were available in all study data sets were used to constructing the genetic scores.

d: Units of the effect size estimates (SD: standard deviation) in the original GWA reports. Units using SD* were transformed to cm (height), kg/m$^2$ (BMI) and g (Birth weight) according 1SD(height) = 6.4cm, 1SD (BMI) = 4.0kg/m$^2$, and 1SD(birth weight)=484g in constructing the genetic scores.

e: Reference GWA studies

f: Reported heritability of the traits.

#### Table S5. Associations between maternal genetic scores and maternal traits

| Maternal trait | Maternal genotype (h1+h2) | | | | Maternal trans (h1) | | | | Maternal non-trans (h2) | | | |
|---|---|---|---|---|---|---|---|---|---|---|---|---|
| | beta | se | p-val | r2 | beta | se | p-val | r2 | beta | se | p-val | r2 |
| Height[a] (cm) | 0.8 | 0.014 | **<2.2E-308** | 0.25 | 0.81 | 0.02 | **<2.2E-308** | 0.14 | 0.8 | 0.02 | **<2.2E-308** | 0.14 |
| BMI[a] (kg/$m^2$) | 0.87 | 0.037 | **1.30E-120** | 0.052 | 0.88 | 0.052 | **1.70E-63** | 0.028 | 0.85 | 0.053 | **2.80E-59** | 0.026 |
| SBP[b] (mmHg) | 0.27 | 0.025 | **4.50E-27** | 0.021 | 0.26 | 0.035 | **4.20E-14** | 0.01 | 0.27 | 0.035 | **1.60E-14** | 0.011 |
| DBP[b] (mmHg) | 0.32 | 0.028 | **2.80E-30** | 0.024 | 0.32 | 0.04 | **8.50E-16** | 0.012 | 0.32 | 0.04 | **6.20E-16** | 0.012 |
| FPG[c] (mmol/L) (HAPO) | 1 | 0.1 | **1.20E-23** | 0.083 | 1.1 | 0.14 | **2.20E-13** | 0.043 | 1 | 0.14 | **1.20E-12** | 0.04 |
| FPG[c] (mmol/L) (ALSPAC) | 0.89 | 0.088 | **9.60E-24** | 0.041 | 0.85 | 0.12 | **8.20E-12** | 0.018 | 0.94 | 0.12 | **5.00E-14** | 0.023 |
| FPG[d] (mmol/L) (HAPO) | 0.047 | 0.018 | **0.0077** | 0.0079 | 0.0067 | 0.025 | 0.79 | 0.00023 | 0.088 | 0.025 | **0.00054** | 0.012 |
| FPG[d] (mmol/L) (ALSPAC) | 0.057 | 0.016 | **0.00027** | 0.0065 | 0.046 | 0.022 | **0.034** | 0.0025 | 0.069 | 0.022 | **0.0023** | 0.0041 |

a: Associations between maternal height, BMI and their corresponding maternal genotype or haplotypes scores (based on the meta results of the six data set, except birth length was not available in the DNBC data set).

b: Associations between maternal systolic and diastolic blood pressure measures and their corresponding maternal genotype or haplotypes scores based on ALSPAC and HAPO. There were no preterm birth sample in HAPO and in ALSPAC samples with blood pressure data.

c: Associations between fasting plasma glucose (FPG) genetic scores and maternal FPG levels either measured during (HAPO) or 18 years after pregnancy (ALSPAC).

d: Associations between type 2 diabetes (T2D) genetic scores and maternal FPG levels either measured during (HAPO) or 18 years after pregnancy (ALSPAC).

#### Table S6. Associations between maternal height genetic scores and maternal height

| Data set | geno (h1+h2) | | | | trans (h1) | | | | non-trans (h2) | | | |
|---|---|---|---|---|---|---|---|---|---|---|---|---|
| | beta | se | p-val | r2 | beta | se | p-val | r2 | beta | se | p-val | r2 |
| ALSPAC | 0.82 | 0.021 | **1.70E-295** | 0.24 | 0.83 | 0.03 | **2.40E-160** | 0.12 | 0.82 | 0.03 | **3.60E-152** | 0.12 |
| FIN | 0.77 | 0.037 | **2.40E-80** | 0.27 | 0.78 | 0.055 | **2.50E-42** | 0.14 | 0.76 | 0.054 | **2.10E-41** | 0.13 |
| MoBa | 0.72 | 0.041 | **5.50E-62** | 0.25 | 0.72 | 0.059 | **1.20E-32** | 0.13 | 0.73 | 0.059 | **4.10E-32** | 0.12 |
| DNBC | 0.77 | 0.033 | **7.90E-106** | 0.25 | 0.75 | 0.048 | **2.90E-52** | 0.12 | 0.79 | 0.048 | **5.10E-57** | 0.13 |
| HAPO | 0.87 | 0.039 | **7.80E-91** | 0.32 | 0.87 | 0.057 | **6.60E-48** | 0.16 | 0.87 | 0.057 | **3.80E-47** | 0.16 |
| GPN | 0.92 | 0.098 | **1.20E-18** | 0.21 | 1 | 0.14 | **1.60E-12** | 0.12 | 0.83 | 0.13 | **1.00E-09** | 0.089 |
| meta[a] | 0.8 | 0.014 | **<2.2E-308** | 0.25 | 0.81 | 0.02 | **<2.2E-308** | 0.14 | 0.8 | 0.02 | **<2.2E-308** | 0.14 |
| p_het | 0.057 | | | | 0.16 | | | | 0.53 | | | |

a: the meta-analysis results. p_het: P value for heterogeneity test.

#### Table S7. Associations between maternal BMI genetic scores and maternal BMI

| Data set | geno (h1+h2) | | | | trans (h1) | | | | non-trans (h2) | | | |
|---|---|---|---|---|---|---|---|---|---|---|---|---|
| | beta | se | p-val | r2 | beta | se | p-val | r2 | beta | se | p-val | r2 |
| ALSPAC | 0.86 | 0.05 | **5.20E-65** | 0.06 | 0.79 | 0.071 | **2.10E-28** | 0.025 | 0.94 | 0.071 | **2.80E-39** | 0.035 |
| FIN | 0.68 | 0.1 | **7.50E-11** | 0.036 | 0.73 | 0.14 | **2.50E-07** | 0.021 | 0.63 | 0.15 | **2.10E-05** | 0.015 |
| MoBa | 0.95 | 0.13 | **2.20E-13** | 0.054 | 0.95 | 0.18 | **1.30E-07** | 0.028 | 0.94 | 0.18 | **3.20E-07** | 0.026 |
| DNBC | 0.91 | 0.099 | **1.40E-19** | 0.049 | 1.1 | 0.14 | **2.20E-15** | 0.037 | 0.71 | 0.14 | **2.00E-07** | 0.015 |
| HAPO | 1 | 0.14 | **6.40E-13** | 0.047 | 1.2 | 0.2 | **5.90E-09** | 0.032 | 0.84 | 0.2 | **2.40E-05** | 0.017 |
| GPN | 1.2 | 0.29 | **6.70E-05** | 0.047 | 1.5 | 0.38 | **8.50E-05** | 0.044 | 0.75 | 0.4 | 0.06 | 0.0086 |
| meta[a] | 0.87 | 0.037 | **1.30E-120** | 0.052 | 0.88 | 0.052 | **1.70E-63** | 0.028 | 0.85 | 0.053 | **2.80E-59** | 0.026 |
| p_het | 0.35 | | | | 0.064 | | | | 0.41 | | | |

a: the meta-analysis results. p_het: P value for heterogeneity test.

#### Table S8. Associations between maternal blood pressure genetic scores and maternal blood pressure (SBP and DBP)

| Data set[a] | geno (h1+h2) | | | | trans (h1) | | | | non-trans (h2) | | | |
|---|---|---|---|---|---|---|---|---|---|---|---|---|
| | beta | se | p-val | r2 | beta | se | p-val | r2 | beta | se | p-val | r2 |
| ***SBP*** | | | | | | | | | | | | |
| ALSPAC | 0.25 | 0.026 | **5.20E-22** | 0.018 | 0.24 | 0.037 | **5.20E-11** | 0.0093 | 0.26 | 0.037 | **1.50E-12** | 0.0086 |
| HAPO | 0.39 | 0.078 | **6.80E-07** | 0.022 | 0.45 | 0.11 | **3.80E-05** | 0.015 | 0.33 | 0.11 | **0.0028** | 0.0074 |
| meta | 0.27 | 0.025 | **4.50E-27** | 0.021 | 0.26 | 0.035 | **4.20E-14** | 0.01 | 0.27 | 0.035 | **1.60E-14** | 0.011 |
| p_het | 0.089 | | | | 0.067 | | | | 0.56 | | | |
| ***DBP*** | | | | | | | | | | | | |
| ALSPAC | 0.32 | 0.029 | **1.50E-26** | 0.02 | 0.31 | 0.042 | **3.90E-13** | 0.01 | 0.33 | 0.042 | **6.10E-15** | 0.0097 |
| HAPO | 0.41 | 0.1 | **5.60E-05** | 0.013 | 0.54 | 0.14 | **0.00017** | 0.011 | 0.28 | 0.14 | **0.042** | 0.0037 |
| meta[b] | 0.32 | 0.028 | **2.80E-30** | 0.024 | 0.32 | 0.04 | **8.50E-16** | 0.012 | 0.32 | 0.04 | **6.20E-16** | 0.012 |
| p_het | 0.38 | | | | 0.11 | | | | 0.77 | | | |

a: Blood pressure data were only available in ALSPAC and HAPO.

b: the meta-analysis results. p_het: P value for heterogeneity test.

#### Table S9. Associations between maternal FPG and T2D genetic scores and maternal FPG

| Data set[a] | geno (h1+h2) | | | | trans (h1) | | | | non-trans (h2) | | | |
|---|---|---|---|---|---|---|---|---|---|---|---|---|
| | beta | se | p-val | r2 | beta | se | p-val | r2 | beta | se | p-val | r2 |
| ***FPG***[b] | | | | | | | | | | | | |
| ALSPAC | 0.89 | 0.088 | **9.60E-24** | 0.041 | 0.85 | 0.12 | **8.20E-12** | 0.018 | 0.94 | 0.12 | **5.00E-14** | 0.023 |
| HAPO | 1 | 0.1 | **1.20E-23** | 0.083 | 1.1 | 0.14 | **2.20E-13** | 0.043 | 1 | 0.14 | **1.20E-12** | 0.04 |
| meta[d] | 0.95 | 0.066 | **4.60E-47** | 0.059 | 0.94 | 0.093 | **8.40E-24** | 0.03 | 0.97 | 0.092 | **1.50E-25** | 0.032 |
| p_het | 0.3 | | | | 0.26 | | | | 0.72 | | | |
| ***T2D***[c] | | | | | | | | | | | | |
| ALSPAC | 0.057 | 0.016 | **0.00027** | 0.0065 | 0.046 | 0.022 | **0.034** | 0.0025 | 0.069 | 0.022 | **0.0023** | 0.0041 |
| HAPO | 0.047 | 0.018 | **0.0077** | 0.0079 | 0.0067 | 0.025 | 0.79 | 0.00023 | 0.088 | 0.025 | **0.00054** | 0.012 |
| meta[d] | 0.053 | 0.012 | **6.70E-06** | 0.0061 | 0.029 | 0.016 | 0.074 | 0.00097 | 0.077 | 0.017 | **4.50E-06** | 0.0063 |
| p_het | 0.68 | | | | 0.24 | | | | 0.57 | | | |

a: Maternal FPG during pregnancy was only available in HAPO. FPG was measure in a subset of ALSPAC mothers 17 years after the pregnancy.

b: Associations between fasting plasma glucose (FPG) genetic scores (based on 22 FPG SNPs) and maternal FPG levels either measured during (HAPO) or 18 years after pregnancy (ALSPAC).

c: Associations between type 2 diabetes (T2D) genetic scores (based on 306 T2D SNPS) and maternal FPG levels either measured during (HAPO) or 18 years after pregnancy (ALSPAC).

d: the meta-analysis results. p_het: P value for heterogeneity test.

**Table S10. Correlation (p-value) between maternal genotype score ($S_{mat}$), fetal genotype score ($S_{fet}$); and the three haplotype genetic scores: $S_{h1}$, $S_{h2}$ and $S_{h3}$.**

**A. Height**

| score | $S_{mat}$ | $S_{fet}$ | $S_{h1}$ | $S_{h2}$ |
|---|---|---|---|---|
| $S_{fet}$ | 0.53 (**<2.2E-308**) | | | |
| $S_{h1}$ | 0.72 (**<2.2E-308**) | 0.71 (**<2.2E-308**) | | |
| $S_{h2}$ | 0.72 (**<2.2E-308**) | **0.058** (1.2e-08) | **0.042** (0.0008) | |
| $S_{h3}$ | **0.044** (3.1e-06) | 0.72 (**<2.2E-308**) | **0.019** (0.024) | **0.044** (2.7e-06) |

**B. BMI**

| score | $S_{mat}$ | $S_{fet}$ | $S_{h1}$ | $S_{h2}$ |
|---|---|---|---|---|
| $S_{fet}$ | 0.5 (**<2.2E-308**) | | | |
| $S_{h1}$ | 0.71 (**<2.2E-308**) | 0.7 (**<2.2E-308**) | | |
| $S_{h2}$ | 0.7 (**<2.2E-308**) | 0.0072 (0.23) | -0.0024 (0.41) | |
| $S_{h3}$ | 0.01 (0.15) | 0.71 (**<2.2E-308**) | 0.0018 (0.45) | 0.011 (0.14) |

**C. BP[a]**

| score | $S_{mat}$ | $S_{fet}$ | $S_{h1}$ | $S_{h2}$ |
|---|---|---|---|---|
| $S_{fet}$ | 0.5 (**<2.2E-308**) | | | |
| $S_{h1}$ | 0.71 (**<2.2E-308**) | 0.7 (**<2.2E-308**) | | |
| $S_{h2}$ | 0.7 (**<2.2E-308**) | -0.0034 (0.36) | -0.0067 (0.27) | |
| $S_{h3}$ | -0.0094 (0.22) | 0.7 (**<2.2E-308**) | -0.01 (0.14) | -0.00067 (0.48) |

**D. FPG**

| score | $S_{mat}$ | $S_{fet}$ | $S_{h1}$ | $S_{h2}$ |
|---|---|---|---|---|
| $S_{fet}$ | 0.49 (**<2.2E-308**) | | | |
| $S_{h1}$ | 0.7 (**<2.2E-308**) | 0.71 (**<2.2E-308**) | | |
| $S_{h2}$ | 0.7 (**<2.2E-308**) | -0.015 (0.063) | -0.012 (0.11) | |
| $S_{h3}$ | 0.013 (0.18) | 0.72 (**<2.2E-308**) | **0.031** (0.046) | -0.0096 (0.16) |

**E. T2D**

| score | $S_{mat}$ | $S_{fet}$ | $S_{h1}$ | $S_{h2}$ |
|---|---|---|---|---|
| $S_{fet}$ | 0.5 (**1.5e-223**) | | | |
| $S_{h1}$ | 0.7 (**<2.2E-308**) | 0.71 (**<2.2E-308**) | | |
| $S_{h2}$ | 0.71 (**<2.2E-308**) | -0.0089 (0.29) | 0.002 (0.42) | |
| $S_{h3}$ | -0.004 (0.37) | 0.71 (**<2.2E-308**) | 0.0074 (0.22) | -0.013 (0.18) |

**F. Birth weight**

| score | $S_{mat}$ | $S_{fet}$ | $S_{h1}$ | $S_{h2}$ |
|---|---|---|---|---|
| $S_{fet}$ | 0.5 (**<2.2E-308**) | | | |
| $S_{h1}$ | 0.71 (**<2.2E-308**) | 0.71 (**<2.2E-308**) | | |
| $S_{h2}$ | 0.7 (**<2.2E-308**) | 0.0019 (0.45) | 0.00047 (0.49) | |
| $S_{h3}$ | 0.0042 (0.36) | 0.71 (**<2.2E-308**) | 0.004 (0.34) | 0.0029 (0.42) |

a: BP score is the average of SBP and DBP scores.

#### Table S11. Phenotypic associations, estimated maternal causal effects and genetically confounded associations (per 1SD changes in maternal traits) between maternal traits and birth outcomes

| Maternal trait (1SD)[a] | Gestational days | | | Preterm birth (log(OR)) | | | Birth weight (g) | | | Birth length (cm) | | |
|---|---|---|---|---|---|---|---|---|---|---|---|---|
| | beta | se | p-val | beta | se | p-val | beta | se | p-val | beta | se | p-val |
| **Height (1SD=6.4cm)** | | | | | | | | | | | | |
| *Phenotypic* | 0.898 | 0.128 | **2.20E-12** | -0.193 | 0.0322 | **2.20E-09** | 96.2 | 4.29 | **1.50E-111** | 0.433 | 0.0235 | **1.60E-75** |
| *Causal (TSLS)* | 1.29 | 0.351 | **0.00024** | -0.298 | 0.0904 | **0.00099** | 57.8 | 12.3 | **2.40E-06** | 0.207 | 0.0671 | **0.002** |
| *Causal (Ratio)* | 1.01 | 0.323 | **0.0018** | -0.348 | 0.0787 | **9.60E-06** | 50.3 | 11.1 | **5.60E-06** | 0.178 | 0.0607 | **0.0033** |
| *Fetal genetic* | -0.711 | 0.326 | **0.029** | 0.13 | 0.0794 | 0.1 | 111 | 11.3 | **5.00E-23** | 0.59 | 0.0617 | **1.00E-21** |
| *Confounded* | -0.289 | 0.133 | **0.029** | 0.0528 | 0.0323 | 0.1 | 45.3 | 4.59 | **5.00E-23** | 0.24 | 0.0251 | **1.00E-21** |
| *Combined* | 0.722 | 0.349 | **0.039** | -0.296 | 0.0851 | **0.00051** | 95.6 | 12 | **1.50E-15** | 0.418 | 0.0657 | **1.90E-10** |
| **BMI (1SD=4.0kg/m$^2$)** | | | | | | | | | | | | |
| *Phenotypic* | 0.196 | 0.127 | 0.12 | -0.0241 | 0.03 | 0.42 | 62.4 | 4.3 | **1.00E-47** | 0.187 | 0.0238 | **3.90E-15** |
| *Causal (TSLS)* | -0.222 | 0.791 | 0.78 | 0.0735 | 0.219 | 0.74 | 59.4 | 28.4 | **0.036** | 0.339 | 0.152 | **0.026** |
| *Causal (Ratio)* | -0.452 | 0.686 | 0.51 | -0.222 | 0.165 | 0.18 | 87.7 | 23.5 | **0.00019** | 0.275 | 0.128 | **0.032** |
| *Fetal genetic* | 0.197 | 0.686 | 0.77 | -0.0845 | 0.163 | 0.6 | 18.2 | 23 | 0.43 | 0.0356 | 0.127 | 0.78 |
| *Confounded* | 0.0511 | 0.178 | 0.77 | -0.022 | 0.0425 | 0.6 | 4.74 | 5.97 | 0.43 | 0.00925 | 0.033 | 0.78 |
| *Combined* | -0.401 | 0.708 | 0.57 | -0.244 | 0.17 | 0.15 | 92.4 | 24.2 | **0.00014** | 0.284 | 0.132 | **0.032** |
| **BP (1SD=5.8mmHg)** | | | | | | | | | | | | |
| *Phenotypic* | -0.487 | 0.125 | **9.20E-05** | | | | -30.2 | 6.11 | **8.00E-07** | -0.0549 | 0.0306 | 0.072 |
| *Causal (TSLS)* | 1.71 | 1.23 | 0.16 | | | | -73.9 | 57.3 | 0.2 | -0.18 | 0.313 | 0.56 |
| *Causal (Ratio)* | -2.28 | 1.09 | **0.036** | 0.73 | 0.264 | **0.0057** | -39.2 | 36 | 0.28 | -0.186 | 0.199 | 0.35 |
| *Fetal genetic* | -1.93 | 1.1 | 0.078 | -0.0668 | 0.259 | 0.8 | -93.5 | 36.8 | **0.011** | -0.166 | 0.199 | 0.41 |
| *Confounded* | -0.228 | 0.129 | 0.078 | -0.0079 | 0.0306 | 0.8 | -11.1 | 4.35 | **0.011** | -0.0196 | 0.0235 | 0.41 |
| *Combined* | -2.51 | 1.09 | **0.022** | 0.722 | 0.266 | **0.0066** | -50.2 | 36.2 | 0.17 | -0.206 | 0.201 | 0.31 |
| **FPG (1SD=0.36mmol/L)** | | | | | | | | | | | | |
| *Phenotypic* | -0.102 | 0.264 | 0.7 | | | | 70.4 | 14 | **4.90E-07** | 0.223 | 0.0636 | **0.00046** |
| *Causal (TSLS)* | -0.731 | 1.26 | 0.56 | | | | 147 | 67.5 | **0.03** | 0.672 | 0.309 | **0.03** |
| *Causal (Ratio)* | -1.73 | 0.561 | **0.002** | 0.269 | 0.133 | **0.043** | 58.7 | 18.8 | **0.0018** | 0.13 | 0.0992 | 0.19 |
| *Fetal genetic* | 0.346 | 0.529 | 0.51 | -0.0482 | 0.127 | 0.7 | -53.8 | 18.3 | **0.0033** | -0.161 | 0.0977 | 0.1 |
| *Confounded* | 0.0865 | 0.132 | 0.51 | -0.0121 | 0.0317 | 0.7 | -13.5 | 4.58 | **0.0033** | -0.0402 | 0.0244 | 0.1 |
| *Combined* | -1.64 | 0.577 | **0.0043** | 0.257 | 0.137 | 0.06 | 45.3 | 19.4 | **0.019** | 0.0894 | 0.102 | 0.38 |

a: "*Combined*" is the sum of maternal "*Causal (Ratio)*" and genetically "*Confounded*" effect.

#### Table S12. Estimated changes in adult phenotypes per 1SD changes in birth outcomes

| Adult phenotype | Gestational days (1SD=11.4day) | | | Preterm birth (yes/no) | | | Birth weight (1SD=426g) | | | Birth length (1SD=2.2cm) | | |
|---|---|---|---|---|---|---|---|---|---|---|---|---|
| | beta | se | p-val | beta | se | p-val | beta | se | p-val | beta | se | p-val |
| **Height (cm)** | | | | | | | | | | | | |
| *Method 1* | -0.074 | 0.131 | 0.57 | -0.114 | 0.228 | 0.62 | 1.31 | 0.119 | **3.40E-28** | 1.25 | 0.125 | **1.50E-23** |
| *Method 2* | -0.00658 | 0.0926 | 0.94 | -0.043 | 0.162 | 0.79 | 1.32 | 0.0848 | **1.10E-54** | 1.26 | 0.0887 | **1.50E-45** |
| **BMI (kg/m²)** | | | | | | | | | | | | |
| *Method 1* | -0.0054 | 0.14 | 0.97 | -0.257 | 0.24 | 0.28 | 0.304 | 0.126 | **0.016** | 0.162 | 0.133 | 0.22 |
| *Method 2* | -0.0106 | 0.101 | 0.92 | -0.0578 | 0.175 | 0.74 | 0.186 | 0.0914 | **0.042** | 0.201 | 0.0961 | **0.037** |
| **BP (mmHg)** | | | | | | | | | | | | |
| *Method 1* | -0.331 | 0.131 | **0.011** | 0.231 | 0.224 | 0.3 | -0.327 | 0.116 | **0.0049** | -0.143 | 0.123 | 0.24 |
| *Method 2* | -0.249 | 0.0956 | **0.0093** | 0.305 | 0.164 | 0.063 | -0.359 | 0.085 | **2.50E-05** | -0.191 | 0.0902 | **0.035** |
| **FPG (mmol/L)** | | | | | | | | | | | | |
| *Method 1* | -0.00892 | 0.0102 | 0.38 | 0.0106 | 0.0176 | 0.54 | -0.0112 | 0.00906 | 0.21 | -0.00842 | 0.00951 | 0.38 |
| *Method 2* | -0.00395 | 0.00744 | 0.6 | 0.00605 | 0.0128 | 0.64 | -0.00278 | 0.00664 | 0.68 | -0.00304 | 0.00699 | 0.66 |
| **T2D (log(OR))** | | | | | | | | | | | | |
| *Method 1* | 0.0263 | 0.0385 | 0.49 | -0.0348 | 0.0648 | 0.59 | -0.0917 | 0.0342 | **0.0073** | -0.0511 | 0.036 | 0.16 |
| *Method 2* | 0.055 | 0.028 | **0.049** | -0.056 | 0.048 | 0.24 | -0.0723 | 0.0252 | **0.004** | -0.0334 | 0.0265 | 0.21 |

The genetically confounded association was estimated by two methods (Supplementary methods):

$$\hat{\beta}^{c}_{YX'} = h^2 \frac{\mathrm{Var}(X')}{\mathrm{Var}(Y)}\left(\frac{\hat{\beta}_{MY}}{2} + \hat{\beta}_{FY}\right) \text{ (Method 1) and } \hat{\beta}^{c}_{YX'} = h^2 \frac{\mathrm{Var}(X')}{\mathrm{Var}(Y)}\left(\frac{\widehat{\beta_{h1}+\beta_{h3}}}{2}\right) \text{ (Method 2)}$$

#### Table S13. Association between fetal birth weight genetic score and gestational age adjusted birth weight (gram)

| Data set | geno (h1+h2) | | | | maternal-trans (h1) | | | | paternal-trans (h3) | | | | diff (h1-h2) | | | |
|---|---|---|---|---|---|---|---|---|---|---|---|---|---|---|---|---|
| | beta[a] | se | p-val | r2 | beta | se | p-val | r2 | beta | se | p-val | r2 | beta | se | p-val | chi |
| ALSPAC | 1.2 | 0.085 | **3.90E-41** | 0.037 | 0.85 | 0.12 | **3.40E-12** | 0.011 | 1.4 | 0.12 | **3.10E-33** | 0.028 | -0.59 | 0.17 | **0.00055** | 12 |
| FIN | 0.58 | 0.16 | **0.00025** | 0.012 | 0.35 | 0.22 | 0.1 | 0.0029 | 0.84 | 0.23 | **0.00031** | 0.01 | -0.48 | 0.32 | 0.13 | 2.3 |
| DNBC | 0.9 | 0.16 | **7.30E-09** | 0.021 | 0.83 | 0.21 | **0.00012** | 0.0089 | 0.98 | 0.22 | **8.90E-06** | 0.012 | -0.15 | 0.3 | 0.62 | 0.25 |
| HAPO | 1.3 | 0.19 | **6.10E-11** | 0.038 | 0.9 | 0.27 | **0.00091** | 0.012 | 1.6 | 0.27 | **4.40E-09** | 0.028 | -0.72 | 0.39 | 0.064 | 3.4 |
| GPN | 1.1 | 0.23 | **3.00E-06** | 0.069 | 1.1 | 0.35 | **0.0016** | 0.033 | 1.1 | 0.3 | **0.00057** | 0.036 | 0.047 | 0.46 | 0.92 | 0.01 |
| meta[a] | 1 | 0.061 | **5.60E-64** | 0.031 | 0.79 | 0.087 | **1.20E-19** | 0.0092 | 1.3 | 0.086 | **3.90E-49** | 0.024 | -0.47 | 0.12 | **0.00013** | 0.0017 |
| p_het | **0.015** | | | | 0.26 | | | | **0.049** | | | | 0.51 | | | |

a: shows the meta-analysis results. p_het: P value for heterogeneity test.

b: tests the difference in effect size between the maternal transmitted (h1) and the paternal transmitted (h3) haplotype scores.

#### Table S14. MR-PRESSO: effects of maternal traits on birth outcomes

| | | Gestational days | | | Preterm birth | | | Birth weight | | | Birth length | | |
|---|---|---|---|---|---|---|---|---|---|---|---|---|---|
| Trait | Method | beta | se | p-val | beta | se | p-val | beta | se | p-val | beta | se | p-val |
| ***Height*** | | | | | | | | | | | | | |
| h1 | raw | 0.038 | 0.056 | 0.5 | -0.030 | 0.015 | 0.05 | 28.13 | 1.88 | **7.90E-44** | 0.133 | 0.010 | **2.90E-35** |
| | corrected | 0.078 | 0.055 | 0.16 | -0.027 | 0.015 | 0.077 | 26.56 | 1.88 | **7.60E-45** | 0.131 | 0.010 | **7.00E-36** |
| h2 | raw | 0.203 | 0.056 | **0.00029** | -0.056 | 0.015 | **0.00016** | 9.84 | 1.88 | **2.90E-07** | 0.033 | 0.010 | **0.0017** |
| | corrected | 0.203 | 0.055 | **0.00021** | -0.058 | 0.015 | **9.80E-05** | 10.00 | 1.88 | **1.00E-07** | 0.034 | 0.010 | **0.0006** |
| h3 | raw | -0.067 | 0.056 | 0.23 | 0.030 | 0.015 | **0.044** | 18.75 | 1.88 | **2.00E-20** | 0.103 | 0.011 | **6.30E-22** |
| | corrected | -0.064 | 0.055 | 0.24 | 0.028 | 0.015 | 0.055 | 18.75 | 1.88 | **1.30E-21** | 0.102 | 0.010 | **4.90E-22** |
| ***BMI*** | | | | | | | | | | | | | |
| h1 | raw | -0.075 | 0.1875 | 0.69 | -0.0625 | 0.05 | 0.2 | 24.25 | 6.75 | **0.00038** | 0.08 | 0.035 | **0.025** |
| | corrected | -0.045 | 0.18 | 0.81 | -0.0575 | 0.05 | 0.23 | 23 | 6.5 | **0.00045** | 0.0575 | 0.035 | 0.097 |
| h2 | raw | -0.1 | 0.1875 | 0.6 | 0.055 | 0.05 | 0.27 | 12.25 | 6.5 | 0.056 | 0.0825 | 0.035 | **0.02** |
| | corrected | -0.09 | 0.18 | 0.61 | 0.0575 | 0.0475 | 0.23 | 10 | 6.25 | 0.11 | 0.095 | 0.035 | **0.0053** |
| h3 | raw | 0.16 | 0.1925 | 0.4 | 0.01025 | 0.0525 | 0.84 | -4 | 6.5 | 0.54 | 0.0475 | 0.035 | 0.19 |
| | corrected | 0.22 | 0.1825 | 0.23 | 0.01425 | 0.05 | 0.77 | -3.5 | 6 | 0.55 | 0.0425 | 0.0325 | 0.21 |
| ***BP*** | | | | | | | | | | | | | |
| h1 | raw | -0.25 | 0.073 | **0.00078** | 0.056 | 0.02 | **0.0045** | -7.5 | 2.4 | **0.002** | -0.028 | 0.013 | **0.036** |
| | corrected | -0.22 | 0.069 | **0.0015** | 0.055 | 0.019 | **0.0044** | -7.5 | 2.3 | **0.0012** | -0.03 | 0.013 | **0.018** |
| h2 | raw | -0.03 | 0.07 | 0.67 | 0.058 | 0.019 | **0.0025** | -3.1 | 2.4 | 0.19 | -0.0093 | 0.013 | 0.48 |
| | corrected | -0.029 | 0.068 | 0.67 | 0.057 | 0.019 | **0.0028** | -3.6 | 2.3 | 0.12 | -0.014 | 0.013 | 0.27 |
| h3 | raw | -0.029 | 0.07 | 0.68 | 0.016 | 0.018 | 0.38 | -6.9 | 2.4 | **0.0044** | -0.016 | 0.013 | 0.2 |
| | corrected | -0.058 | 0.068 | 0.4 | 0.015 | 0.018 | 0.39 | -8.3 | 2.3 | **0.00035** | -0.012 | 0.013 | 0.33 |
| ***FPG*** | | | | | | | | | | | | | |
| h1 | raw | -3.4 | 2.1 | 0.11 | 0.19 | 0.67 | 0.78 | 14 | 62 | 0.82 | -0.11 | 0.3 | 0.72 |
| | corrected | -2.7 | 1.9 | 0.17 | 0.15 | 0.51 | 0.78 | -12 | 53 | 0.82 | | | |
| h2 | raw | -2.9 | 2.2 | 0.2 | 0.59 | 0.52 | 0.27 | 270 | 74 | **0.0018** | 0.54 | 0.27 | 0.056 |
| | corrected | -2.8 | 1.4 | 0.058 | NA | NA | NA | 250 | 69 | **0.0016** | NA | NA | NA |
| h3 | raw | 3.8 | 1.7 | **0.037** | -0.5 | 0.52 | 0.34 | -74 | 77 | 0.35 | -0.07 | 0.38 | 0.86 |
| | corrected | NA | NA | NA | NA | NA | NA | -43 | 59 | 0.48 | 0.039 | 0.35 | 0.91 |
| ***T2D*** | | | | | | | | | | | | | |
| h1 | raw | 0.02 | 0.30 | 0.96 | -0.009 | 0.074 | 0.9 | -11.7 | 11.3 | 0.3 | -0.069 | 0.056 | 0.2 |
| | corrected | NA | NA | NA | NA | NA | NA | -13.5 | 11.3 | 0.22 | NA | NA | NA |
| h2 | raw | 0.05 | 0.31 | 0.88 | 0.014 | 0.078 | 0.85 | 33.9 | 10.4 | **0.0014** | 0.039 | 0.052 | 0.46 |
| | corrected | NA | NA | NA | NA | NA | NA | 33.4 | 10.4 | **0.0012** | NA | NA | NA |
| h3 | raw | 0.83 | 0.32 | **0.011** | -0.104 | 0.078 | 0.17 | -28.2 | 10.0 | **0.0064** | -0.048 | 0.052 | 0.35 |
| | corrected | NA | NA | NA | NA | NA | NA | NA | NA | NA | NA | NA | NA |

a: The raw estimates were computed based on all GWA SNPs using standard IVW analysis. The corrected estimates were calculated after excluding outliers only when the MR-PRESSO global test suggested horizontal pleiotropy ($P < 0.05$).

b: The beta calculated based on h1, h2 or h3 may not reflect real causal effect. It indicates the slope of the regression line between the reported effects of the SNPs on an adult phenotype and the estimated allele-specific effects on a pregnancy outcome, i.e. it evaluates the allele-specific genetic effect of one-unit change in a maternal (adult) phenotype corresponds to how many units change in an outcome.

#### Table S15. MR-PRESSO: effects of fetal growth on birth outcomes (A) and maternal blood pressure and glucose levels (B)

##### A. Effects of birth weight associated SNPs on birth outcomes

| | | Gestational days | | | Preterm birth | | | Birth weight | | | Birth length | | |
|---|---|---|---|---|---|---|---|---|---|---|---|---|---|
| Birth weight | Method | beta | se | p-val | beta | se | p-val | beta | se | p-val | beta | se | p-val |
| h1 | raw | -0.0058 | 0.0029 | 0.056 | 0.0025 | 0.0007 | **0.00074** | 0.74 | 0.10 | **1.8E-11** | 0.0017 | 0.0005 | **0.0008** |
| | corrected | NA | NA | NA | NA | NA | NA | NA | NA | **5.8E-12** | NA | NA | NA |
| h2 | raw | 0.0010 | 0.0027 | 0.7 | -0.0005 | 0.0007 | 0.44 | -0.17 | 0.11 | 0.11 | -0.0003 | 0.0005 | 0.56 |
| | corrected | NA | NA | NA | NA | NA | NA | NA | NA | **0.012** | NA | NA | NA |
| h3 | raw | -0.0101 | 0.0031 | **0.0012** | 0.0020 | 0.0006 | **0.003** | 1.22 | 0.08 | **6.5E-25** | 0.0029 | 0.0004 | **1.7E-09** |
| | corrected | NA | NA | NA | NA | NA | NA | NA | NA | NA | NA | NA | NA |

##### B. Effects of birth weight associated SNPs on maternal blood pressures and glucose levels

| | | BP | | | SBP | | | DBP | | | FPG | | |
|---|---|---|---|---|---|---|---|---|---|---|---|---|---|
| Birth weigth | Method | beta | se | p-val | beta | se | p-val | beta | se | p-val | beta | se | p-val |
| h1 | raw | -0.0016 | 0.0014 | 0.28 | -0.0029 | 0.0018 | 0.12 | -0.0003 | 0.0015 | 0.85 | -0.0002 | 0.0002 | 0.34 |
| | corrected | NA | NA | NA | NA | NA | NA | NA | NA | NA | NA | NA | 0.63 |
| h2 | raw | 0.0005 | 0.0012 | 0.66 | 0.0006 | 0.0016 | 0.72 | 0.0007 | 0.0012 | 0.57 | -0.0003 | 0.0002 | 0.092 |
| | corrected | NA | NA | NA | NA | NA | NA | NA | NA | NA | NA | NA | 0.19 |
| h3 | raw | 0.0033 | 0.0013 | **0.014** | 0.0043 | 0.0017 | **0.011** | 0.0021 | 0.0013 | 0.11 | -0.0001 | 0.0001 | 0.68 |
| | corrected | NA | NA | NA | NA | NA | NA | NA | NA | NA | NA | NA | NA |

#### Supplementary Figures

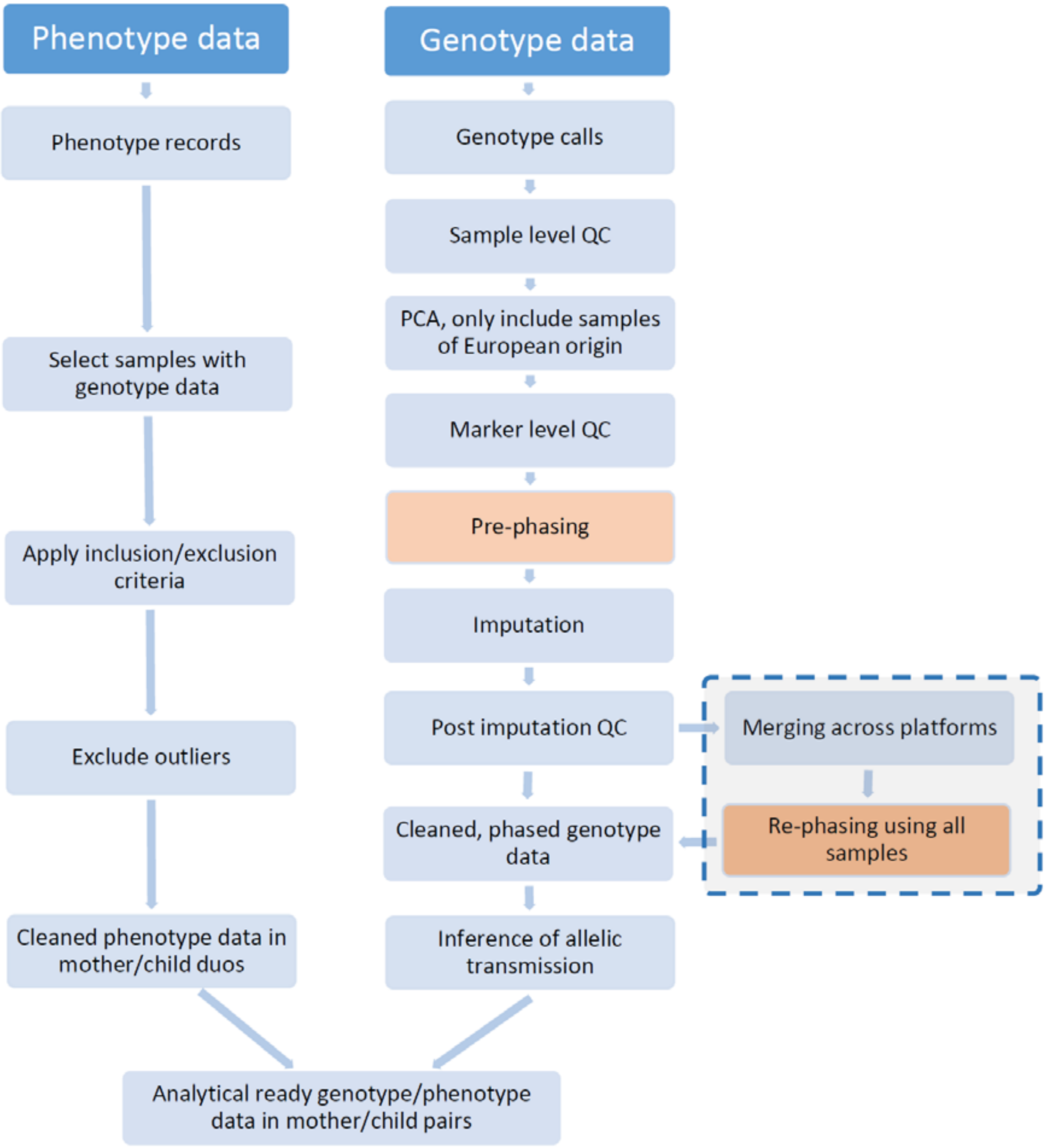

**Figure S1. Processing of phenotype and genotype data**

The processing of phenotype (left) and genotype data (right). Haplotype transmission between mothers and their infants was obtained from the haplotypes inferred from the pre-phasing or re-phasing steps (highlighted in orange). For the FIN data set, a merging and a re-phasing step was added to merge and phase the imputed genotypes of mothers and infants genotyped by different platforms (dashed box).

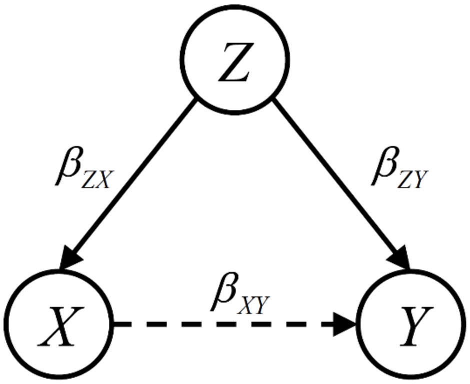

**Figure S2. General representation of confounded association**

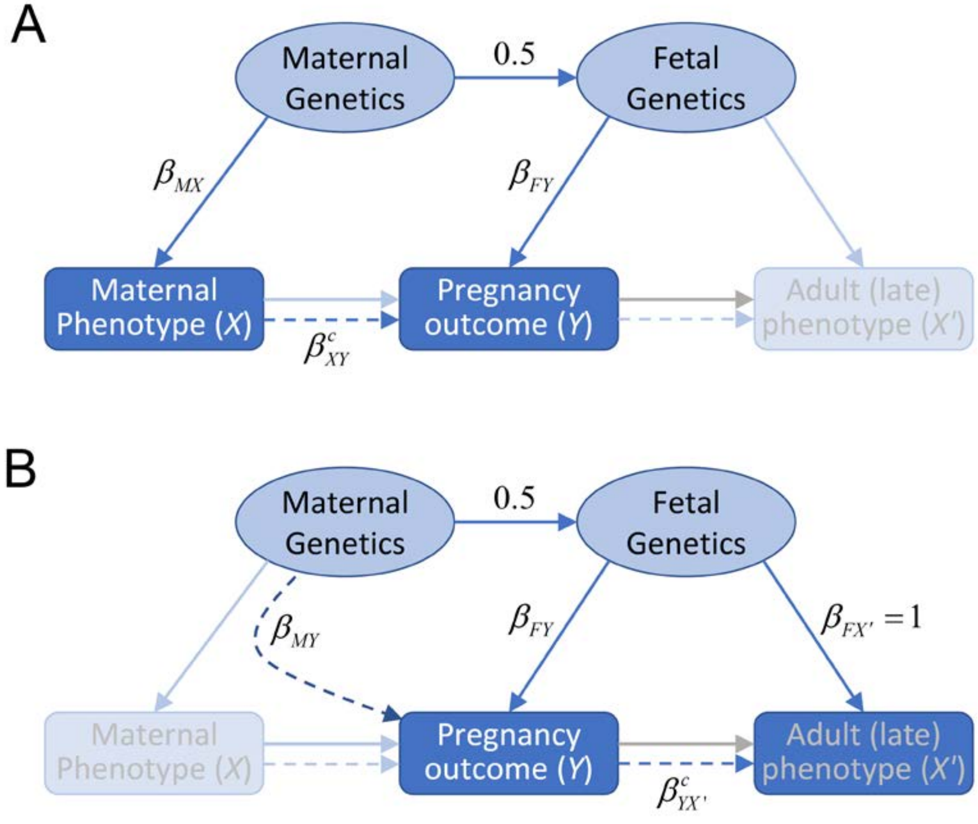

**Figure S3. Genetically confounded association between a maternal phenotype and a birth outcome (A) and between a birth outcome and an adult phenotype in offspring (B) due to shared genetics**

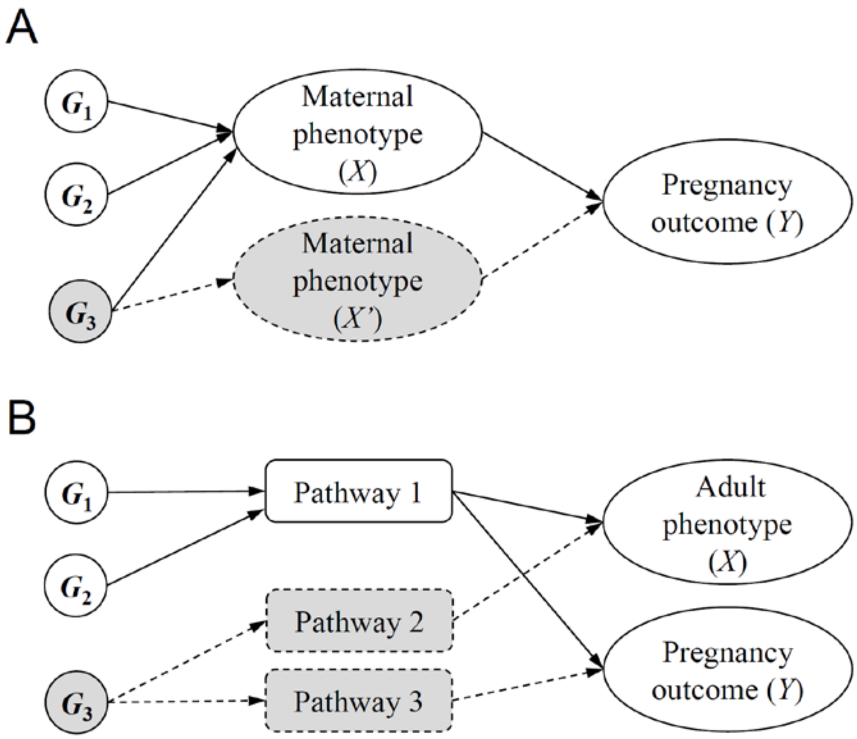

#### Figure S4. Multivariable Mendelian randomization analysis of maternal and fetal genetic effect

**A. Maternal effect.** A list of genetic variants ($G_1$, $G_2$, $G_3$, …) are primarily associated with a maternal phenotype *X*, which causally influences a pregnancy outcome *Y*. In this case, these genetic variants can be used to instrument the maternal causal effect of *X* on *Y* and the estimated effect sizes of these variants on *Y*, especially those based on the non-transmitted alleles (the transmitted alleles can have fetal genetic effect), should be generally proportional to their effects on the maternal phenotype, which can be obtained from existing large GWA studies. Some variants (e.g. $G_3$) may associate with other maternal phenotype (*X'*), which also can influence *Y* (horizontal pleiotropy). Thus, the effects of these variants on *Y* are less correlated with their reported effects on *X*, which could be detected as outliers by MR-PRESSO. The situation of "fetal drive" (a fetal phenotype causally influences a maternal phenotype) is similar, in which case the paternal transmitted alleles of the SNPs associated with a fetal phenotype (e.g. birth weight) could be associated with a maternal phenotype (e.g. maternal blood pressure during pregnancy). **B. Fetal genetic effect.** Genetic variants ($G_1$, $G_2$, $G_3$, …) are associated with both an adult phenotype *X* and a pregnancy outcome *Y*. Some of the variants (e.g. $G_1$ and $G_2$) may be associated with both *X* and *Y* through a common pathway 1. Therefore, the estimated effects of these variants on *Y* (especially based on the paternal transmitted alleles to avoid maternal effect) should be proportional to their effects on *X* because these variants act through the same pathway, although there is no real causal relationship between *X* and *Y*. Some of the variants (e.g. $G_3$) may be associated with *X* and *Y* through distinct pathways (2 and 3) and therefore their effects on *X* and *Y* are relatively independent and could be detected as outliers with horizontal pleiotropy by MR-PRESSO.

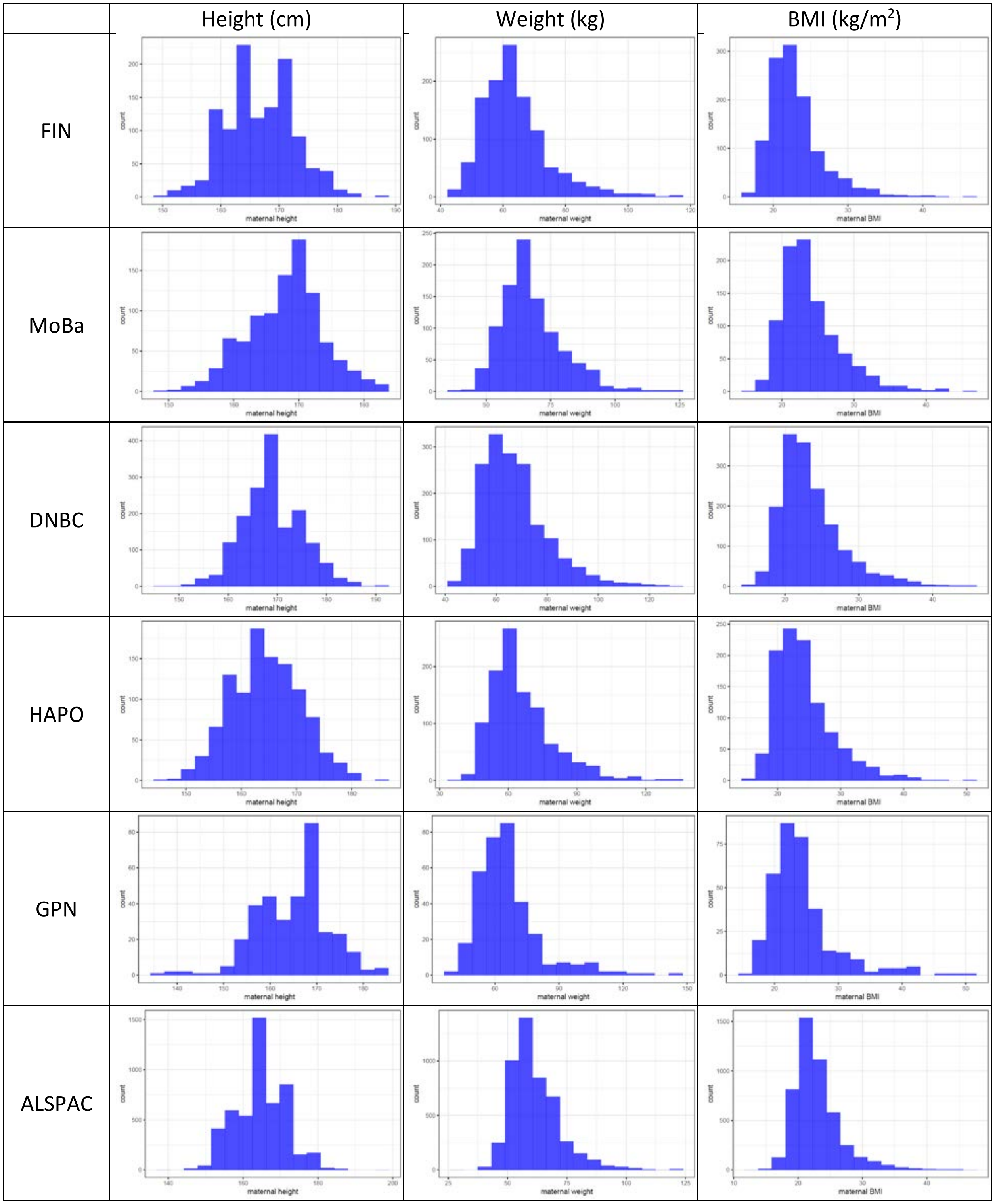

**Figure S5. Distributions of maternal height, weight and BMI**

| | SBP (mmHg) | DBP (mmHg) | FPG* (mmol/L) |
|---|---|---|---|
| HAPO | 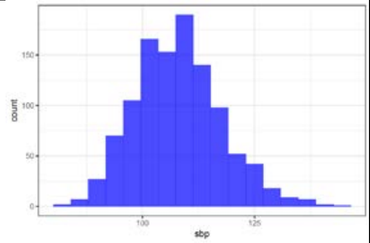 | 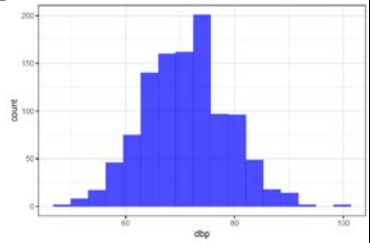 | 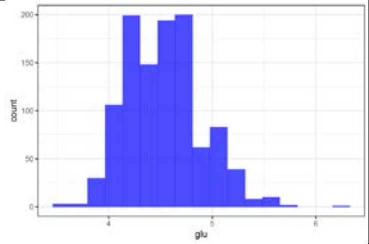 |
| ALSPAC | 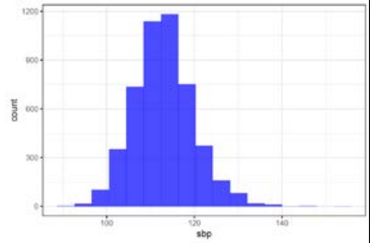 | 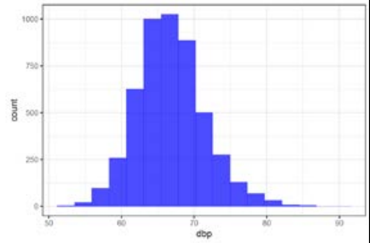 | 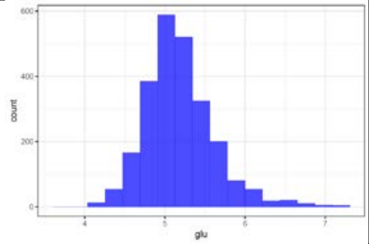 |

#### Figure S6. Distributions of maternal blood pressure and glucose levels

In the HAPO study, fasting plasma glucose (FPG) in the mothers were measured between 24 and 32 weeks gestation. In the ALSPAC study, FPG was measured in a follow-up data collection 18 years after the pregnancy.

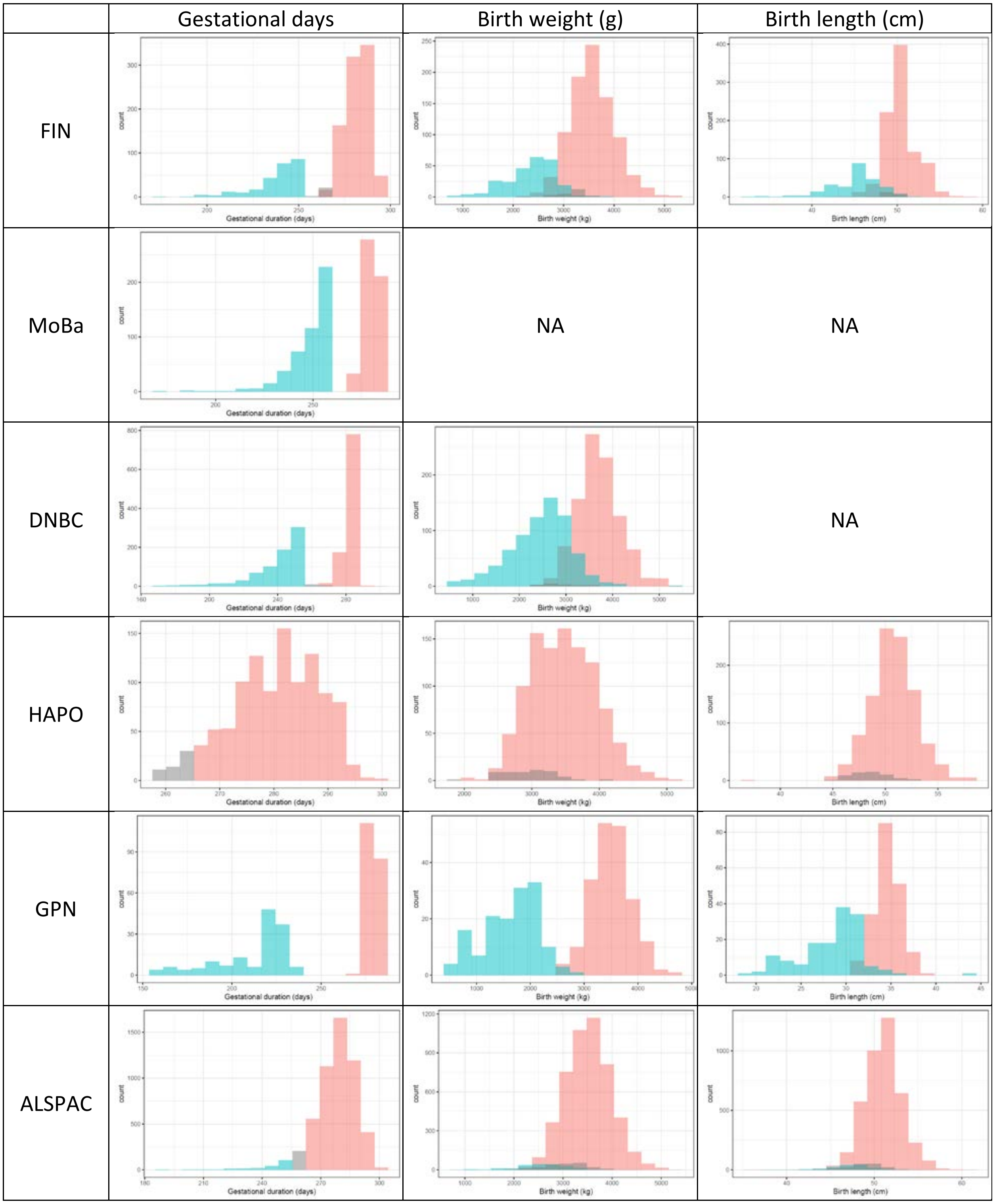

**Figure S7. Distributions of gestational days, birth weight and birth length**

The FIN, MoBa, DNBC, and the GPN were collected for case/control study of preterm birth; therefore, the gestational duration was truncated and enriched for cases (gestational duration < 37 weeks)

<table>
<tr><th>Scores*</th><th>Gestational days</th><th>Preterm log(OR)</th><th>Birth weight (g)</th><th>Birth length (cm)</th></tr>
<tr><td>$S_{h1}$</td>
<td>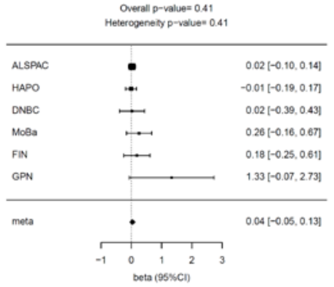
</td>
<td>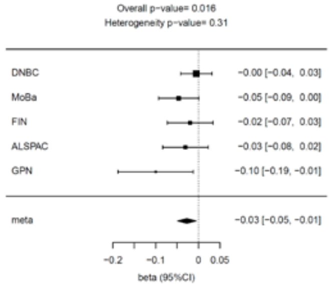
</td>
<td>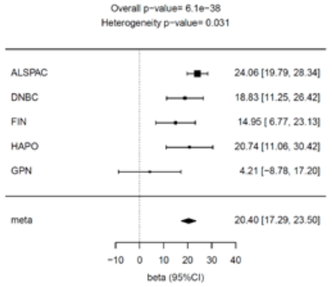
</td>
<td>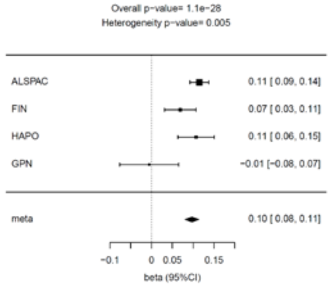
</td></tr>
<tr><td>$S_{h2}$</td>
<td>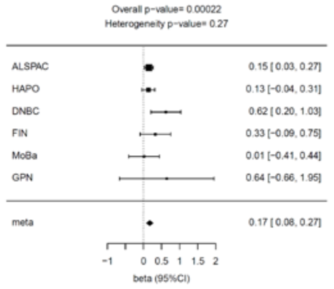
</td>
<td>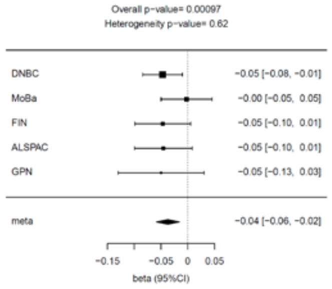
</td>
<td>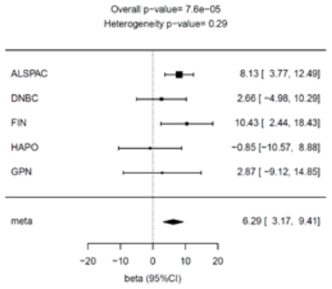
</td>
<td>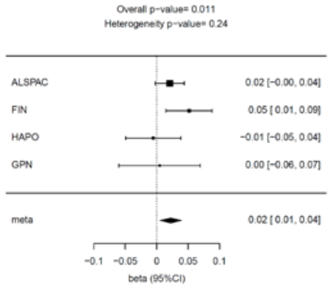
</td></tr>
<tr><td>$S_{h3}$</td>
<td>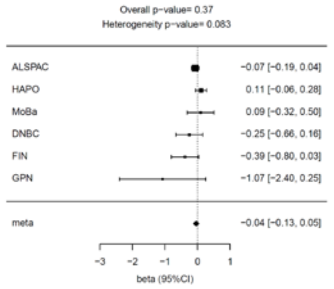
</td>
<td>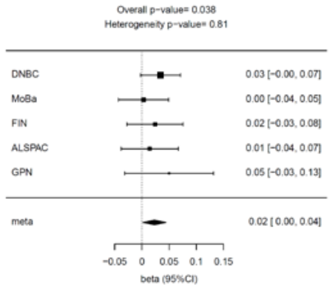
</td>
<td>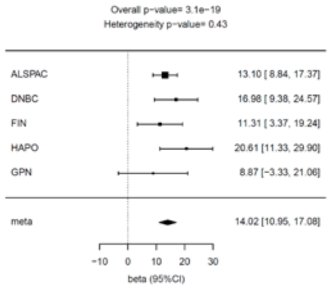
</td>
<td>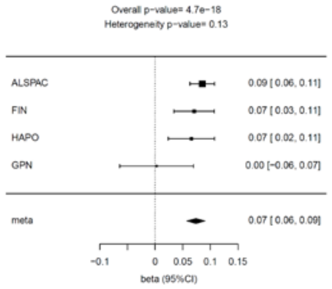
</td></tr>
</table>

**Figure S8. Estimated effect sizes of adult height haplotype scores on pregnancy outcomes**

* $S_{h1}$, $S_{h2}$ and $S_{h3}$ are height haplotype genetic scores built on maternal transmitted, maternal non-transmitted and paternal transmitted alleles.

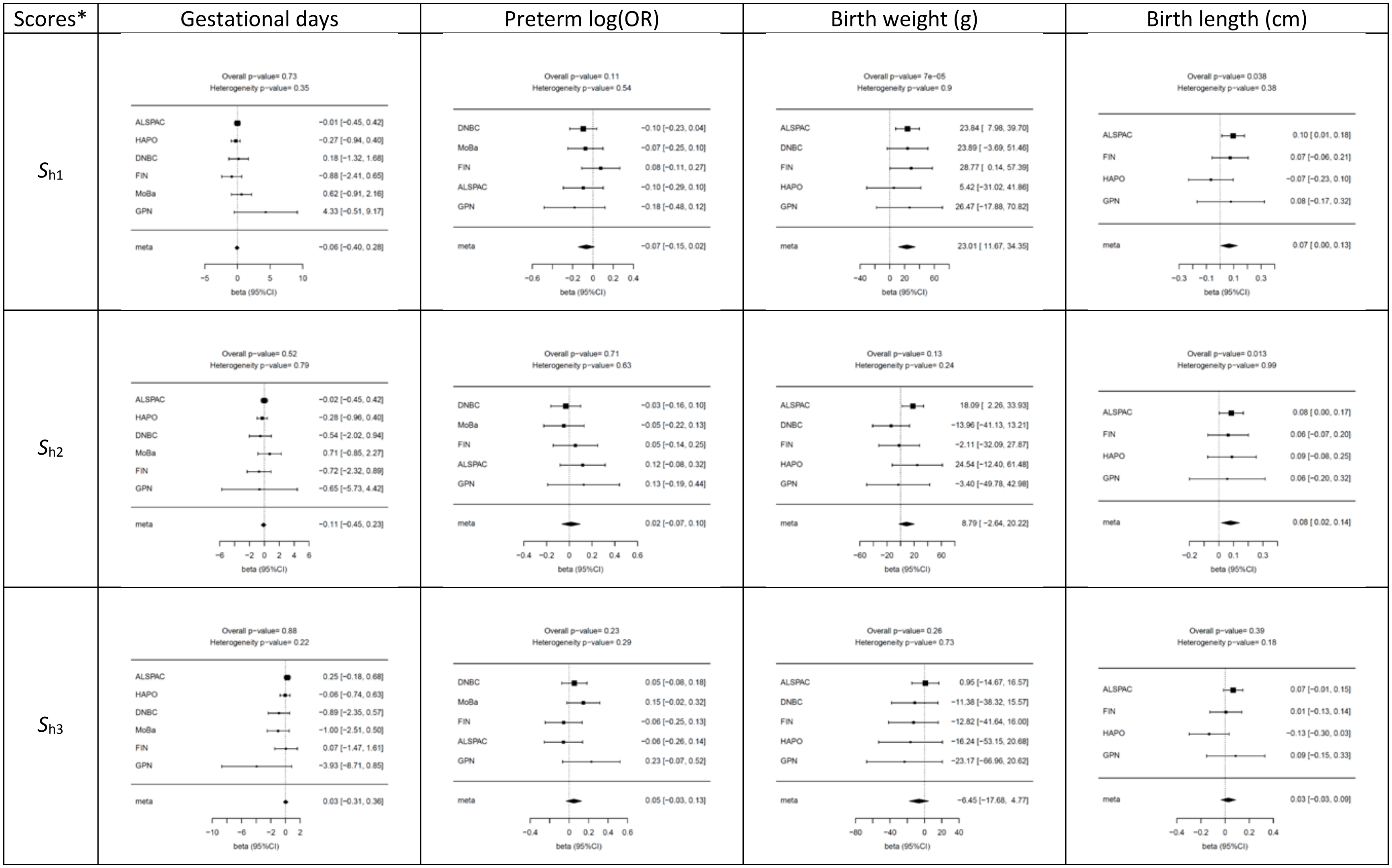

**Figure S9. Estimated effect sizes of BMI haplotype scores on pregnancy outcomes**

* $S_{h1}$, $S_{h2}$ and $S_{h3}$ are the BMI haplotype genetic scores built on maternal transmitted, maternal non-transmitted and paternal transmitted alleles.

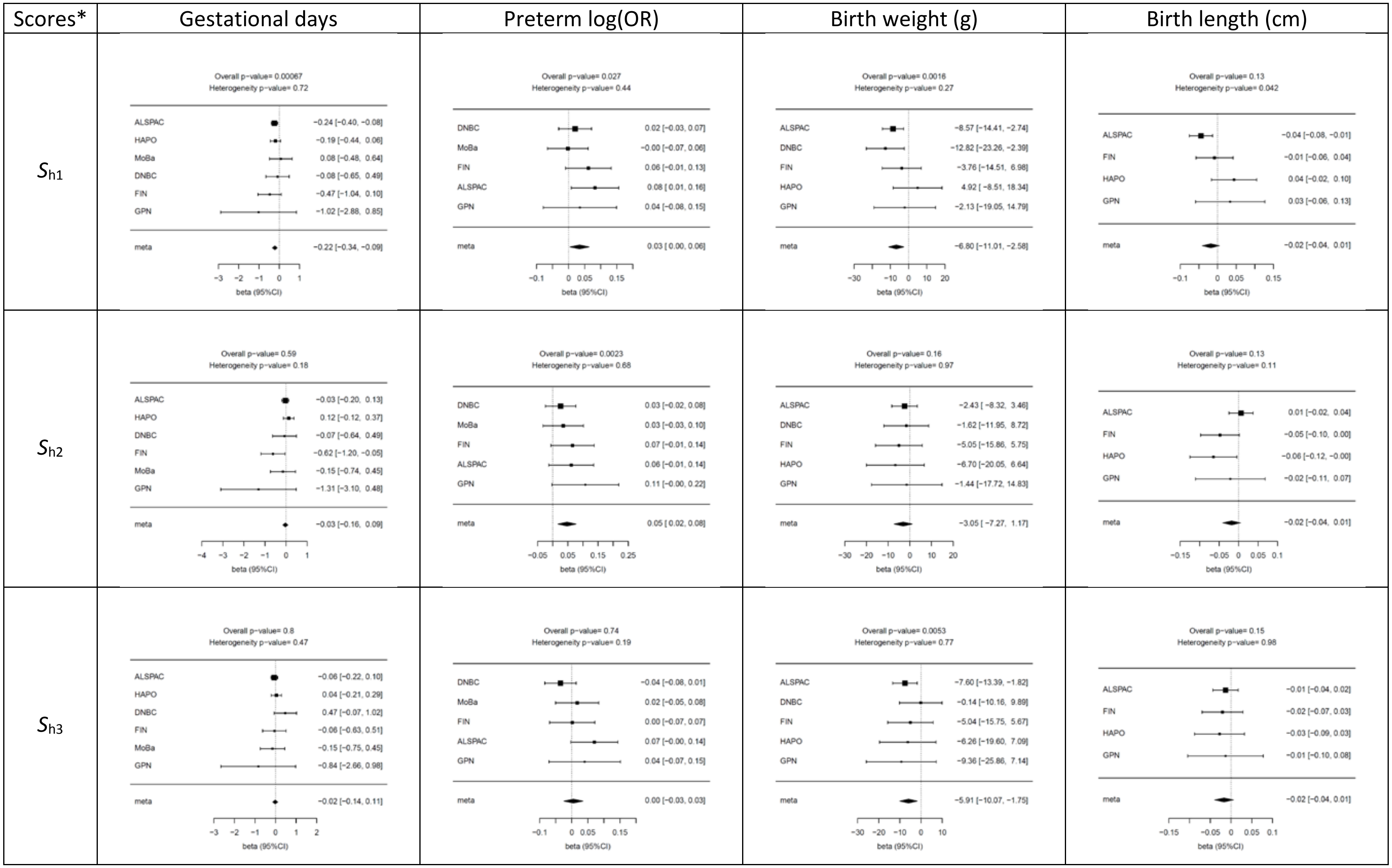

**Figure S10. Estimated effect sizes of blood pressure haplotype scores on pregnancy outcomes**

* $S_{h1}$, $S_{h2}$ and $S_{h3}$ are the blood pressure (BP) haplotype genetic scores built on maternal transmitted, maternal non-transmitted and paternal transmitted alleles.

| Scores* | Gestational days | Preterm log(OR) | Birth weight (g) | Birth length (cm) |
|---|---|---|---|---|
| *S*h1 | 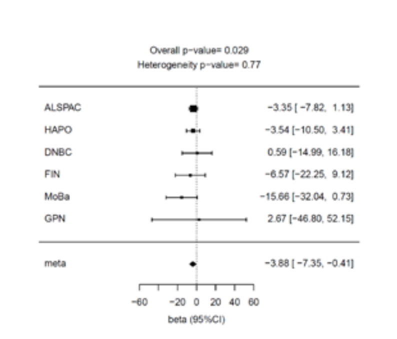 | 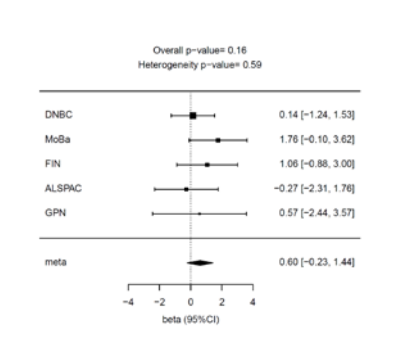 | 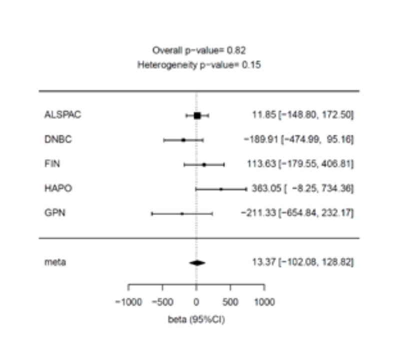 | 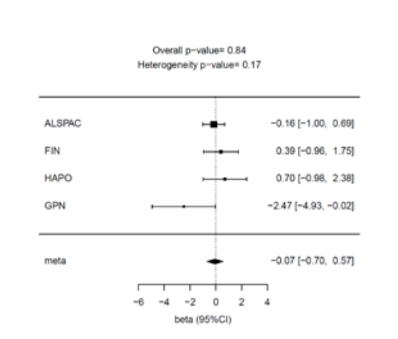 |
| *S*h2 | 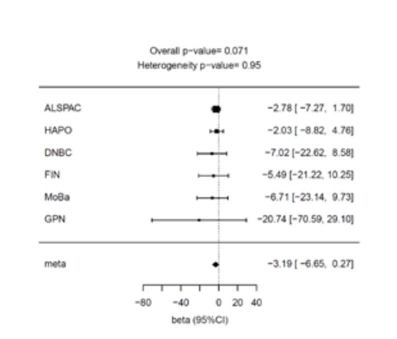 | 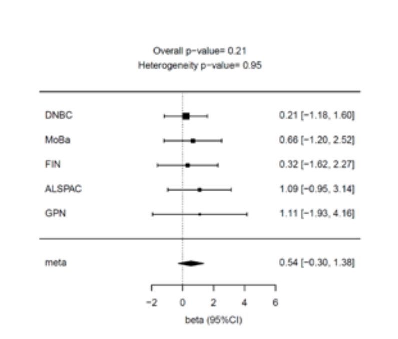 | 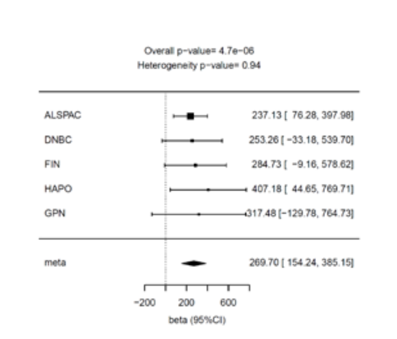 | 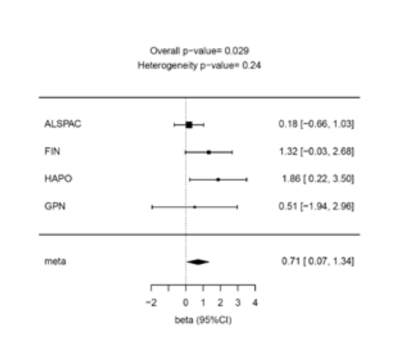 |
| *S*h3 | 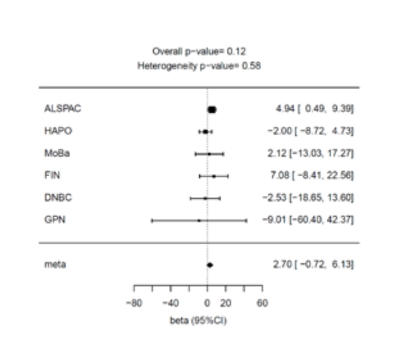 | 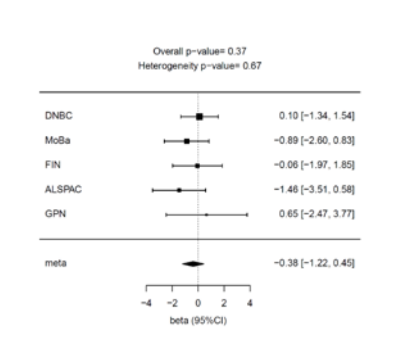 | 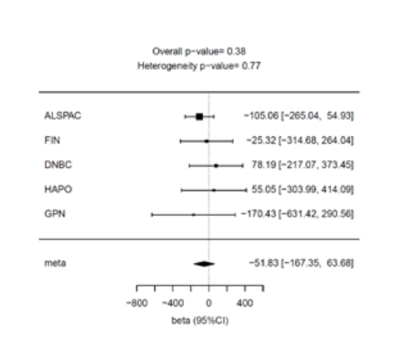 | 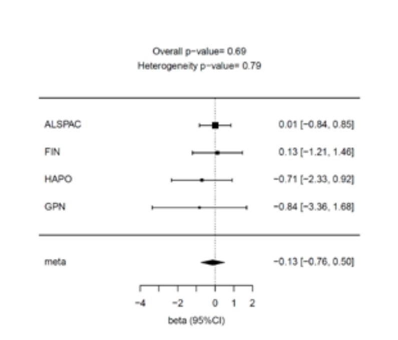 |

**Figure S11. Estimated effect sizes of blood glucose haplotype scores on pregnancy outcomes**

* $S_{h1}$, $S_{h2}$ and $S_{h3}$ are the blood glucose (FPG) haplotype genetic scores built on maternal transmitted, maternal non-transmitted and paternal transmitted alleles.

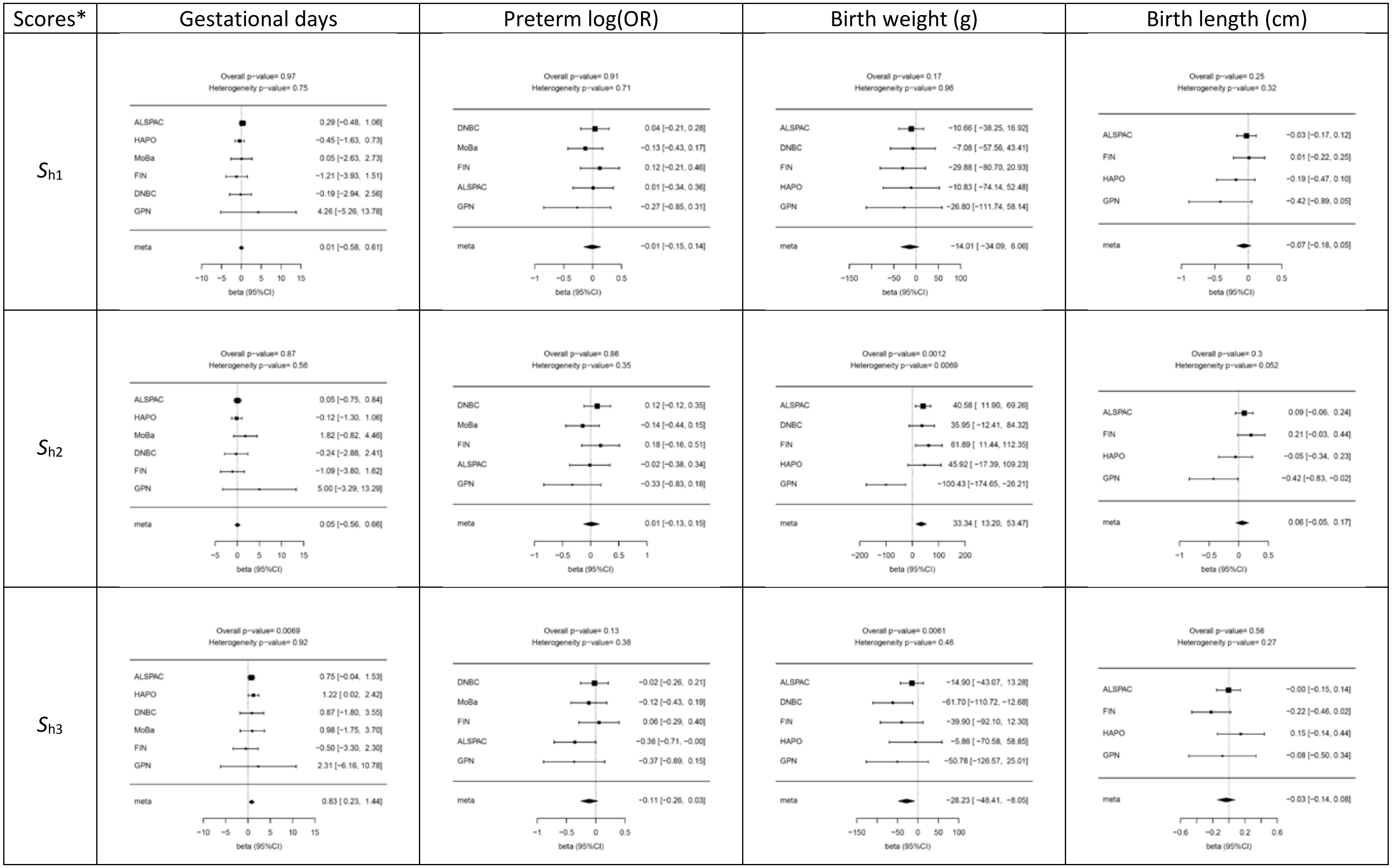

**Figure S12. Estimated effect sizes of type 2 diabetes haplotype scores on pregnancy outcomes**

* $S_{h1}$, $S_{h2}$ and $S_{h3}$ are the type 2 diabetes (T2D) (with SNPs primarily associated with blood glucose levels removed) haplotype genetic scores built on maternal transmitted, maternal non-transmitted and paternal transmitted alleles.

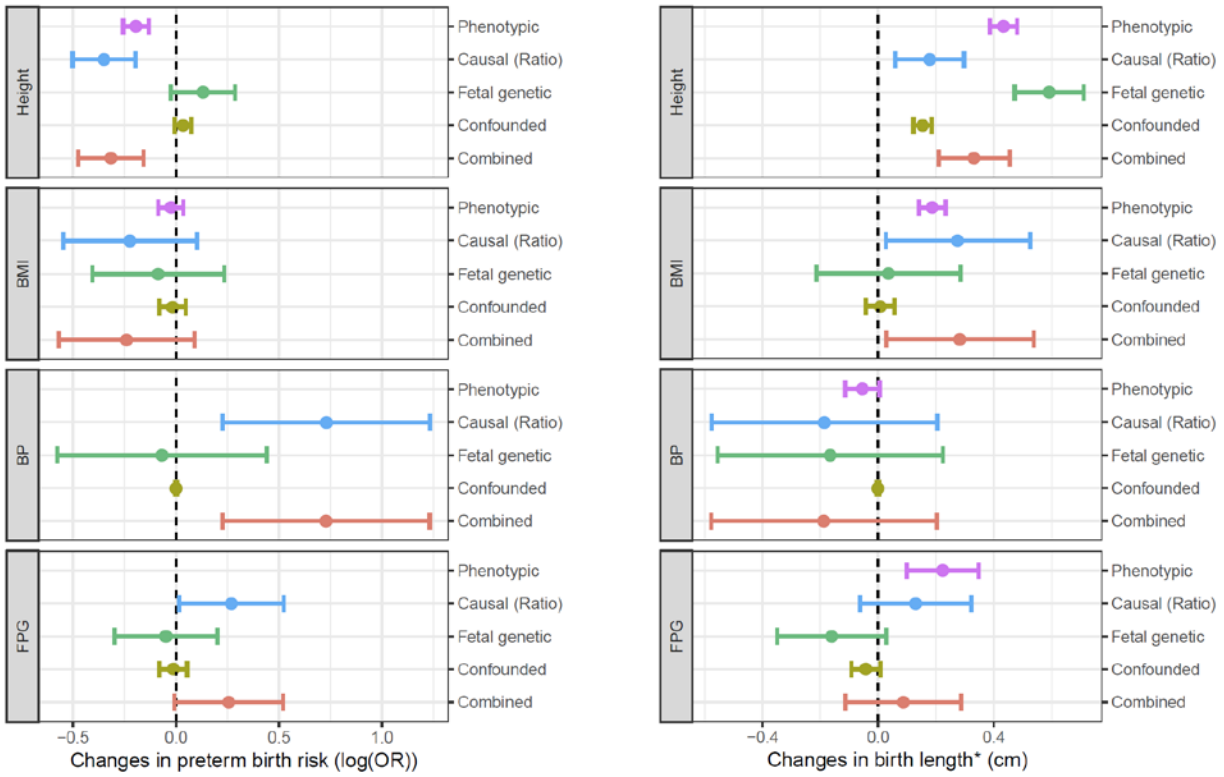

**Figure S13. Estimated sizes of causal effects and genetically confounded associations per 1SD changes in maternal traits on preterm birth risk and birth length**

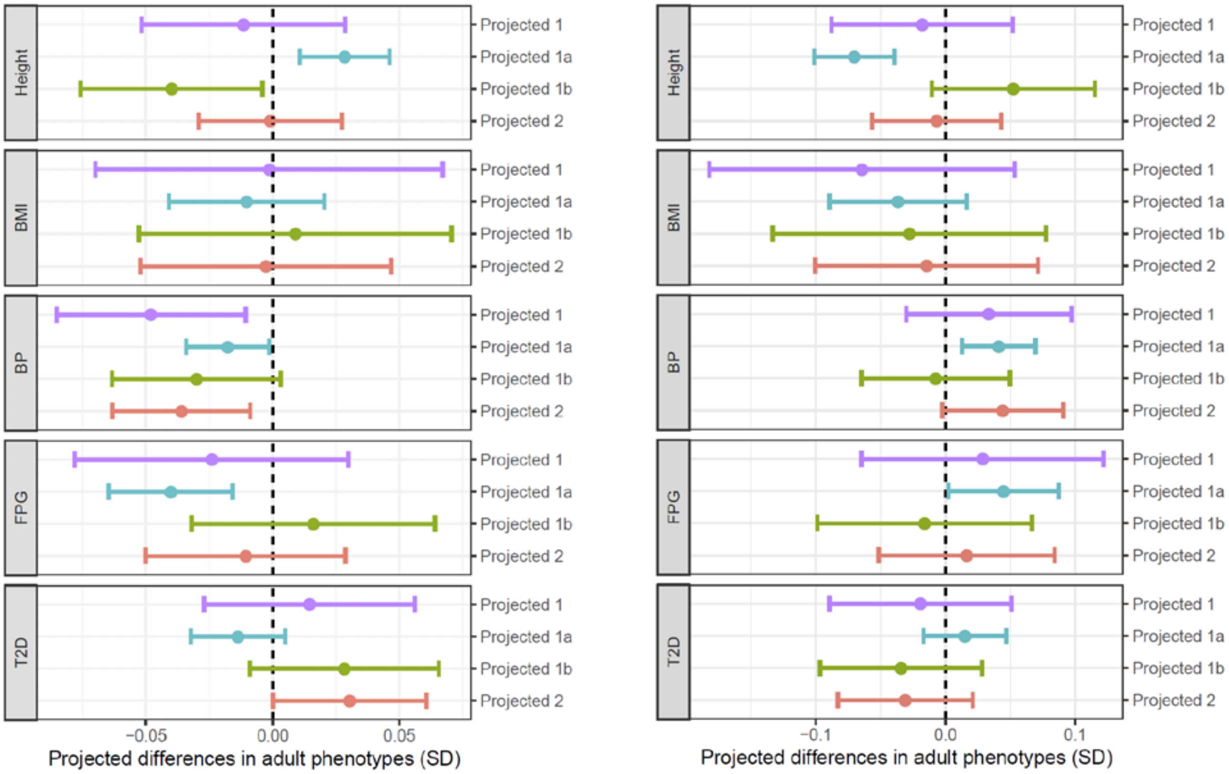

**Figure S14. Estimated differences in adult phenotypes (SD) per 1SD difference in gestational days (left) and preterm birth risk(right)**

The genetically confounded association was estimated by two methods (Supplementary methods):

$$\hat{\beta}^{c}_{YX'} = h^2 \frac{\mathrm{Var}(X')}{\mathrm{Var}(Y)}\left(\frac{\hat{\beta}_{MY}}{2} + \hat{\beta}_{FY}\right) \text{ (Method 1) and } \hat{\beta}^{c}_{YX'} = h^2 \frac{\mathrm{Var}(X')}{\mathrm{Var}(Y)}\left(\frac{\widehat{\beta_{\mathrm{h1}}+\beta_{\mathrm{h3}}}}{2}\right) \text{ (Method 2)}$$

Method 1a and Method 1b show the the maternal ($\frac{\beta_{MY}}{2}$) and the fetal component ($\beta_{FY}$) of the estimated genetically confounded associations.

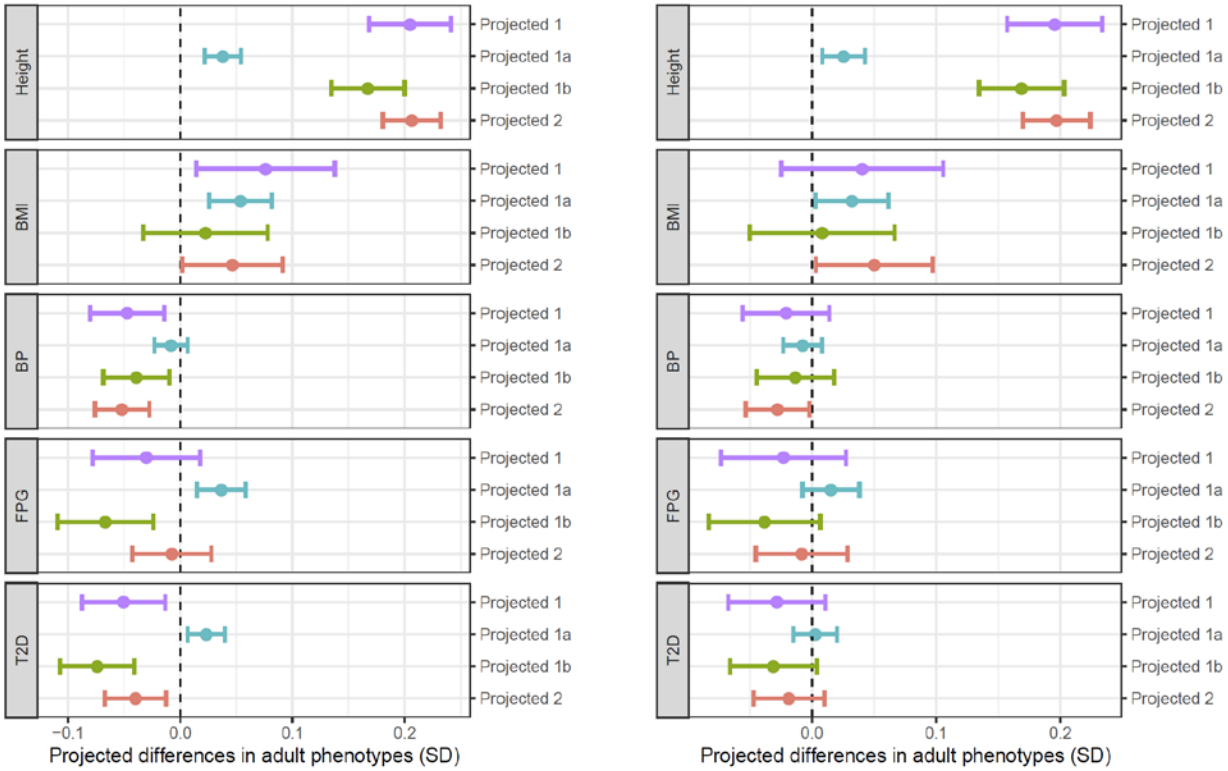

**Figure S15. Estimated differences in adult phenotypes (SD) per 1SD difference in gestational age adjusted birth weight (left) and birth length (right)**

The genetically confounded association was estimated by two methods (Supplementary methods):

$$\hat{\beta}^{c}_{YX'} = h^2 \frac{\mathrm{Var}(X')}{\mathrm{Var}(Y)}\left(\frac{\hat{\beta}_{MY}}{2} + \hat{\beta}_{FY}\right) \text{ (Method 1) and } \hat{\beta}^{c}_{YX'} = h^2 \frac{\mathrm{Var}(X')}{\mathrm{Var}(Y)}\left(\frac{\widehat{\beta_{h1}+\beta_{h3}}}{2}\right) \text{ (Method 2)}$$

Method 1a and Method 1b show the the maternal ($\frac{\beta_{MY}}{2}$) and the fetal component ($\beta_{FY}$) of the estimated genetically confounded associations.

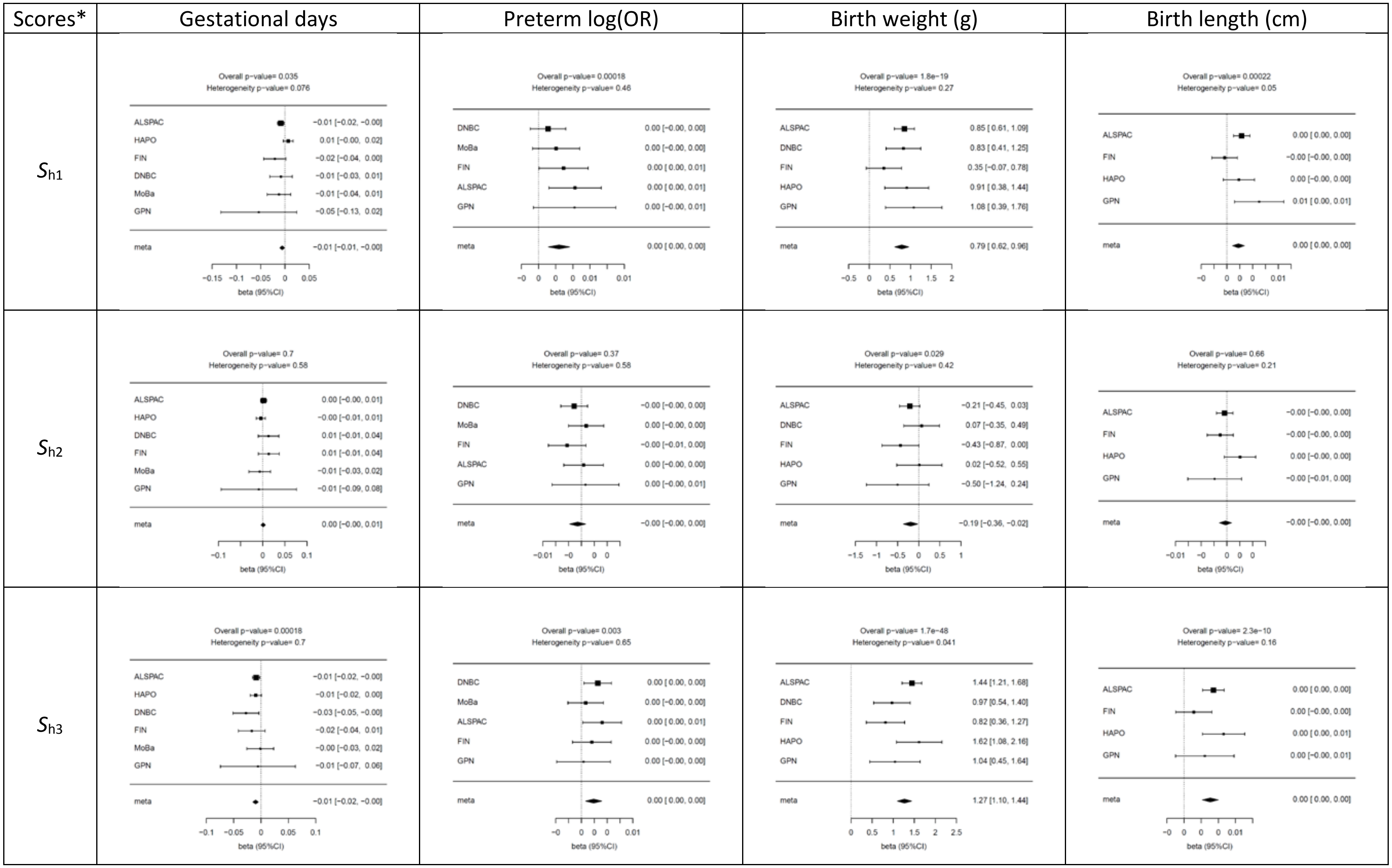

**Figure S16. Estimated effect sizes of birth-weight haplotype scores on pregnancy outcomes**

* $S_{h1}$, $S_{h2}$ and $S_{h3}$ are the birth weight haplotype genetic scores built on maternal transmitted, maternal non-transmitted and paternal transmitted alleles.

| Scores* | BP (mmHg) | SBP (mmHg) | DBP (mmHg) | FPG (mmol/L) |
|---|---|---|---|---|
| $S_{h1}$ | 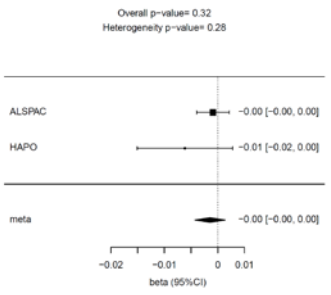 | 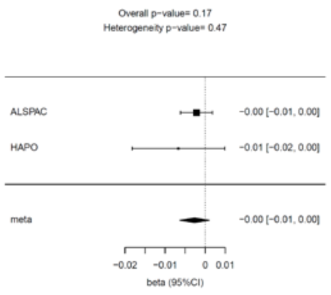 | 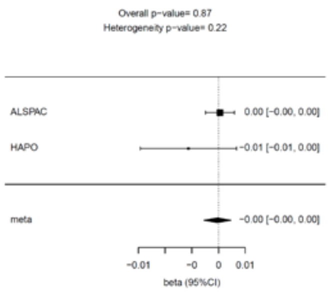 | NA |
| $S_{h2}$ | 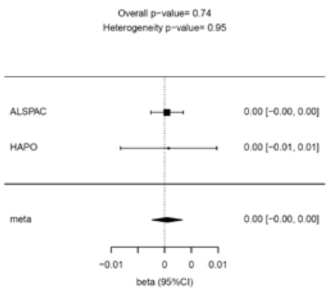 | 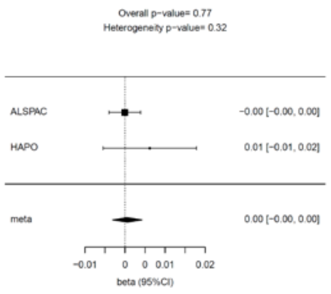 | 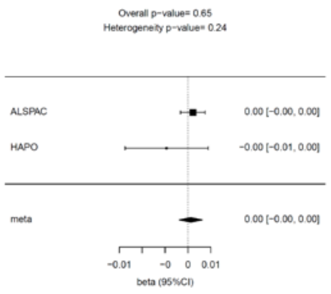 | NA |
| $S_{h3}$ | 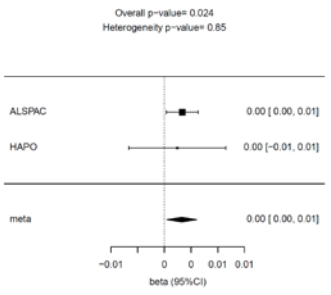 | 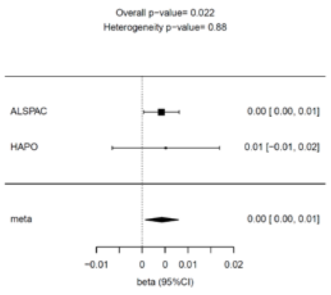 | 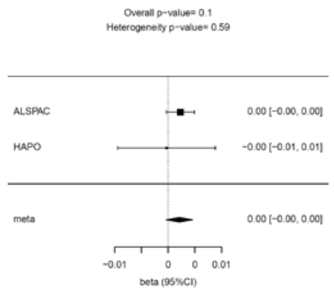 | NA |

**Figure S17. Estimated effect sizes of birth-weight haplotype scores on maternal blood pressure and glucose levels**

* $S_{h1}$, $S_{h2}$ and $S_{h3}$ are the birth weight haplotype genetic scores built on maternal transmitted, maternal non-transmitted and paternal transmitted alleles.

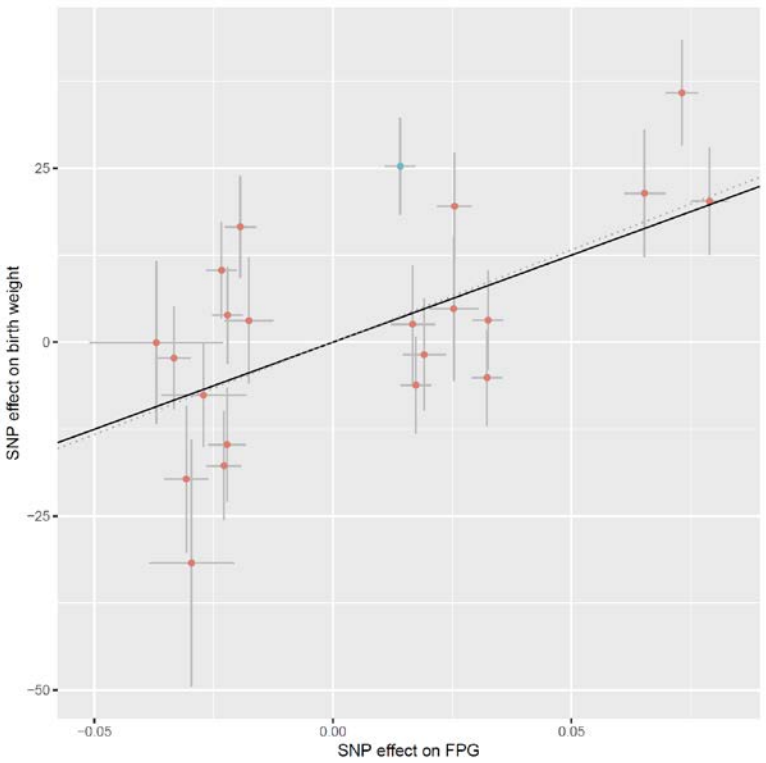

#### Figure S18. An example of multivariable MR estimate

Multivariable MR estimate of causal effect of maternal FPG on birth weight using maternal non-transmitted alleles. Each point represents the per allele association of the 22 glucose associated SNPs – x-axis: effect sizes on FPG level (and standard error) reported by the reference GWA study; y-axis: effect sizes on birth weight estimated from the maternal non-transmitted alleles (h2) from this study. The line shows the multivariable MR estimate. Removal of the outlier SNP (blue) did not change the causal estimate (the regression line).
